# Mechanism of degrader-targeted protein ubiquitinability

**DOI:** 10.1101/2024.02.05.578957

**Authors:** Charlotte Crowe, Mark A. Nakasone, Sarah Chandler, Michael H. Tatham, Nikolai Makukhin, Ronald T. Hay, Alessio Ciulli

## Abstract

Small molecule degraders of disease-driving proteins offer a clinically proven modality with enhanced therapeutic efficacy and the potential to tackle previously undrugged targets. Thermodynamically stable and kinetically long-lived degrader-mediated ternary complexes can drive faster, more profound and durable target degradation, however the mechanistic features by which they impact on target ubiquitination remain elusive. Here, we solve cryo-EM structures of the VHL Cullin 2 RING E3 ligase complexed with degrader MZ1, target protein Brd4^BD2^ and primed for catalysis with its cognate E2-ubiquitin bound. We find that Brd4^BD2^ adopts a favourable orientation towards the E2 active site. In vitro ubiquitination coupled with mass spectrometry illuminates a patch of ubiquitinable lysines on one face of Brd4^BD2^, with Lys456 showing optimal distance and geometry for nucleophilic attack. Our results demonstrate the proficiency of MZ1 in directing the substrate towards catalysis, explains the favourability of Brd4^BD2^ for ubiquitination, and reveals the flexibility of the enzyme in capturing sub-optimal lysines. We propose a model for ubiquitinability of degrader-recruited targets that provides a mechanistic blueprint for further rational drug design and optimization.

**One-Sentence Summary:** Structural assembly a PROTAC-mediated complex of whole Cullin RING E3 ligase with bound target and E2-ubiquitin reveals structural and mechanistic insights of specificity for target protein ubiquitination.

## Main Text

Targeted protein degradation has emerged as a powerful new modality of chemical biology and therapeutic intervention against proteins that drive disease (*1*). The approach most developed to date involve the design or discovery of small molecules, so-called degraders, that harness the activity of the endogenous ubiquitin-proteasome system to induce ubiquitination and subsequent degradation of target proteins (*2*). This is most effectively achieved by recruiting a *neo*-substrate (i.e. a protein not normally processed as a native substrate) to the key enzymatic machineries that catalyse protein ubiquitination, namely the ubiquitin E3 ligases, and predominantly the Cullin-RING ligases (CRLs) (*3-5*). Degraders are typically categorized as either PROteolysis Targeting Chimeras (PROTACs), i.e. bifunctional molecules composed of a binder to the E3 ligase and a binder to the protein target, joined by a linker unit (*6*); or monovalent for binding to either E3 ligase or target, referred to as molecular glues (*7*). PROTACs, since their development and validation as active in cells and *in vivo* (*8-10*), have emerged as a rapidly growing area of research in both academic research and drug discovery, as illustrated by the >25 drug candidates that are currently in clinical trials for various diseases (*6, 11*). The vast majority of PROTAC degraders co-opt the activity of one of two CRLs: the von Hippel-Lindau Cullin 2 ligase complex (CRL2^VHL^) (*12-14*) and the cereblon Cullin 4 ligase complex (CRL4^CRBN^) (*15-17*), which is also the target of molecular glue degraders thalidomide and lenalidomide amongst others (*18, 19*). The repertoire of hijackable E3 ligases has recently expanded as small-molecule ligands are being developed for more Cullin RING E3s and used in PROTACs (*20*), e.g. DCAF1 (*21*) and KLHDC2 (*22*), and new E3s are being identified as co-opted by molecular glue degraders, including DCAF15 (*23*), DCAF16 (*24*) and the CRL4 adaptor DDB1 (*25*), amongst others. Despite their chemical distinction, PROTACs and molecular glues converge on the same mechanism of action that requires the formation of the ternary complex E3:degrader:target as the key species driving productive target ubiquitination and subsequent proteasomal degradation (*13, 26-28*). However, most structural, biophysical and mechanistic studies on PROTAC ternary complexes disclosed to date have been restricted to the substrate receptor/adaptor components of the CRL (*13, 17, 29*), thus lacking the fully assembled catalytically-active CRL complex (*30*). Furthermore, mechanistic and structural investigations of PROTAC-mediated ubiquitination have remained sparse or limited in resolution (*31-33*). To fully enable and guide the design and optimization of degrader drugs, there is a growing need to understand how degraders recruit the whole native catalytic enzymatic machinery to illuminate their mechanism of action.

Structural and mechanistic features of degrader-mediated ternary complexes directly impact on the pharmacological activity of degraders and thus represent an important optimization species for rational structure-based drug design (*26*). Multiple studies have revealed that thermodynamically cooperative, stable and kinetically long-lived degrader-mediated ternary complexes, in most cases, underpin efficient and selective protein degradation profiles of degraders (*13, 29, 34-39*). Yet how ternary complex formation influence productive and selective *neo*-substrate ubiquitination within the PROTAC-co-opted catalytic mechanism of the multiprotein CRL complex has to date remained largely elusive. To this end, we decided to investigate the structure and mechanisms of the PROTAC MZ1 (**Figure 1A**), our potent and fast VHL-recruiting BET degrader, which exhibits preferential degradation of Brd4 over other BET proteins (*8, 29, 40*). MZ1 is a well-characterized and widely-used PROTAC degrader whose binding and ternary complex formation has been probed by varioous biophysical and structural methods by us and others since our first PROTAC co-crystal structure published in 2017 (*13, 29, 39, 41-43*). These studies have revealed that MZ1 recruits all BET bromodomains to VHL with positive cooperativity, yet despite binding all BET-BDs with comparable binding affinity at binary level, it preferentially recruits the BD2s over the BD1s in ternary complexes. Of note, MZ1 forms the most cooperative, stable and long-lived ternary complex with Brd4^BD2^, even over its close homologue Brd3^BD2^, explaining its preferential Brd4 degradation selectivity (*13, 29*). Indeed, Brd4^BD2^ is the BET bromodomain isoform that has been co-crystallized by far the most to date, not only with MZ1 but also with other VHL-based PROTACs including our structure-based designed macroPROTAC-1 (*41*) and AT7 (*44*), and other analogues (*45*). In all these co-crystal structures, the orientation of the bromodomain relative to VHL within the ternary complex is very similar, highlighting a highly conserved substrate binding mode.

**Figure 1.**
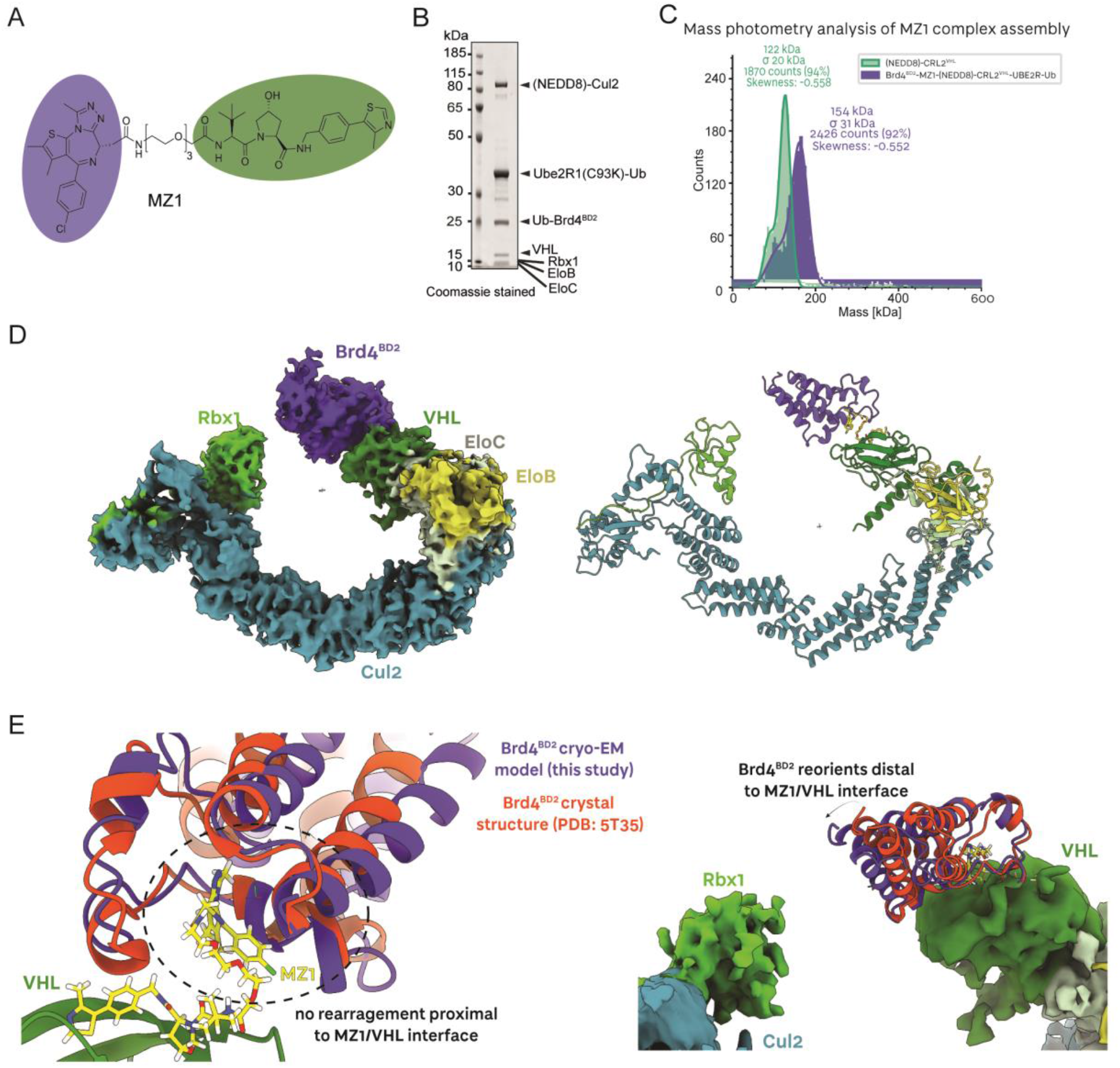
MZ1 orients the Brd4 bromodomain 2 neo-substrate towards Rbx1. (**A**) Chemical structure of the bivalent PROTAC MZ1 (in purple: the Brd4^BD2^ ligand; in green: the VHL ligand). (**B**) SDS-PAGE of the assembled Brd4^BD2^-MZ1- (NEDD8)-CRL2^VHL^-UBE2R1(C93K)-Ub complex applied to cryo-EM grids. (**C**) Mass photometry histogram of the fully assembled Brd4^BD2^-MZ1-(NEDD8)-CRL2^VHL^-UBE2R1(C93K)-Ub complex (purple) applied to cryo-EM grids, versus (NEDD8)-CRL2^VHL^ (green). The peaks have been fitted with Gaussian curves. (**D**) The ∼4.0 Å resolution cryo-EM reconstruction of Brd4^BD2^-MZ1-(NEDD8)-CRL2^VHL^. The E3 ligase NEDD8-CRL2^VHL^ (green) recruits MZ1 (yellow) which in turn binds the neo-substrate Brd4^BD2^ (purple). To the right, a ribbon representation of the complex modelled into the displayed cryo-EM volume (same colour scheme). (**E**) The cryo-EM model (this study) of the VHL-MZ1-Brd4^BD2^ interface overlaid with the same interface from the crystal structure by Gadd *et al*. (PDB: 5T35) (*13*). Brd4^BD2^ displays a conserved interface proximal to the MZ1/VHL binding site, and a re-orientation distal to the MZ1/VHL binding site, towards the E3 ligase RING box domain.

To gain a deeper understanding of degraders’ mechanism of action, we determined structures of multi-subunit catalytic complexes, composed of the full CRL2^VHL^, an E2 conjugating enzyme loaded with ubiquitin, and Brd4^BD2^ as the “*neo*-substrate”, all assembled together by MZ1 as representative PROTAC degrader. CRL2^VHL^ is a 150 kDa E3 ligase composed of five subunits: the substrate receptor von Hippel-Lindau (short isoform VHL19, 18 kDa or long isoform VHL30, 24 kDa), the adaptor proteins Elongin C (EloC, 12 kDa) and Elongin B (EloB, 15 kDa), the Cullin 2 scaffold (Cul2, 87 kDa), and the RING box protein 1 (Rbx1, 12 kDa). VHL-EloC-EloB (VCB) is recruited to the N-terminus of Cul2 at the interface between VHL and EloC (*12, 46*). At the C-terminal domain of cullin 2, the E1 NEDD8-activating enzyme APPBP1-UBA3 and the E2 NEDD8-conjugating enzyme UBE2M work with the NEDD8 E3 ligase Rbx1 to neddylate Cul2, i.e. to covalently modifying the K689 residue of Cul2 WHB domain with NEDD8 (*47, 48*). This modification has been shown to induce conformational rearrangements of cullin-RING complexes as well as facilitate engagement of E2 ubiquitin-conjugating enzymes (*49-51*). We therefore set out to determine the structures of fully active NEDD8-CRL2^VHL^ with Brd4^BD2^ using single particle cryogenic electron microscopy (cryo-EM). The initial focus was on the fully assembled Brd4^BD2^-MZ1-(NEDD8)-CRL2^VHL^-E2-Ub complex. We expressed and purified each CRL2^VHL^-complex components, namely VCB and Cul2-Rbx1 separately, then *in vitro* neddylated the assembled VCB-Cul2-Rbx1 (CRL2^VHL^) complex, purified it by size-exclusion chromatography, and showed our NEDD8-CRL2^VHL^ was catalytically competent (**Fig. S1**). As for the E2 conjugating enzyme, we leveraged our previously reported strategy of engineering an active site Cys-to-Lys mutant suitable for loading donor-ubiquitin *via* a stable isopeptide bond (*52*), and designed, assembled and purified *in vitro* a mono donor-ubiquitin adduct for the poly-ubiquitinating E2 enzyme Ube2R1 (also known as Cdc34 (*53*)), referred herein as Ube2R1(C93K)-Ub (**Fig. S2**). We next reconstituted a full Brd4^BD2^-MZ1-(NEDD8)-CRL2^VHL^-Ube2R1(C93K)-Ub complex, and evidenced the mono-dispersed nature of its full-assembly using mass photometry as deemed suitable for cryo-EM grid sample preparation (**Figure 1B,C**). Given the complex preference to ‘lie flat’ in the cryogenic sample, the reconstructed cryo-EM volume presented anisotropic artifacts, nonetheless the map was of sufficient quality to build atomic models.

Our ∼4.0 Å resolution cryo-EM structure presents a first opportunity to appreciate the overall conformation of the *neo*-substrate Brd4^BD2^ in complex with a PROTAC and full-length active E3 ligase (**Figure 1D**, see **Fig. S3** for cryo-EM data analysis). We could confidently build atomic models corresponding to Brd4^BD2^-MZ1-CRL2^VHL^ within the density. Despite the E2-Ub conjugate being present in this sample, data processing showed a much lower population of substrate-PROTAC-E3-E2-Ub compared to just substrate-PROTAC-E3 (**Fig. S3**), likely reflecting the rapid dynamics of assembly and disassembly of the E3-E2 complex (*54, 55*). Distinct crystal structures and cryo-EM structures of non-neddylated Cul2-Rbx1 had previously revealed that the WHB domain of Cul2 and Rbx1 are closely packed (*30, 56, 57*) (**Fig. S4**). Cul2 neddylation has been shown to abolish these contacts, allowing Rbx1 to be ‘freed’ and recruit E2∼Ub (*30*). This was indeed observed in our structure, where volume corresponding to Rbx1 is observed, albeit at lower resolution relative to the rest of the structure due to its mobility, pointing towards the substrate (**Fig. S4**). We did not observe density for NEDD8 or the WHB domain of Cul2, also consistent with the highly dynamic neddylation state of the CRL2 (**Figure 1D**).

Our prior models of full-length CRL2^VHL^ based on structural alignments and superpositions had suggested that the substrate-binding interface of VHL is located ∼45 Å from the Rbx1 domain (**Fig. S5A**) (*12, 46*). In our cryo-EM structure, the *neo*-substrate Brd4^BD2^ is recruited to the substrate receptor VHL in the same overall orientation as observed in our previous co-crystal structure (*13*), with no significant rearrangement at the VHL-Brd4 protein-protein interface as “glued” by MZ1, consistent with the tight neo-interactions mediated by MZ1 within the ternary complex (**Figure 1E, left**). Nonetheless, in our cryo-EM consensus volume we observed a slight ‘tilt’ of the bromodomain towards Rbx1 (**Figure 1E, right**), bringing the gap between substrate receptor and the Rbx domain down to ∼10 Å (**Fig. S5B**). The ‘open ring’-like structure suggested that an E2-donorUb conjugate could potentially approach substrate lysines or an acceptor ubiquitin within this ∼10 Å gap. Of note, while this shortened gap is consistent with what is observed with other PROTAC-bound cocrystal structure of VHL with Brd4^BD2^ (**Fig. S5C-D**), other ternary crystal structures with Brd4^BD1^ show the bromodomain pointing in a different direction, away from Rbx1 (**Fig. S5F-H**). Together, our full-length CRL2^VHL^-MZ1-Brd4^BD2^ tertiary structure validates the crystallographic pose of the *neo*-substrate relative to VHL and reveals subtle flexibility within the system allowing bridging of the gap between the substrate-binding and the catalytic sites of the E3 ligase towards ubiquitination.

We next aimed to identify which lysine residues on the Brd4 bromodomain are most accessible to and targeted for ubiquitination. *In vitro* ubiquitination products with both UBE2R1 and UBE2D2 (also known as UbcH5b, which is thought to preferentially prime a substrate with a first ubiquitin (*58*)) were resolved by SDS-PAGE, bands of Brd4-ubiquitinated products excised, digested with trypsin, and K-GG-modified peptides identified by mass spectrometry (**Figure 2A** and **Fig. S6**). We consistently identified eight ubiquitination sites on Brd4^BD2^ (K333, K346, K249, K355, K362, K368, K445 and K456), each from at least one peptide identified in the MS with Andromeda score greater than 100 (see Methods). This overall cluster of ubiquitination sites is seen at 3 h both in the excised UBE2R1-catalysed octa-Ub-Brd4^BD2^ product (79 kDa, **Figure 2A** and **Fig. S7**), as well as the products analysed from the UBE2D2-catalysed reaction (39, 47, and 79 kDa, for tri-, tetra- and octa-Ub, respectively, **Fig. S7**). Remarkably, all the modified lysines cluster on the face of Brd4^BD2^ closest to Rbx1, as well as on the unstructured N-terminal tail of the bromodomain (**Figure 2B**, lysine residues highlighted in pink). By contrast, lysines on the opposing face were not identified as being ubiquitinated in any of our samples (**Figure 2B**, lysine residues highlighted in blue). In an attempt to identify some preferential Lys and hint to ubiquitination specificity, we also excised bands at lower timepoints and identified Lys-modified peptides for a mono-Ub-Brd4 ^BD2^ product of the UBE2D2 reaction at 1 h (**Fig. S6-7**), and for a tetra-Ub-Brd4^BD2^ product of the UBE2R1 reaction at 30 min (**Figure 2A** and **Fig. S6-7**). While 7 out of 8 Lysines were found to be modified in the product of the UBE2D2-catalyzed reaction, suggesting a mixture of different monoubiquitinated Brd4^BD2^ products, we found only one peptide corresponding to K456 ubiquitination in the UBE2R1-catalysed tetra-Ub-Brd4^BD2^ product (**Figure 2C** and **Fig. S7**). It is likely that this difference reflects the relatively slower intrinsic catalytic activity of the elongating E2 UBE2R1 at placing the first ubiquitin on a substrate, as compared to UBE2D2, such that it is much more discriminatory for Lys residues at the short timepoints, thus suggesting that K456 may be one of the most preferentially ubiquitinated residues in the MZ1-bound Brd4-^BD2^ mechanism. There are caveats in that: a) other Lys-modified peptides may not have been detected, and b) the Adromeda score of 91 was just short of 100 (**Figure 2C** and **Fig. S7**). Nonetheless, K456 had the highest Andromeda scores amongst all modified lysines in the UBE2D2-catalyzed mono-Ub-Brd4 ^BD2^ product, and featured Andromeda scores of >100 in all the other analyzed samples. Analysis of the ubiquitin peptides in UBE2R1-produced tetra-Ub-Brd4^BD2^ sample exhibited a mixture of ubiquitin modification sites with a predominant intensity of K48 linkages, consistent with the proposition that UBE2R family E2s preferentially extend K48-linked chains (*58*), and suggesting that at least in part the sample contains multiply-ubiquitinated K456 modified Ub-Brd4 ^BD2^ species (**Fig. S7**).

**Figure 2.**
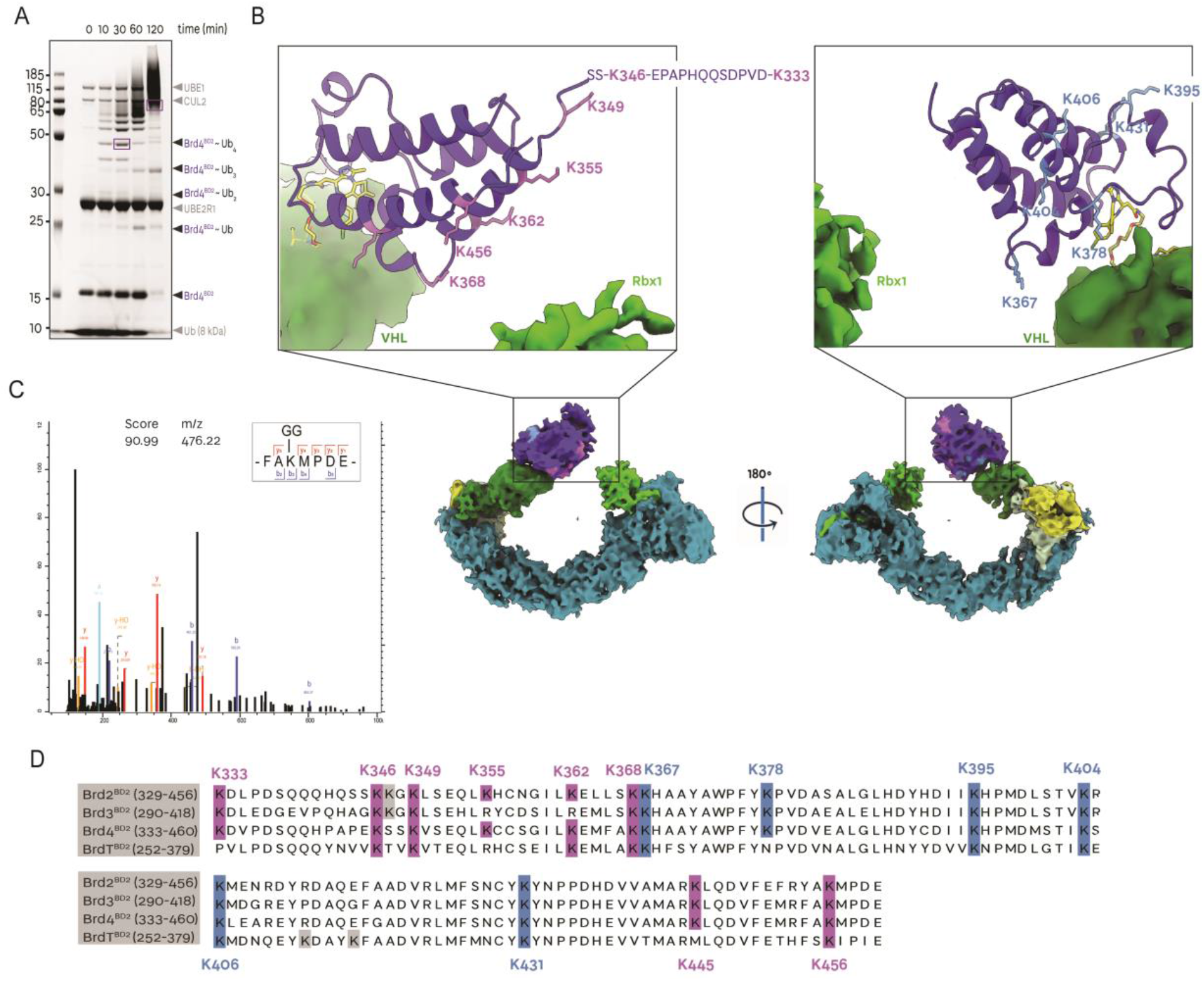
Brd4 bromodomains are ubiquitinated at lysines accessible to the E3 ligase RING box domain. (**A**) Coomassie-stained SDS-PAGE of *in vitro* Brd4^BD2^ ubiquitination in the presence of MZ1-CRL2^VHL^-UBE2R1-Ub (in purple: gel bands subjected to mass spectrometry analysis). (**B**) Cryo-EM volume of Brd4^BD2^ in complex with MZ1-(NEDD8)-CRL2^VHL^ with ubiquitinated lysines on Brd4^BD2^ highlighted in pink and unmodified lysines highlighted in light blue on as mapped by mass spectrometry following *in vitro* ubiquitination. (**C**) Identified mass spectrum for the Brd4^BD2^ K-GG modified peptide at K456. ‘Score’ refers to the Andromeda score. (**D**) Sequence alignments of the bromodomain 2 of Brd2, Brd3, Brd4 and BrdT. Ubiquitinated lysines on Brd4^BD2^ (as shown by mass spectrometry) are highlighted in pink and unmodified lysines highlighted in light blue, with sequence-aligned lysines of other BD2s represented in the same colour, suggesting which lysines may be modified in other BET BD2s.

As MZ1 exhibits ubiquitination and degradation selectivity for Brd4 and Brd2 over Brd3 (*13, 29, 40*), we inspected the conservation of the identified ubiquitinated lysine residues from sequence alignments of the BD2 in the different BET proteins (**Figure 2C** and **Fig. S8**). The K456 position is found to be strictly conserved amongst the BET BD2s, and so are K346, K349 and K368, while other two residues (K333 and K445) are strictly conserved amongst the ubiquitously expressed Brd2/3/4 but not on BrdT (**Figure 2C**). There are however two residue positions that bear a ubiquitinable Lys in Brd4^BD2^ and Brd2^BD2^ but a non-ubiquitinable Arg residue in Brd3^BD2^. This analysis suggests that, while the main contributor and driver of the mechanistic selectivity for Brd4^BD2^ and Brd2^BD2^ over Brd3^BD2^ reside largely on the preferential neo-substrate recognition within the ternary complex (*13, 29*), it is possible at least partially that the presence of these two ubiquitinable Lysine residues in Brd2/4 but not Brd3 could also contribute. Together, our ubiquitination data identifies a cluster of ubiquitinable lysine which reside on the face of the target protein closest to Rbx1, clearly distinct from those present in the opposite face which are non-ubiquitinable. We also identify a single Lys residue appears to be preferentially ubiquitinable, while overall lysine conservation and positioning cannot clearly explain the mechanistic selectivity of MZ1 for the different BET BD2s.

To better understand how cullin RING ligases mediate ubiquitination of *neo*-substrates, we developed a method to capture transition-state analogue species corresponding to the active ubiquitin chain extension on Ub-BRD4^BD2^ by UBE2R1 (**Figure 3A**, and **Fig. S9**). In this ubiquitination structural mimetic, wild-type Brd4^BD2^ was first N-terminally fused to the C-terminus of ubiquitin (G76S, K48C), mimicking an acceptor ubiquitin (Brd4^BD2^ -Ub^A^). A UBE2R1(C93K, S138C, C191S, C223S) mutant was loaded irreversibly with wild-type ubiquitin *via* a stable isopeptide bond at the C93K residue (UBE2R1-Ub^D^). The resulting Ub(G76S, K48C)-Brd4^BD2^ was biochemically crosslinked with a small bismaleimidoethane (BMOE) crosslinker to the S138C residue near the E2 acceptor site of UBE2R1(C93K, S138C, C191S, C223S)-Ub and purified (Brd4^BD2^-Ub^A^-BMOE-Ube2R1-Ub^D^) (**Figure 3A, Fig. S9**). Upon incubation with MZ1 and (NEDD8)-CRL^VHL^, full complex formation was observed by gel electrophoresis and mass photometry analyses (**Figure 3B-C**). Within the (NEDD8)-CRL2^VHL^-MZ1-Brd4^BD2^-Ub^A^-BMOE-Ube2R1-Ub^D^ complex, MZ1 and the cross-linked substrate-E2 conjugate serve as a stable bridge between the N- and C-terminal regions of CRL2^VHL^, with the goal of encircling the full complex into a closed ‘ring-like’ structure.

**Figure 3.**
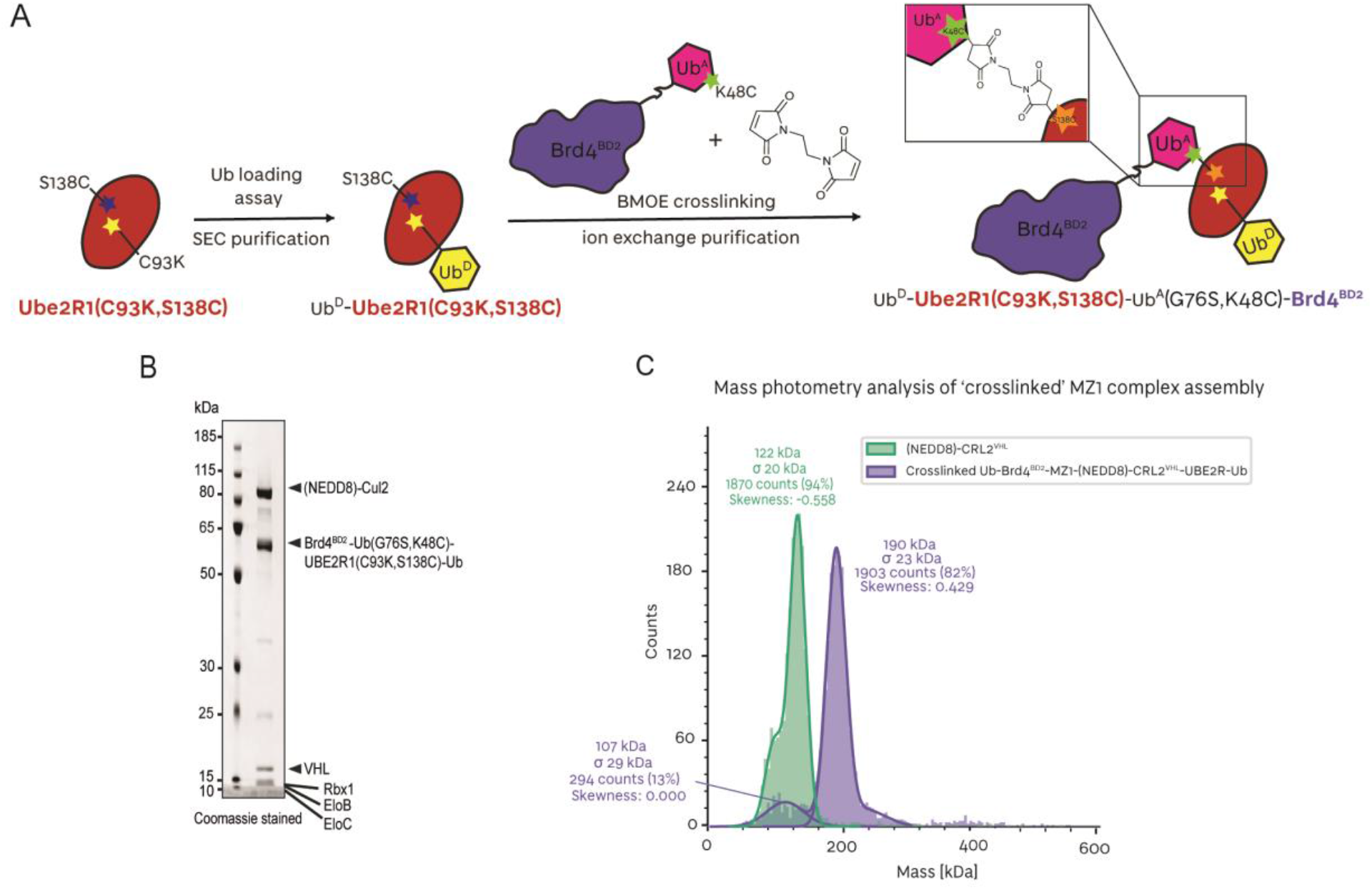
Assembly of a Brd4^BD2^ ubiquitination structural mimetic. (**A**) Schematic of the biochemical strategy used to trap MZ1-Brd4^BD2^-Ub(G76S, K48C)-UBE2R1(C93K, S138C, C191S, C223S)-Ub in complex with (NEDD8)-CRL2^VHL^. (**B**) SDS-PAGE of the assembled (NEDD8)-CRL2^VHL^-MZ1-Brd4^BD2^-Ub(G76S, K48C)-UBE2R1(C93K, S138C)-Ub complex applied to cryo-EM grids. (**C**) Mass photometry histogram of the fully assembled (NEDD8)-CRL2^VHL^-MZ1-Brd4^BD2^-Ub(G76S, K48C)-UBE2R1(C93K, S138C, C191S, C223S)-Ub complex (purple) applied to cryo-EM grids, versus (NEDD8)-CRL2^VHL^ (green). The peaks have been fitted with Gaussian curves.

To solve the structure of a closed fully-assembled complex, we subjected the sample to structural studies by cryo-EM (see **Fig. S10** for data analysis). Whereas the non-crosslinked ‘open’ cryo-EM structure suffered from anisotropy (**Figure 1** and **Fig. S3**), these anisotropic effects were reduced with this sample due to imaging in thicker ice of the cryogenic sample at 300 kV, allowing the particles to adopt a wider range of orientation distribution and achieve a higher overall resolution (**Fig. S10B, C** and **E**).

With this sample, we solved a number of discrete cryo-EM structures corresponding to fixed states, from a single multi-class *ab initio* classification. Although mass photometry suggested complete complex formation (**Figure 3C**), ‘State I’ corresponding to the (NEDD8)-CRL2^VHL^ was still present, possibly due to the complex partially dissociating in the cryogenic sample (**Figure 4A, left**). ‘State II’ corresponding to an open form of (NEDD8)-CRL2^VHL^-MZ1-Brd4^BD2^-Ub^A^-BMOE-Ube2R1-Ub^D^ with MZ1-Brd4^BD2^ visibly engaged with VHL and UBE2R1(C93K, S138C, C191S, C223S)-Ub tethered to Brd4^BD2^-Ub(G76S, K48C) but not engaging with Rbx1 at the C-terminus of the CRL2, and likely sampling many conformations around the complex (**Figure 4A, middle**). Notably, State II highlights and is consistent with the stability and longevity of the VHL-MZ1-Brd4^BD2^ ternary complex, and the rapid association/dissociation kinetics of UBE2R1-CRL2 interaction (*54*). We also obtained a population of species corresponding to a continuum between States III and IV, where the MZ1-Brd4^BD2^-Ub^A^-BMOE-Ube2R1-Ub^D^ central component appears to have successfully bridged the two sides of the (NEDD8)-CRL2^VHL^ by engaging the substrate receptor VHL at one end and the Rbx1-Cul2 C-terminal region at the other end. Owing to the highly dynamic states, the local resolution for the flexible components was reduced (**Fig. S10F**).

**Figure 4.**
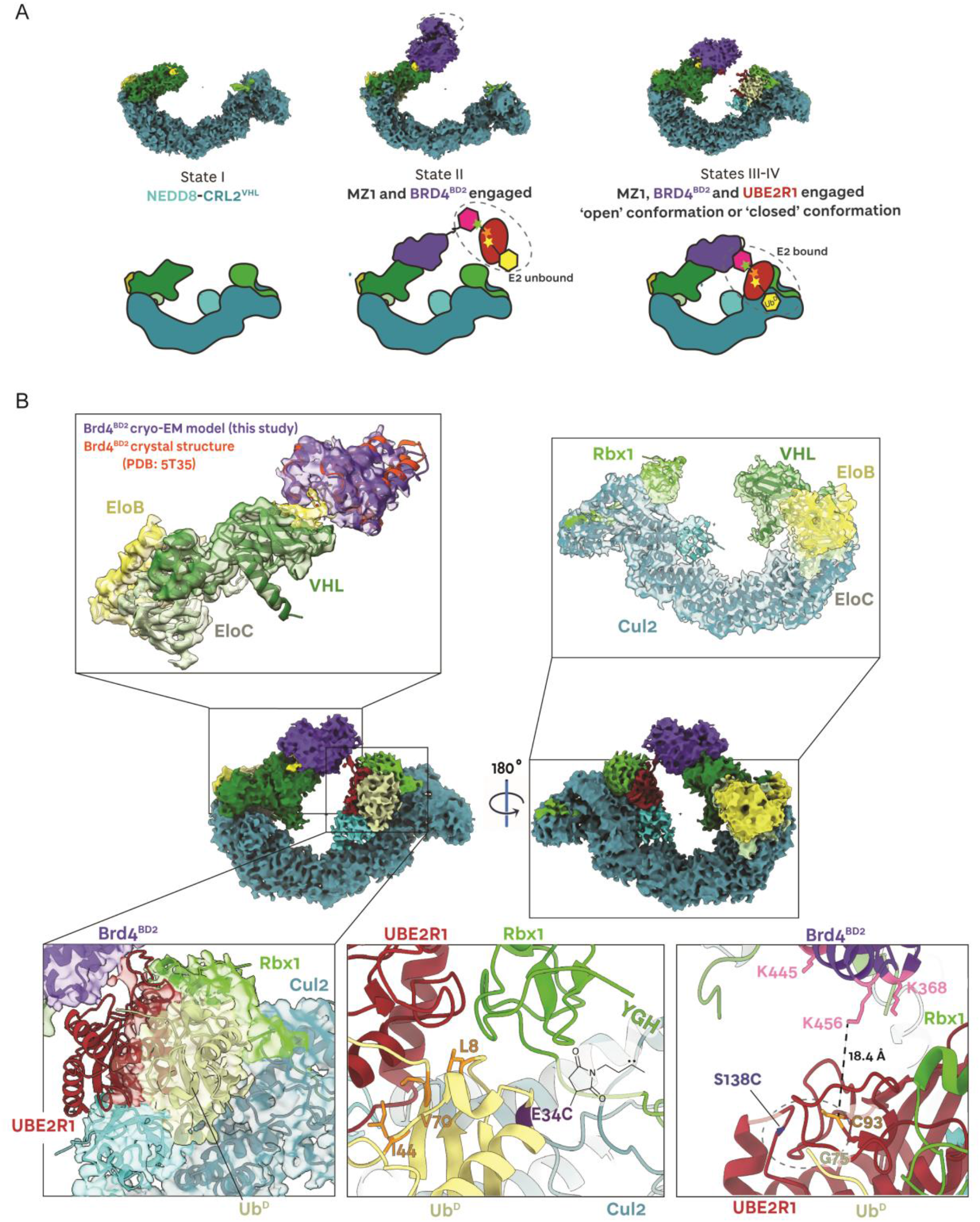
Structural snapshots of Brd4^BD2^ ubiquitination complex assembly. (**A**) Fixed discrete states and continuous states are captured by cryo-EM corresponding to the dynamics of the crosslinked (NEDD8)-CRL2^VHL^-MZ1-Brd4^BD2^-Ub(G76S, K48C)-UBE2R1(C93K, S138C)-Ub polyubiquitination species. (**B**) The ∼3.7 Å resolution cryo-EM reconstruction of ‘closed’ crosslinked (NEDD8)-CRL2^VHL^-MZ1-Brd4^BD2^-Ub(G76S, K48C)-UBE2R1(C93K, S138C)-Ub. The E3 ligase (NEDD8)-CRL2^VHL^ (shades of green) recruits MZ1 (yellow) which in turn binds the neo-substrate Brd4^BD2^ (purple). UBE2R1 (red) is engaged by Rbx1, with donor ubiquitin (pale yellow) covalenty bound to UBE2R1(C93K).

In recent years a number of deep learning neural network models and algorithms have been developed to explore hetereogeneity in cryo-EM samples, including 3DFlex (*59*), cryoDRGN (*60*), 3D variability analysis (3DVA) (*61*), and e2gmm (*62*) amongst others. To model the conformational landscape of the (NEDD8)-CRL2^VHL^-MZ1-Brd4^BD2^-Ub(G76S, K48C)-UBE2R1(C93K, S138C, C191S, C223S)-Ub complex, we used 3D variability analysis (*61*) to resolve continuous flexibility (**Supplementary Videos S1** and **S2**). For the population of species corresponding to States III-IV, we obtained a consensus volume corresponding to an average state. In this overall ∼3.7 Å reconstruction, (NEDD8)-CRL2^VHL^ is locally resolved to ∼3.5 Å, whereas Brd4^BD2^-Ub(G76S, K48C) and UBE2R1(C93K, S138C, C191S, C223S)-Ub components display resolutions in a range of 4 to 8 Å (**Fig. S10F**). From this consensus volume we were able to build an atomic model supported by in-solution NMR and structural cross-linking techniques and knowledge from previously published CRL-E2 structures and our previous open structure, revealing indeed a closing of the CRL ring as aimed for and potential interactions at the substrate:E2-Ub^D^ interface.

In this ‘closed’ structure, the VHL-MZ1-Brd4^BD2^ portion conserves its overall geometry (**Figure 4B, top left** and **Fig. S11**) with no major rearrangements seen. The same tilt relative to the cocrystal structure (*13*) is also seen in this closed state, albeit to a lesser extent than observed in the ‘open’ structure (**Fig. S11C-D**). As the resolution of this region was higher compared to the ‘open’ structure, we can now observed density volume for the PROTAC MZ1 itself (yellow in **Fig. S11A-B**). However, the resolution diminishes as the bromodomain moves away from VHL-MZ1, suggesting high-flexibility of the terminal regions, and no volume is observed corresponding to the acceptor ubiquitin bound to Brd4. The core of the CRL2^VHL^ ligase complex is extremely well resolved as the highest resolution region in the structure (**Figure 4B, top right**). Overlays with our ‘open’ structure and with our pentameric crystal structure across the whole of Cul2 shows the overall architecture is strictly conserved, with some minor bending over the first 160 residues of the Cul2 NTD (**Fig. S12**). Volume corresponding to NEDD8 is visible and could be modelled in (cyan in **Figure 4B, top right**), whereas the WHB domain is not.

Pleasingly, we observed enough volume corresponding to the UBE2R1-Ub^D^ conjugate that allowed us to model it bound to Rbx1 (red, yellow and green, respectively, in **Figure 4B, bottom left**). Although the volume corresponding to the E2 did not cover its entire structure, we could model this based on the good-quality density for Ub^D^ and the available co-crystal structure of RING-E2∼Ub^D^ (*52*) amongst others, as well as the cryo-EM structure of E2 bound to full CRL which guided positioning relative to NEDD8 and the C-terminal region of the Cullin (*30*), giving us confidence on the positioning of the key catalytic E2∼Ub^D^ unit in the structure.

As no structure has been determined for UBE2R1/2 E2 enzymes bound to any E3 ligase, we then used protein-observed NMR spectroscopy (*63*) to determine how UBE2R1 interacts with Ub^D^ in solution in the context of the UBE2R1∼Ub conjugate. Analysis of the ^15^N-^1^H HSQC NMR spectrum of ^15^N-labelled Ub^D^ loaded on UBE2R1(C93K) showed large chemical shift perturbations and signal attenuation, corresponding to clear stabilization of Ub^D^ on the E2, in particular through the canonical hydrophobic patch including residues L8, I44, and V70 (**Fig.S13**). Crucially, we observe that this hydrophobic patch surface is fully buried in contact with UBE2R1 also in our cryo-EM structure, evidencing the UBE2R1∼Ub bound in a closed, pre-catalytic state (**Figure 4B, bottom middle**). Several interactions between the E2 and Rbx1 were observed as expected. Nonetheless, the modelled Asp102 and Asp103 residues on the acidic loop of UBE2R1 were found close to key basic residues on Rbx1 known to interact with, such as Arg91 (*64*). Previous work has shown that the UBE2R acidic tail binds a basic ‘canyon’ on Cul1 that is highly conserved throughout all Cullins (*54*). We could not identify volume for the acidic region of UBE2R1 in our cryo-EM structure, hence could not confidently model this. To further validate interactions in this highly dynamic RING-E2-Ub^D^ region of our cryo-EM structure, we designed a ubiquitin-directed photoreactive probe (UDPRP) and used it in cross-linking mass spectrometry experiments. We synthesized the photocrosslinker *N*-maleimido-diazirine (see Experimental Methods and **Fig. S15**) that would site-specifically react with E34C mutant ubiquitin, chosen based on its proximity to Rbx1 in existing structures (*30*), and formed a stable and active isopeptide-linked Ub-Ube2R1 conjugate via the E2 C93K (see **Fig. S15A-E**). In the photocrosslinking assay containing neddylated-CRL2^VHL^ the UDPRP photo-crosslinked to Rbx1 (**Fig. S16F-G**). The photo-crosslinked product was excised from the gel and analyzed by mass-spectrometry to identify insertion in the ^106^YGH^108^ sequence at the flexible C-terminus of Rbx1 (**Fig. S16H-I**). While density was not visible for the terminal YGH tail of Rbx1 in our structure, the distance of ∼18 Å between Ub^D^(E34C) and K105 right upstream of YGH in Rbx1 is consistent with positioning of the crosslinker-YGH (**Figure 4B, bottom middle**). Together our complementary structural NMR and cross-linking studies in solution validate our models of the E2∼Ub:Rbx1 components in the cryo-EM structure volume.

To gain structural insights into preferences for acceptor Lysine on the *neo*-substrate, we inspected closely the interface between UBE2R1 and Brd4^BD2^, and measured distances with the ubiquitinable lysine residues of Brd4 (**Figure 4B, bottom right**, and **Fig S16**). While Brd4^BD2^ and UBE2R1 are not in direct contact, we observed that K456, which we had found as preferentially ubiquitinated, points directly towards the E2∼Ub electrophilic site (**Figure 4B, bottom right**). We measured a distance of 18.4 Å between the N-epsilon of K456 and the carbonyl of the modelled thioester at C93 (**Figure 4B, bottom right**, and **Fig S16**). Albeit not at shortest distance, it appears that K456 is primed for an optimal geometry to approach the electrophilic thioester on the E2∼Ub^D^. As K456 is located at the C-terminal end of the C-helix of the bromodomain, distal to the bound MZ1 interface, the shown tilt could help to bridge such a distance. Residues K368 and K445 are both also close to C93, but in constrast with K456 they are much closer to the VHL-MZ1-Brd4^BD2^ interface, and have thus far less mobility. Based on these observations, we posit they should be less preferred for ubiquitination than K456. Together our structural work exploited cross-linking stabilization to achieve a high-resolution cryo-EM structure for a fully closed (NEDD8)-CRL2^VHL^-PROTAC-target-Ub^A^-E2∼Ub^D^ structure that we validate using in-solution technique and that allowed to gain unprecedented insights into *neo*-substrate ubiquitination as catalyzed by a PROTAC-bound CRL machinery.

## Discussion

Degrader drugs co-opt the catalytic activity of ubiquitin E3 Cullin RING ligases to drive efficient ubiquitination and degradation of disease-causing proteins (*5, 6*). Catalytic activity and substrate specificity of CRLs are critically determined by the spatial organization and relative interactions of their subunits, and by their ability to work in concert in order to achieve flexibility (*30, 56*). These are important requirements not only to bring together a substrate and ubiquitin that would otherwise be too far apart, but also to allow hitting the substrate at multiple positions and to accommodate the building of ubiquitin chains during the catalytic cycles (*58*). Here our objective was to determine the structural and mechanistic bases of how optimal degraders mediate productive target ubiquitination. This information is required to establish the degradability or PROTACability of targets, for which rules have remained unclear (*65, 66*). By combining cryo-EM structures, cross-linking trapping strategies and biochemical identification of ubiquitinated lysines, we determine the structure and mechanism of the entire PROTAC-induced Cullin 2 RING ligase VHL catalytic machinery, and how it operates to catalyse ubiquitination on its highly specific *neo*-substrate – the second bromodomain of Brd4 – that is recruited to VHL by the PROTAC MZ1.

Our results explain the *‘ubiquitinability’* of Brd4^BD2^ and allow us to establish general principles regarding the structural requirements for *neo*-substrate ubiquitination specificity (**Figure 5**). We show that Brd4^BD2^, once tightly glued to VHL by MZ1, adopts a preferred orientation projecting towards the RING activated E2-ubiquitin catalytic module of the CRL. This geometry projects one face of the substrate towards the E2, such that that lysine residues on this surface are within range of reacting with the Ub∼E2 thioester, i.e. the ‘ubiquitination zone’ (*33*), and are thus susceptible to ubiquitin modification. Ubiquitinated residues are on a ‘light face’, as though they would be illuminated by a source of light projecting from the E2∼Ub^D^ (**Figure 5, top left**). This suggests that if a lysine residue is present on the ‘light face’ of the target protein it has the potential for ubiquitin modification. In contrast, lysine residues present on the opposite ‘dark face’ of the protein cannot be ubiquitinated. The arrangement of ‘light’ and ‘dark’ faces on a substrate will be determined by its orientation relative to the E3 ligase as dictated by the degrader. Tight, stably bound ternary complexes are necessary for fast, potent protein degradation (*13, 29, 34-39*). Here we show that depending on the geometry of such stable complex, different ‘light faces’ could arise on the target protein. For a given E3 ligase, varying either of the other two components within the ternary complex could lead to very different orientations of the target protein relative to the E2∼Ub, different ‘light faces’, and so different degreed of ubiquitinability. Examples of this include when varying chemical structures of PROTACs for the same target, as done in drug discovery optimization projects; as well as for a given degrader against different target mutants, isoforms or paralogues, as is the case here for MZ1 with the various BET bromodomains (*13, 29*). Recruiting the substrate in a disfavoured orientation, pointing away from the E2∼Ub site, would result in less efficient ubiquitination, hence less effective degraders (**Figure 5, top right**).

**Fig. 5.**
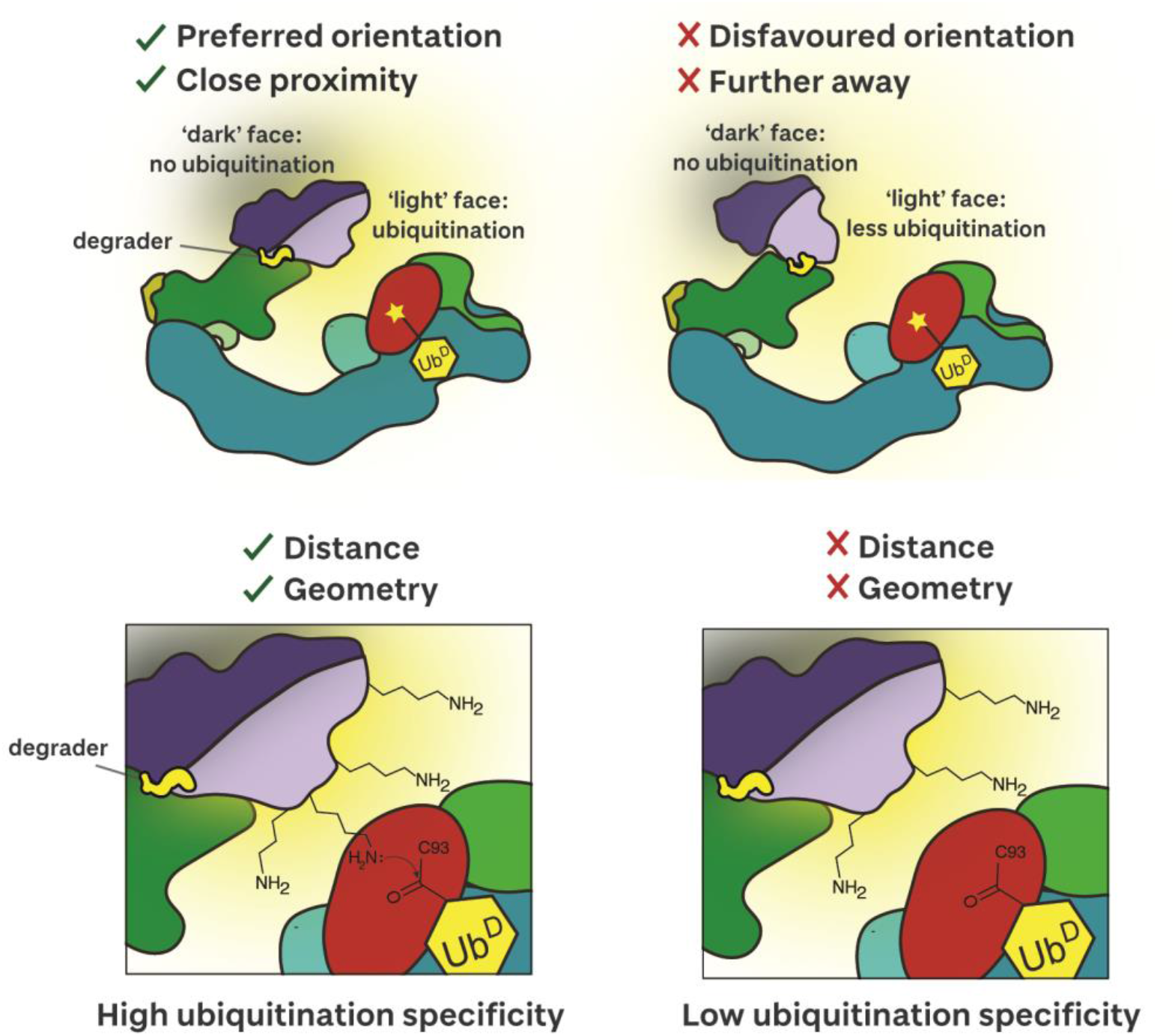
Schematic of structural and mechanistic feature of ‘*ubiquitinability*’ of target proteins revealed from this study.

Our data together is also consistent with the notion that, even within a highly favoured system such as VHL-MZ1-Brd4^BD2^, which bears several lysine residues on the ‘light face’, not all lysine residues will be equal as their specific position, geometry and reactivity will all be factors in determining their efficiency for ubiquitination. We find that a single lysine (K456) on the ‘light face’ of Brd4^BD2^ is preferentially ubiquitinated by CRL2^VHL^. Thus, to achieve fast, potent and profound target degradation, the degrader must recruit the target not only in a stable, long-lived complex at low concentration, but also orienting it favourably towards the catalytic site. In addition to this, our work shows that it is also important that one or more lysine residues within the ‘light face’ are properly positioned with respect to both distance and geometry to drive highly specific and efficient ubiquitination (**Figure 5, bottom**).

Together, these fresh mechanistic insights advance our knowledge of degraders’ mode of action, further explain the intra-BET specificity of the archetypical PROTAC MZ1, and reveal new guiding rules for understanding and manipulating the exquisite degradation specificity of PROTACs and molecular glues. While this work has focused on CRL2^VHL^ as E3 ligase and Brd4 as target protein of interest, these conclusions are expected to be general and apply to other E3 ligases e.g. CRL4^CRBN^ and beyond, and to other proteins that are targeted for degradation (*67*).

We therefore anticipate that our findings will inspire in-depth mechanistic studies and will guide more refined structure-based rational optimization of improved degrader drugs.

## Acknowledgements

We acknowledge the technical and research staff of the Ciulli Laboratories at CeTPD for the set-up and upkeep of protein purification and computational infrastructure, and Alena Kroupova for discussions. We thank Emma Branigan (University of Dundee) for the gift of pure recombinant ubiquitin E34C protein. We thank Ramasubramanian Sundaramoorthy for access and support with the Dundee’s School of Life Sciences in-house Cryo-EM Facility; Paula da Fonseca and Ed Morris (University of Glasgow) for microscopy support at eBIC; and the team at the Dundee MRC PPU Proteomics Facility for running the mass-spectrometry samples. We acknowledge Diamond for access and support of the cryo-EM facilities at the UK national electron Bio-Imaging Centre (eBIC), proposal BI31827-13, funded by the Wellcome Trust, MRC and BBSRC. C.C. thanks the Cold Spring Harbor Laboratory course on cryo-electron microscopy instructors and organisers for general training in single particle cryo-EM.

## Author contributions

C.C., M.A.N. and A.C. conceived and planned this project. C.C., M.A.N. and S.C. expressed and purified recombinant proteins and performed biochemical assays. C.C. and M.A.N. performed cryo-EM sample preparation, imaging, data processing and modelling. S.C. and M.T. performed mass spectrometry experiments and data analysis. N.M. synthesised compounds. C.C. and A.C. analysed and interpreted data, and co-wrote the manuscript with input from all co-authors. A.C. and R.T.H. supervised this project and acquired funding.

## Data and materials availability

Atomic models have been deposited to the Protein Data Bank (PDB) under IDs: 8RWZ and 8RX0. Electron microscopy reconstructions have been deposited to the Electron Microscopy Databank (EMDB) under accession codes: EMD-19569 and EMD-19567. MS data has been deposited to PRIDE under accession codes: PXD049047, PXD049043, PXD049050.

## References

1. L. Zhao, J. Zhao, K. Zhong, A. Tong, D. Jia, Targeted protein degradation: mechanisms, strategies and application. Signal Transduct Target Ther 7, 113 (2022).

2. D. Chirnomas, K. R. Hornberger, C. M. Crews, Protein degraders enter the clinic - a new approach to cancer therapy. Nat Rev Clin Oncol 20, 265–278 (2023).

3. N. Zheng, N. Shabek, Ubiquitin Ligases: Structure, Function, and Regulation. Annu Rev Biochem 86, 129–157 (2017).

4. J. W. Harper, B. A. Schulman, Cullin-RING Ubiquitin Ligase Regulatory Circuits: A Quarter Century Beyond the F-Box Hypothesis. Annu Rev Biochem 90, 403–429 (2021).

5. A. D. Cowan, A. Ciulli, Driving E3 Ligase Substrate Specificity for Targeted Protein Degradation: Lessons from Nature and the Laboratory. Annu Rev Biochem 91, 295–319 (2022).

6. M. Bekes, D. R. Langley, C. M. Crews, PROTAC targeted protein degraders: the past is prologue. Nat Rev Drug Discov 21, 181–200 (2022).

7. G. Dong, Y. Ding, S. He, C. Sheng, Molecular Glues for Targeted Protein Degradation: From Serendipity to Rational Discovery. J Med Chem 64, 10606–10620 (2021).

8. M. Zengerle, K. H. Chan, A. Ciulli, Selective Small Molecule Induced Degradation of the BET Bromodomain Protein BRD4. ACS Chem Biol 10, 1770–1777 (2015).

9. G. E. Winter et al., DRUG DEVELOPMENT. Phthalimide conjugation as a strategy for in vivo target protein degradation. Science 348, 1376–1381 (2015).

10. D. P. Bondeson et al., Catalytic in vivo protein knockdown by small-molecule PROTACs. Nat Chem Biol 11, 611–617 (2015).

11. A. Ciulli et al., The 17(th) EFMC Short Course on Medicinal Chemistry on Small Molecule Protein Degraders. ChemMedChem 18, e202300464 (2023).

12. T. A. F. Cardote, M. S. Gadd, A. Ciulli, Crystal Structure of the Cul2-Rbx1-EloBC-VHL Ubiquitin Ligase Complex. Structure 25, 901–911 e903 (2017).

13. M. S. Gadd et al., Structural basis of PROTAC cooperative recognition for selective protein degradation. Nat Chem Biol 13, 514–521 (2017).

14. C. J. Diehl, A. Ciulli, Discovery of small molecule ligands for the von Hippel-Lindau (VHL) E3 ligase and their use as inhibitors and PROTAC degraders. Chem Soc Rev 51, 8216–8257 (2022).

15. E. S. Fischer et al., Structure of the DDB1-CRBN E3 ubiquitin ligase in complex with thalidomide. Nature 512, 49–53 (2014).

16. P. P. Chamberlain et al., Structure of the human Cereblon-DDB1-lenalidomide complex reveals basis for responsiveness to thalidomide analogs. Nat Struct Mol Biol 21, 803–809 (2014).

17. R. P. Nowak et al., Plasticity in binding confers selectivity in ligand-induced protein degradation. Nat Chem Biol 14, 706–714 (2018).

18. J. Yamamoto, T. Ito, Y. Yamaguchi, H. Handa, Discovery of CRBN as a target of thalidomide: a breakthrough for progress in the development of protein degraders. Chem Soc Rev 51, 6234–6250 (2022).

19. V. Oleinikovas, P. Gainza, T. Ryckmans, B. Fasching, N. H. Thoma, From Thalidomide to Rational Molecular Glue Design for Targeted Protein Degradation. Annu Rev Pharmacol Toxicol 64, 291–312 (2024).

20. T. Ishida, A. Ciulli, E3 Ligase Ligands for PROTACs: How They Were Found and How to Discover New Ones. SLAS Discov 26, 484–502 (2021).

21. M. Schroder et al., DCAF1-based PROTACs with activity against clinically validated targets overcoming intrinsic-and acquired-degrader resistance. Nat Commun 15, 275 (2024).

22. C. M. Hickey et al., Co-opting the E3 ligase KLHDC2 for targeted protein degradation by small molecules. Nat Struct Mol Biol (2024) Jan 4. doi: 10.1038/s41594-023-01146-w.

23. A. Hanzl et al., E3-Specific Degrader Discovery by Dynamic Tracing of Substrate Receptor Abundance. J Am Chem Soc 145, 1176–1184 (2023).

24. O. Hsia et al., Targeted protein degradation via intramolecular bivalent glues. BioRxiv (2023) 2023.02.14.528511. doi: 10.1101/2023.02.14.528511

25. M. Slabicki et al., The CDK inhibitor CR8 acts as a molecular glue degrader that depletes cyclin K. Nature 585, 293–297 (2020).

26. S. J. Hughes, A. Ciulli, Molecular recognition of ternary complexes: a new dimension in the structure-guided design of chemical degraders. Essays Biochem 61, 505–516 (2017).

27. R. Casement, A. Bond, C. Craigon, A. Ciulli, Mechanistic and Structural Features of PROTAC Ternary Complexes. Methods Mol Biol 2365, 79–113 (2021).

28. B. E. Smith et al., Differential PROTAC substrate specificity dictated by orientation of recruited E3 ligase. Nat Commun 10, 131 (2019).

29. M. J. Roy et al., SPR-Measured Dissociation Kinetics of PROTAC Ternary Complexes Influence Target Degradation Rate. ACS Chem Biol 14, 361–368 (2019).

30. K. Baek et al., NEDD8 nucleates a multivalent cullin-RING-UBE2D ubiquitin ligation assembly. Nature 578, 461–466 (2020).

31. N. Bai et al., Modeling the CRL4A ligase complex to predict target protein ubiquitination induced by cereblon-recruiting PROTACs. J Biol Chem 298, 101653 (2022).

32. D. Lv et al., Development of a BCL-xL and BCL-2 dual degrader with improved anti-leukemic activity. Nat Commun 12, 6896 (2021).

33. T. Dixon et al., Predicting the structural basis of targeted protein degradation by integrating molecular dynamics simulations with structural mass spectrometry. Nat Commun 13, 5884 (2022).

34. D. P. Bondeson et al., Lessons in PROTAC Design from Selective Degradation with a Promiscuous Warhead. Cell Chem Biol 25, 78–87 e75 (2018).

35. W. Farnaby et al., BAF complex vulnerabilities in cancer demonstrated via structure-based PROTAC design. Nat Chem Biol 15, 672–680 (2019).

36. R. P. Law et al., Discovery and Characterisation of Highly Cooperative FAK-Degrading PROTACs. Angew Chem Int Ed Engl 60, 23327–23334 (2021).

37. P. S. Dragovich et al., Antibody-Mediated Delivery of Chimeric BRD4 Degraders. Part 1: Exploration of Antibody Linker, Payload Loading, and Payload Molecular Properties. J Med Chem 64, 2534–2575 (2021).

38. S. Imaide et al., Trivalent PROTACs enhance protein degradation via combined avidity and cooperativity. Nat Chem Biol 17, 1157–1167 (2021).

39. R. P. Wurz et al., Affinity and cooperativity modulate ternary complex formation to drive targeted protein degradation. Nat Commun 14, 4177 (2023).

40. K. M. Riching et al., Quantitative Live-Cell Kinetic Degradation and Mechanistic Profiling of PROTAC Mode of Action. ACS Chem Biol 13, 2758–2770 (2018).

41. A. Testa, S. J. Hughes, X. Lucas, J. E. Wright, A. Ciulli, Structure-Based Design of a Macrocyclic PROTAC. Angew Chem Int Ed Engl 59, 1727–1734 (2020).

42. R. Beveridge et al., Native Mass Spectrometry Can Effectively Predict PROTAC Efficacy. ACS Cent Sci 6, 1223–1230 (2020).

43. J. H. Song et al., Native mass spectrometry and gas-phase fragmentation provide rapid and in-depth topological characterization of a PROTAC ternary complex. Cell Chem Biol 28, 1528–1538 e1524 (2021).

44. A. Hanzl et al., Functional E3 ligase hotspots and resistance mechanisms to small-molecule degraders. Nat Chem Biol 19, 323–333 (2023).

45. J. Krieger et al., Systematic Potency and Property Assessment of VHL Ligands and Implications on PROTAC Design. ChemMedChem 18, e202200615 (2023).

46. H. C. Nguyen, H. Yang, J. L. Fribourgh, L. S. Wolfe, Y. Xiong, Insights into Cullin-RING E3 ubiquitin ligase recruitment: structure of the VHL-EloBC-Cul2 complex. Structure 23, 441–449 (2015).

47. L. Gong, E. T. Yeh, Identification of the activating and conjugating enzymes of the NEDD8 conjugation pathway. J Biol Chem 274, 12036–12042 (1999).

48. D. T. Huang et al., E2-RING expansion of the NEDD8 cascade confers specificity to cullin modification. Mol Cell 33, 483–495 (2009).

49. D. M. Duda et al., Structural insights into NEDD8 activation of cullin-RING ligases: conformational control of conjugation. Cell 134, 995–1006 (2008).

50. N. H. Saifee, N. Zheng, A ubiquitin-like protein unleashes ubiquitin ligases. Cell 135, 209–211 (2008).

51. K. Wang, K. M. Reichermeier, X. Liu, Quantitative analyses for effects of neddylation on CRL2(VHL) substrate ubiquitination and degradation. Protein Sci 30, 2338–2345 (2021).

52. A. Plechanovova, E. G. Jaffray, M. H. Tatham, J. H. Naismith, R. T. Hay, Structure of a RING E3 ligase and ubiquitin-loaded E2 primed for catalysis. Nature 489, 115–120 (2012).

53. R. A. Chong et al., Pivotal role for the ubiquitin Y59-E51 loop in lysine 48 polyubiquitination. Proc Natl Acad Sci U S A 111, 8434–8439 (2014).

54. G. Kleiger, A. Saha, S. Lewis, B. Kuhlman, R. J. Deshaies, Rapid E2-E3 assembly and disassembly enable processive ubiquitylation of cullin-RING ubiquitin ligase substrates. Cell 139, 957–968 (2009).

55. A. Saha, R. J. Deshaies, Multimodal activation of the ubiquitin ligase SCF by Nedd8 conjugation. Mol Cell 32, 21–31 (2008).

56. D. Horn-Ghetko et al., Ubiquitin ligation to F-box protein targets by SCF-RBR E3-E3 super-assembly. Nature 590, 671–676 (2021).

57. H. Zhou, M. S. Zaher, J. C. Walter, A. A.-O. Brown, Structure of CRL2Lrr1, the E3 ubiquitin ligase that promotes DNA replication termination in vertebrates. Nucleic Acids Res 49, 13194–13206 (2021).

58. K. Wu, J. Kovacev, Z. Q. Pan, Priming and extending: a UbcH5/Cdc34 E2 handoff mechanism for polyubiquitination on a SCF substrate. Mol Cell 37, 784–796 (2010).

59. A. Punjani, D. J. Fleet, 3DFlex: determining structure and motion of flexible proteins from cryo-EM. Nat Methods 20, 860–870 (2023).

60. E. D. Zhong, T. Bepler, B. Berger, J. H. Davis, CryoDRGN: reconstruction of heterogeneous cryo-EM structures using neural networks. Nat Methods 18, 176–185 (2021).

61. A. Punjani, D. J. Fleet, 3D variability analysis: Resolving continuous flexibility and discrete heterogeneity from single particle cryo-EM. J Struct Biol 213, 107702 (2021).

62. M. Chen, S. J. Ludtke, Deep learning-based mixed-dimensional Gaussian mixture model for characterizing variability in cryo-EM. Nat Methods 18, 930–936 (2021).

63. J. N. Pruneda, K. E. Stoll, L. J. Bolton, P. S. Brzovic, R. E. Klevit, Ubiquitin in motion: structural studies of the ubiquitin-conjugating enzyme approximately ubiquitin conjugate. Biochemistry 50, 1624–1633 (2011).

64. S. Hill, J. S. Harrison, S. M. Lewis, B. Kuhlman, G. Kleiger, Mechanism of Lysine 48 Selectivity during Polyubiquitin Chain Formation by the Ube2R1/2 Ubiquitin-Conjugating Enzyme. Mol Cell Biol 36, 1720–1732 (2016).

65. M. Schneider et al., The PROTACtable genome. Nat Rev Drug Discov 20, 789–797 (2021).

66. W. Zhang et al., Machine Learning Modeling of Protein-intrinsic Features Predicts Tractability of Targeted Protein Degradation. Genomics Proteomics Bioinformatics 20, 882–898 (2022).

67. Y. Liu et al., Expanding PROTACtable genome universe of E3 ligases. Nat Commun 14, 6509 (2023).

