## Supplementary Materials for "Mechanism of degrader-targeted protein ubiquitinability"

**The PDF file includes:**

Materials and Methods
Figs. S1 to S16
Tables S1 to S2
Movies S1 to S2
References 1 to 35

### Materials and Methods

Protein expression and purification

**VCB** (VHL residues 54-213, EloB residues 1-104 and EloC residues 17-112) was expressed and purified as previously described. In short, single colonies of cDNA transformed into *E. coli* BL21(DE3) cells were grown overnight in lysogeny broth (LB) supplemented with ampicillin (100 μg $mL^{-1}$) and streptomycin (50 μg $mL^{-1}$) at 37 °C with shaking. The overnight culture was diluted 1:100 in LB supplemented with ampicillin (100 μg $mL^{-1}$) and streptomycin (50 μg $mL^{-1}$) and grown to an optical density (OD600) of ~0.6. Protein expression was induced overnight at 18 °C with 0.3 mM isopropyl β-d-1-thiogalactopyranoside (IPTG). The cells were harvested by centrifugation and frozen at -80 °C as pellets until further purification. The bacterial pellets were resuspended in buffer and lysed by cell disruption using as One Shot Cell Disruptor (Constant Systems) operating at 4 °C. Cellular debris was removed by centrifugation. $His_6$-tagged proteins were purified on a HisTrap$^{TM}$ FF Ni NTA affinity column (Cytiva) and eluted with an imidazole concentration gradient. The protein was dialysed into a low concentration imidazole buffer, incubated with TEV protease, and flowed through a HisTrap FF Ni NTA affinity column. VCB was additionally purified by anion exchange using a HiTrap$^{TM}$ Q HP column. The protein sample was finally purified by size exclusion chromatography on a HiLoad$^{TM}$ 16/600 Superdex$^{TM}$ 75 pg column (Cytiva) in 20 mM HEPES, pH 7.5, 150 mM sodium chloride and 0.5 mM TCEP, concentrated, flash frozen with liquid nitrogen and stored at -80 °C.

**NEDD8** (residues 1-76) Single colonies of cDNA transformed into *E. coli* BL21(DE3) cells were grown overnight in lysogeny broth (LB) supplemented with ampicillin (100 μg $mL^{-1}$) at 37 °C with shaking. The overnight culture was diluted 1:100 in 9 L of LB supplemented with ampicillin (100 μg $mL^{-1}$) and grown to an optical density (OD600) of ~0.6. Protein expression was induced overnight at 16 °C with 0.5 mM isopropyl β-d-1-thiogalactopyranoside (IPTG). The cells were harvested by centrifugation and frozen at -80 °C as pellets until further purification. The bacterial pellets were resuspended in 30 mM Tris, 200 mM NaCl, 5 mM DTT, pH 7.5, supplemented with 5 mM $MgCl_2$, 1 μg/mL DNAse I and 1X EDTA-free Roche protease inhibitor cocktail, and lysed by cell disruption using as One Shot Cell Disruptor (Constant Systems) operating at 4 °C. Cellular debris was removed by centrifugation. The $His_6$-tagged protein was purified on a HisTrap$^{TM}$ FF Ni NTA affinity column (Cytiva) and eluted with an imidazole concentration gradient (20 mM to 500 mM). The resulting eluate was incubated with 10 mL Glutathione Sepharose 4B resin (Cytiva). The protein-bound resin was washed with 50 mM Tris-HCl, 200 mM NaCl, 5 mM DTT, pH 7.6. TEV was added to the resin, and NEDD8 was cleaved at room temperature for 3 hours. NEDD8 was finally purified by size exclusion chromatography on a HiLoad$^{TM}$ 16/600 Superdex$^{TM}$ 75 pg column (Cytiva) in 20 mM HEPES, pH 7.5, 150 mM sodium chloride and 0.5 mM TCEP, concentrated, flash frozen with liquid nitrogen and stored at -80 °C.

**APPBP1-Uba3** and **Ube2M** Single colonies of cDNA (*1, 2*) transformed into *E. coli* BL21(DE3) cells were grown overnight in lysogeny broth (LB) supplemented with ampicillin (100 μg $mL^{-1}$) at 37 °C with shaking. The overnight culture was diluted 1:100 in 24 L of LB supplemented with ampicillin (100 μg $mL^{-1}$) and grown to an optical density (OD600) of ~0.6. Protein expression was induced overnight at 16 °C with 0.5 mM isopropyl β-d-1-thiogalactopyranoside (IPTG). The cells were harvested by centrifugation and frozen at -80 °C as pellets until further purification. The bacterial pellets were resuspended in 50 mM Tris, 200 mM NaCl, 5 mM DTT, pH 7.5, supplemented with 5 mM $MgCl_2$, 1 µg/mL DNAse I and 1X EDTA-free Roche protease inhibitor cocktail, and lysed by cell disruption using as One Shot Cell Disruptor (Constant Systems) operating at 4 °C. Cellular debris was removed by centrifugation. The lysate was incubated with 10 mL Glutathione Sepharose 4B (Cytiva) resin for 1 hour at room temperature. The resin was washed with 50 mM Tris-HCl, 200 mM NaCl, 5 mM DTT, pH 7.6 and the protein was eluted with 50 mM Tris, 200 mM NaCl, 5 mM DTT, pH 8.0, 10 mM reduced L-glutathione. 1 unit of thrombin per mg of total protein was added and the mixture was incubated overnight at -4 °C. APP-BP1-Uba3 was further purified by size exclusion chromatography on a HiLoad™ 16/600 Superdex™ 200 pg column (Cytiva) in 20 mM HEPES, pH 7.5, 150 mM sodium chloride and 0.5 mM TCEP. Ube2M was further purified by size exclusion chromatography on a HiLoad™ 16/600 Superdex™ 75 pg column (Cytiva) in 20 mM HEPES, pH 7.5, 150 mM sodium chloride and 0.5 mM TCEP. The resulting proteins was flowed over Glutathione Sepharose 4B resin to remove any trace amounts of GST-APP-BP1-Uba3, GST-Ube2M and GST. The purified protein samples were each concentrated, flash frozen with liquid nitrogen and stored at -80 °C.

**NEDD8-CRL2$^{VHL}$** Single colonies of Cul2-Rbx1 cDNA (*3*) transformed into *E. coli* BL21(DE3) cells were grown overnight in lysogeny broth (LB) supplemented with ampicillin (100 µg $mL^{-1}$) at 37 °C with shaking. The overnight culture was diluted 1:100 in 24 L of LB supplemented with ampicillin (100 µg $mL^{-1}$) and grown to an optical density (OD600) of ~0.6. Protein expression was induced for 16 hours at 16 °C with 0.2 mM isopropyl β-d-1-thiogalactopyranoside (IPTG). The cells were harvested by centrifugation and frozen at -80 °C as pellets until further purification. The cells were thawed, resuspended in 30 mM Tris-HCl, 200 mM NaCl, 5 mM DTT, pH 7.5, supplemented with 5 mM $MgCl_2$, 1 µg/mL DNAse I and 1X EDTA-free Roche protease inhibitor cocktail, and lysed by cell disruption at 30 kpsi. Cellular debris was removed by centrifugation. Cul2-Rbx1 was purified on a 5 mL HisTrap™ FF Ni NTA affinity column (Cytiva) and eluted with an imidazole concentration gradient from 0 to 300 mM. The protein was desalted into 20 mM HEPES, 150 mM NaCl, 0.5 mM TCEP, 5% (v/v) glycerol, pH 7.5 with a HiPrep 26/10 desalting column (Cytiva) and incubated with TEV protease and an excess of recombinant VHL-EloB-EloC. VHL-EloB-EloC-Cul2-Rbx1 (CRL2$^{VHL}$) was purified on a 5 mL StrepTrap XT column, washing with 100 mM Tris-HCl, 150 mM NaCl, pH 8.0 and eluting in 100 mM Tris-HCl, 300 mM NaCl, 50 mM biotin and 5% (v/v) glycerol, pH 8.0. Purified CRL2$^{VHL}$ (1.7 µM) was incubated with Uba3-APP-BP1 (250 nM), Ube2M (1.2 µM), NEDD8 (20 µM), ATP (1 mM) and $MgCl_2$ (5 mM) for 10 minutes at 37 °C. NEDD8-CRL2$^{VHL}$ was finally purified by size exclusion chromatography on a HiLoad™ 16/600 Superdex™ 200 pg column (Cytiva) in 20 mM HEPES, 150 mM NaCl, 0.5 mM TCEP, 5% (v/v) glycerol, pH 7.5, concentrated, flash frozen with liquid nitrogen and stored at -80 °C.

**GACG-ubiquitin** A single colony of SoluBL21™(DE3) competent cells (Genlantis) encoding for $His_6$-TEV-GACG-ubiquitin (*4*) was grown overnight in lysogeny broth (LB) supplemented with kanamycin (50 µg $mL^{-1}$) at 37 °C with shaking. The overnight culture was diluted 1:100 in 6 L of LB supplemented with kanamycin (50 µg $mL^{-1}$) and grown to an optical density (OD600) of ~0.75. Protein expression was induced for 4 hours at 37 °C with 1 mM isopropyl β-d-1-thiogalactopyranoside (IPTG). The cells were harvested by centrifugation and frozen at -80 °C as pellets until further purification. The cells were thawed, resuspended in 20 mM HEPES, 500 mM NaCl, 0.5 mM TCEP, pH 7.5, supplemented with 5 mM $MgCl_2$, 1 µg /mL DNAse I and 1X EDTA-free Roche protease inhibitor cocktail, and lysed by cell disruption at 30 kpsi using as One Shot Cell Disruptor (Constant Systems) operating at 4 °C. Cellular debris was removed by centrifugation. $His_6$-TEV-GACG-ubiquitin was purified on a HisTrap™ FF Ni NTA affinity column (Cytiva) and eluted with an imidazole concentration gradient from 0 to 500 mM. The protein was dialysed into 50 mM HEPES, 150 mM NaCl, 0.5 mM TCEP, pH 7.5, incubated with TEV protease, and flowed through a HisTrap FF Ni NTA affinity column. GACG-Ubiquitin was finally purified by size exclusion chromatography on a HiLoad™ 16/600 Superdex™ 75 pg column (Cytiva) in 50 mM Tris-HCl, 150 mM NaCl, 0.5 mM TCEP, pH 7.5, concentrated, flash frozen with liquid nitrogen and stored at -80 °C.

**$^{15}$N-labelled ubiquitin** was prepared using N-terminal 6xHis-TEV on human wild-type Ub residues 1-76, refered to as 6xHis-TEV-Ub. The construct transformed into *Escherichia coli* BL21(DE3) Rosetta II (Novagen) and $^{15}$N-6xHis-TEV-Ub was expressed in autoinducing NPS buffer with $^{15}$N-ammonium chloride (Cambridge Isotope Laboratories) as the only source of nitrogen following (*5*). 2 L of $^{15}$N-NPS autoinducing media was grown at 37 °C for 20 hours, the cells were harvested, resuspended in buffer A (20 mM sodium phosphate, 500 mM NaCl, 50 mM imidazole, 1 mM TCEP, pH 7.4), and lysed at 20 kpsi using as One Shot Cell Disruptor (Constant Systems) operating at 4 °C. The lysate was cleared by centrifugation at 19,000 rpm for 30 min, passed through a 0.45-μm syringe filter, loaded on to an equilibrated 5 mL His-Trap HP column, washed for 30 cv, and eluted with 100% buffer B (20 mM sodium phosphate, 500 mM NaCl, 350 mM imidazole, 1 mM TCEP, pH 7.4). This was followed by dialysis in PBS buffer with TEV protease overnight at 18 °C. The reaction was concentrated and loaded on a 16/60 Superdex 75 column (GE Life Sciences) in PBS. The expected peak for $^{15}$N-Ub was observed around 0.76 CV and its purity was confirmed with SDS-PAGE. Fractions containing pure $^{15}$N-Ub were concentrated using a 3,500 Da centrifugal filter unit (Merck-Millipore) and exchanged into NMR buffer (20 mM sodium phosphate, 150 mM NaCl, 0.5 mM TCEP, pH 7.0).

**Brd4$^{BD2}$** (residues 333-460) was expressed and purified as previously described (*6*). In short, single colonies of cDNA transformed into *E. coli* BL21(DE3) cells were grown overnight in lysogeny broth (LB) supplemented with kanamycin (50 μg mL$^{-1}$) at 37 °C with shaking. The overnight culture was diluted 1:100 in LB supplemented with kanamycin (50 μg mL$^{-1}$) and grown to an optical density (OD600) of ~0.6. Protein expression was induced overnight at 18 °C with isopropyl β-d-1-thiogalactopyranoside (0.3 mM). The cells were harvested by centrifugation and frozen at -80 °C as pellets until further purification. The bacterial pellets were resuspended in buffer and lysed by cell disruption using as One Shot Cell Disruptor (Constant Systems) operating at 4 °C. Cellular debris was removed by centrifugation. The $His_6$-tagged protein was purified on a HisTrap™ FF Ni NTA affinity column (Cytiva) and eluted with an imidazole concentration gradient of 0 to 500 mM. The protein was dialysed into a low concentration imidazole buffer, incubated with TEV protease, and flowed through a HisTrap FF Ni NTA affinity column. Brd4$^{BD2}$ was finally purified by size exclusion chromatography on a HiLoad™ 16/600 Superdex™ 75 pg column (Cytiva) in 20 mM HEPES, pH 7.5, 150 mM sodium chloride and 0.5 mM TCEP, concentrated, flash frozen with liquid nitrogen and stored at -80 °C.

**$^{15}$N-labelled Brd4$^{BD2}$** was obtained as a His-GST-TEV-Brd4(349-459) pGEX4T1 construct. Expression and purification were carried as $^{15}$N-6xHis-TEV-Ub (above). Following the 16/60

Superdex 75 column (GE Life Sciences) step pure fractions of $^{15}$N-Brd4$^{BD2}$ were exchanged into NMR buffer using a 3,500 Da centrifugal filter unit (Merck-Millipore).

**Ub$^{K48C,G76S}$-Brd4$^{BD2}$** variant for thiol crosslinking was prepared from a N-terminal 6xHis-TEV-Ub$^{K48C,G76S}$-Brd4(349-459) "Ub$^{K48C,G76S}$-Brd4$^{BD2}$" construct in pRSF-Duet1 in *Escherichia coli* BL21(DE3) Rosetta II (Novagen). Cultures of lysogeny broth (LB) supplemented with kanamycin (50 µg mL$^{-1}$) were grown at 37 °C until optical density (OD600) of ~0.8, the temperature was lowered 16 °C and expressed with 0.5 mM isopropyl β-d-1-thiogalactopyranoside (IPTG) overnight. Cells were harvested by centrifugation, resuspended in IMAC buffer A (20 mM sodium phosphate, 500 mM NaCl, 50 mM imidazole, 1 mM TCEP, pH 7.4), supplemented with 5 mM $MgCl_2$, and 1 µg/mL DNAse I. Cells were lysed at 20 kpsi using as One Shot Cell Disruptor (Constant Systems) operating at 4 °C. The lysate was cleared by centrifugation at 19,000 rpm for 30 min, passed through a 0.45-µm syringe filter, loaded on to an equilibrated 5 mL His-Trap HP column, washed for 25 cv in IMAC Buffer A, and eluted with 100% IMAC buffer B (20 mM sodium phosphate, 500 mM NaCl, 350 mM imidazole, 1 mM TCEP, pH 7.4). This was followed by dialysis in PBS buffer with TEV protease overnight at 18 °C. The reaction passed through a fresh 10 mL His-Trap HP column to remove TEV protease and uncleaved protein. This was concentrated using a 10,000 Da MWCO centrifugal filter unit (Merck-Millipore) and loaded on 16/60 Superdex 75 column (GE Life Sciences) in cross-linking buffer (20 mM HEPES, 150 mM NaCl, 0.5 mM TCEP, pH 7.0). The purity of Ub$^{K48C,G76S}$-Brd4$^{BD2}$ was confirmed by SDS-PAGE and the protein was stored at -80 °C until use.

**UBE2R1(C93K)** was prepared as a His-SUMO fusion of the full-length human UBE2R1 in pET28b. Expression was carried out in *Escherichia coli* BL21(DE3) Rosetta II (Novagen) cells growin in lysogeny broth (LB) supplemented with kanamycin (50 µg mL$^{-1}$). Cells were grown to optical density (OD600) of ~0.8 at 37 °C, the temperature was adjusted to 16 °C, and induced with 0.35 mM isopropyl β-d-1-thiogalactopyranoside (IPTG) overnight. Cells were harvested by centrifugation, resuspended in IMAC buffer A (20 mM sodium phosphate, 500 mM NaCl, 50 mM imidazole, 1 mM TCEP, pH 7.4), supplemented with 5 mM $MgCl_2$, and 1 µg/mL DNAse I. Cells were lysed at 20 kpsi using as One Shot Cell Disruptor (Constant Systems) operating at 4 °C. The lysate was cleared by centrifugation at 19,000 rpm for 30 min, passed through a 0.45-µm syringe filter, loaded on to an equilibrated 5 mL His-Trap HP column, washed for 25 CV in IMAC Buffer A, and eluted with 100% IMAC buffer B (20 mM sodium phosphate, 500 mM NaCl, 350 mM imidazole, 1 mM TCEP, pH 7.4). This was followed by dialysis in 20 mM HEPES, 150 mM NaCl, 1 mM TCEP, pH 7.5 buffer with 2 µM ULP1 overnight at 4 °C. The reaction passed through a fresh 10 mL His-Trap HP column to remove ULP1 protease and the His-SUMO tag. This was concentrated using a 10,000 Da MWCO centrifugal filter unit (Merck-Millipore) and loaded on 16/60 Superdex 75 column (GE Life Sciences) in 20 mM HEPES, 150 mM NaCl, 1 mM TCEP, pH 7.5). The purity of Ube2R1$^{C93K}$ was confirmed by SDS-PAGE and the protein was concentrated to a 1 mM stock, flash frozen and stored at -80 °C until use. The Ube2R1$^{C92K,S138C, C191S,C223S}$ variant used for chemical cross-linking was prepared following the same method, with the only exception being the buffer for final 16/60 Superdex 75 column (GE Life Sciences) was 20 mM HEPES, 150 mM NaCl, 0.5 mM TCEP, pH 7.0

**Ubiquitin-activating enzyme 1 (UBA1)** was expressed following (*7*) to obtain the full length human UBA1 with C-terminal 6xHis in pRSF-Duet1. *Escherichia coli* BL21(DE3) Rosetta II

(Novagen) cells were grown at 37 °C in lysogeny broth (LB) supplemented with kanamycin (50 μg $mL^{-1}$) and 1 mM $MgSO_4$ until optical density (OD600) of ~0.8. The temperature was adjusted to 16 °C, isopropyl β-d-1-thiogalactopyranoside (IPTG) was added to 0.2 mM, and induction proceeded overnight. Cells were harvested by centrifugation, resuspended in IMAC buffer A (20 mM sodium phosphate, 500 mM NaCl, 50 mM imidazole, 1 mM TCEP, pH 7.4), supplemented 5 mM $MgCl_2$, and 1 μg/mL DNAse I. Lysis was performed at 20 kpsi using as One Shot Cell Disruptor (Constant Systems) operating at 4 °C. The lysate was cleared by centrifugation at 19,000 rpm for 30 min, passed through a 0.45-μm syringe filter, loaded on to an equilibrated 5 ml His-Trap HP column, washed for 25 cv in IMAC Buffer A, and eluted with 100% IMAC buffer B (20 mM sodium phosphate, 500 mM NaCl, 350 mM imidazole, 1 mM TCEP, pH 7.4). The elution was concentrated to ~2 mL using a 30,000 Da MWCO centrifugal filter unit (Merck-Millipore) and diluted 1:100 in anion buffer A (50 mM TRIS, 5 mM 2-mercaptoethanol, pH 8.0) and loaded on an equilibrated 5 ml Q-HP column (Cytiva). A 50% gradient over 40 cv with anion buffer B (50 mM TRIS, 1M NaCl, 5 mM 2-mercaptoethanol, pH 8.0) was applied and SDS-PAGE showed UBA1 eluted in the first major peak at ~18% buffer B. These fractions were concentrated on loaded on to a final 16/60 Superdex 200 column (Cytiva) equilibrated in 20 mM HEPES, 150 mM NaCl, 1 mM TCEP, pH 8.0. Pure UBA1 was concentrated to 100 μM and stored at -80 °C until use.

**Ubiquitin(E34C) and Ube2R1(C93K) (His construct).** Both were prepared using an N-terminal $His_6$-TEV cleavable tag. Single colonies transformed into *E. coli* BL21 (DE3) were grown overnight in lysogeny broth (LB) supplemented with kanamycin (50 μg/ml) at 37 °C with shaking at 220 rpm. The overnight culture was diluted 1:100 in 0.5 L LB supplemented with kanamycin (50 μg/ml) and grown to an optical density (OD600) between 0.6-0.8 at 37 °C with shaking at 220 rpm. Isopropyl β-d-1-thiogalactopyranoside (IPTG) was added to a final concentration of 0.1 mM to induce protein expression and cultures were incubated at 18°C with shaking at 220 rpm for 18 hours. Bacterial cells were pelleted by centrifugation (6,500 RPM / 20 min / 4 °C) and resuspended in 50 mM Tris, 150 mM NaCl, 10 mM imidazole, complete protease inhibitor cocktail (EDTA-free, Roche), pH 7.5. The cell suspension was either flash-frozen in liquid nitrogen and stored at -80 °C until further use, or purification was carried out straight away. The bacterial cells were lysed by cell disruption (Avestin) and centrifuged (27,200 g / 45 min / 4 °C) to remove any insoluble material. The supernatant was filtered through a 0.2 μM filter and loaded onto a Ni-NTA agarose (Qiagen) column (1 mL of beads per 0.5 litre of bacterial culture) pre-equilibrated with 50 mM Tris, 150 mM NaCl, 10 mM imidazole, pH 7.5. The column was washed successively with 50 mM Tris, 150 mM NaCl, 10 mM imidazole, pH 7.5 (~8 column volumes) and 50 mM Tris, 150 mM NaCl, 30 mM imidazole, pH 7.5 (~6-8 column volumes) and eluted with 50 mM Tris, 150 mM NaCl, 150 mM imidazole, 0.5 mM TCEP, pH 7.5. Fractions containing $His_6$-Ub/Ube2R1 C93K were pooled, TEV protease was added (1 mg TEV:50 mg (Ub/Ube2R1)) and dialyzed for 18 hours at 4°C against 50 mM Tris, 150 mM NaCl, 0.5 mM TCEP, pH 7.5. Once most of the protein was cleaved, imidazole was added to the final concentration of 10 mM and passed through the Ni-NTA agarose column pre-equilibrated with 50 mM Tris, 150 mM NaCl, 10 mM imidazole, 0.5 mM TCEP, pH 7.5. The flow-through fraction was collected, concentrated to between 5-10 mg/mL using a Vivaspin centrifugal concentrator (Sartorius) (3 kDa molecular weight cut-off, 3,000 g, 4 °C), flash-frozen in liquid nitrogen and stored at -80 °C.

Ubiquitin labelling with maleimide Alexa Fluor 488

AlexaFluor488 C5 maleimide dye was dissolved to 5 mM in anhydrous DMSO. Ubiquitin cysteine mutant recombinant protein was buffer exchange into 20 mM HEPES, 150 mM NaCl, pH 7.0 using a 10/300 GL Superdex 75 Increase prepacked column (Cytiva). The fractions containing the protein were combined, concentrated to 100 μM and labelled with a 5 times molar excess of fluorescent dye for 2 hours at room temperature. The excess dye was removed by gel filtration on a 10/300 GL Superdex 75 Increase prepacked column (Cytiva), eluting in 20 mM HEPES, 150 mM NaCl, 0.5 mM TCEP, pH 7.5. The dye and subsequently dye-labelled protein were protected from light throughout, flash frozen and stored at -80 °C.

Brd4$^{BD2}$ labelling with maleimide Alexa Fluor 647

AlexaFluor647 C5 maleimide dye was dissolved to 5 mM in anhydrous DMSO. Brd4$^{BD2}$ recombinant protein was buffer exchange into 20 mM HEPES, 150 mM NaCl, pH 7.0 using 0.5 mL 7K MWCO Zeba™ Spin desalting columns (Thermo Fisher Scientific). The protein was concentrated to 100 μM and labelled with a 5 times molar excess of fluorescent dye for 2 hours at room temperature. The excess dye was removed using a CentriPure P2 desalting column (Generon), eluting in 20 mM HEPES, 150 mM NaCl, 0.5 mM TCEP, pH 7.5. The dye and subsequently dye-labelled protein were protected from light throughout, flash frozen and stored at -80 °C.

Preparative scale ubiquitin loading assay with Ube2R1(C93K) and Ube2R1(C93K, S138C, C191S, C223S)

UBE2R1(C93K) and UBE2R1(C93K,S138C,C191S,C223S) (5 μM) were incubated with 0.5 μM Ub E1 and 50 μM Ub for 18 hrs at 37 °C in 50 mM TRIS pH 9.5, 50 mM NaCl, 5 mM ATP, 10 mM MgCl2, and 1 mM BME. The Ub-UBE2R1 conjugate was purified by gel filtration on a Superdex 75 pg 16/600 in 50 mM TRIS, 150 NaCl, 0.5 mM TCEP, pH 7.5, concentrated, flash frozen with liquid nitrogen and stored at -80 °C. Samples were resolved by SDS-PAGE and unmodified/modified Ube2R1 was visualized by Coomassie staining.

BMOE crosslinking of Ube2R1(C93K,S138C,C191S,C223S)-Ub with His$_6$-TEV-Ub(G76S,K48C)-Brd4$^{BD2}$

125 nmol of Ube2R1$^{C93K,S138C,C191S,C223S}$-Ub was incubated with 15 mM TCEP for 20 min at ambient temperature and desalted on HiTrap desalting column (Cytiva) in cross-linking buffer (20 mM HEPES, 150 mM NaCl, pH 7.0). Fractions were pooled and immediately a 15-fold molar excess of bismaleimidoethane (BMOE) was added from a 30 mM DMSO stock. 625 nmol of His-TEV-Ub$^{K48C,G76S}$-Brd4$^{BD2}$ was treated with 15 mM TCEP and desalted in cross-linking buffer. Following a 60 min incubation, BMOE Ube2R1$^{C93K,S138C,C191S,C223S}$-Ub was directly desalted into the 6xHis-TEV-Ub$^{K48C,G76S}$-Brd4$^{BD2}$ sample, and cross-linking was allowed to occur overnight at 4 °C. The reaction was quenched with 2-mercaptoethanol (10 mM). The desired cross-linked product, 6xHis-TEV-Ub$^{G76S,K48C}$-Brd4$^{BD2}$-BMOE- Ube2R1$^{C93K,S138C,C191S,C223S}$-Ub, was confirmed with SDS-PAGE. The complex was then bound to 1.5 mL Ni NTA resin (Cytiva), washed with 20 mM imidazole and eluted with 500 mM imidazole. The complex was then buffer exchanged into 50 mM Tris-HCl, 0.5 mM TCEP, pH 7.5 and further purified by anion exchange

chromatography on a 5 mL HiTrap Q HP prepacked column (Cytiva). The complex was eluted on a gradient of buffer A (50 mM Tris-HCl, 0.5 mM TCEP, pH 7.5) to buffer B (50 mM Tris-HCl, 500 mM NaCl, 0.5 mM TCEP, pH 7.5). The fractions containing pure crosslinked 6xHis-TEV-$Ub^{G76S,K48C}$-$Brd4^{BD2}$-BMOE- $Ube2R1^{C93K,S138C,C191S,C223S}$-Ub were concentrated, flash frozen with liquid nitrogen and stored at -80 °C.

*In vitro* ubiquitination assay

The E1 Ube1 (150 nM), the UBE2D2 or UBE2R1 (5 μM), $Brd4^{BD2}$ (5 μM), NEDD8-$CRL2^{VHL}$ (150 nM), and MZ1 (5 μM) were mixed with ubiquitin (100 μM) and incubated in 20 mM HEPES, 150 mM NaCl, 0.5 mM TCEP, 5 mM $MgCl_2$, pH = 7.5 for 5 minutes at room temperature. ATP (3 mM) was added and the mixture was incubated at room temperature. The reaction was quenched with reducing SDS sample buffer. The protein species were resolved by SDS-PAGE on a 12% Bis-Tris NuPAGE gel using MOPS running buffer, run at 200 V for 40 minutes. The gel was stained with Coomassie Instant Blue.

Ubiquitin-directed photoreactive probe (UDPRP) production

The cysteine reactive photocrosslinker, N-maleimido diazirine, was synthesized as described in (*8*). Stocks of N-Maleimido diazirine were dissolved to 10 mM using anhydrous DMSO and stored in 50 μL aliquots at -20 °C for months. Ubiquitin E34C was buffer exchanged into degassed 50 mM Tris, 150 mM NaCl, pH 7.0 using a Centri Pure Zetadex-25 gel filtration column (Generon) and labelled with N-Maleimido diazirine at room temperature for 2 hours using a 5 times molar excess of N-Maleimido diazirine. Excess N-Maleimido diazirine was removed using a Centri Pure Zetadex-25 gel filtration column and 50 mM Tris, 150 mM NaCl, and 0.5 mM TCEP, pH 7.5 as running buffer. All buffers were degassed, all dye labelling reactions and dye labelled proteins were protected from light and all products were analyzed by intact LC-MS. Photocrosslinker-labelled ubiquitin was conjugated to the active site of Ube2R1 C93K in a ubiquitin-loading assay (described above). The stable photocrosslinker labelled Ubiquitin-Ube2R1 conjugate was purified by gel filtration on a Superdex 75 pg 16/600 column in, 20 mM HEPES, 150 NaCl, 0.5 mM TCEP, pH 7.5, concentrated, flash frozen with liquid nitrogen and stored at -80 °C.

Sortase-mediated biotinylation of UBE2R1

$His_6$-Sortase was used to conjugate the peptide Biotin-LPTGG (synthesized by Peptide2) to the N-terminal glycine residue of UBE2R1. 1 μM $His_6$-Sortase, 20 μM Ube2R1, 200 μM Biotin-LPTGG peptide were incubated at 37°C for 15 minutes in 50mM TRIS, 150 mM NaCl. Imidazole was added to 10 mM and the $His_6$-Sortase was removed immediately by Ni-NTA chromatography. Excess peptide was removed by Centri Pure Zetadex-25 gel filtration column using 50 mM Tris, 150 mM NaCl, and 0.5 mM TCEP, pH 7.5 as running buffer. Biotinylated-Ube2R1 was concentrated to between 5-10 mg/mL using a Vivaspin centrifugal concentrator (Sartorius) (10 kDa molecular weight cut-off, 3,000 g, 4 °C), flash-frozen in liquid nitrogen and stored at -80 °C.

UDPRP crosslinking assay

Photo-cross linking reactions (20 μL) were performed with UDPRP (10 μM) and (NEDD8)-CRL2$^{VHL}$ (3 μM) in an 18-well glass-bottom plate (ibidi) in reaction buffer (20 mM HEPES, pH 7.5, 150 mM NaCl, 1 mM TCEP) for visualisation by western blot. Higher concentrations of both UDPRP (20 μM) and the (NEDD8)-CRL2$^{VHL}$ (5 μM) were used when visualising the crosslinked product by Coomassie staining. In both cases, samples were divided into two portions. One portion was irradiated at 365 nm, the plate was kept on an ice-cold metal block located 10 cm away from a handled UV lamp (BLE-8T365, Spectroline), for 5 min and the other portion was preserved in the dark on ice. Samples were resolved by SDS-PAGE and photo-crosslinked products were visualized by Coomassie staining or immunoblotting.

Immunoblotting

Primary antibodies: Rbx1 anti-rabbit, 1:10,000 (Cell signaling, #11922). Secondary antibodies and dyes: IRDye 680LT Donkey anti-Rabbit and IRDye® 800CW Streptavidin, both 1:5000 (Licor). Samples were diluted 1:3 in 2X reducing LDS sample buffer (NuPage) and proteins were separated by SDS-Page on a 12% polyacrylamide Bis-Tris gel (NuPage) in 1% MES buffer. Proteins were transferred to Nitrocellulose membrane using iBlot 2 Gel Transfer Device (Invitrogen). Membranes were blocked for 1 hour in 5% milk in PBS-T and incubated overnight with primary antibodies, then for 1 hour with secondary antibodies and dyes (1:5000) before imaging on the Licor (Odyssey DLx).

*In vitro* stUbl-mediated ubiquitination assay to verify the activity of the photocrosslinker labelled ubiquitin E34C

The activity of the photocrosslinker-labelled ubiquitin E34C was compared to wild-type ubiquitin in a stUbl assay containing the SUMO-targeted E3 ligase, RNF4, Ubch5a and UbA1. The assay and protein purification methods are described in detail in (*4*).

Preparation of samples for cryo-electron microscopy

Frozen stocks of recombinant protein and PROTAC were thawed and kept at 4 °C throughout the sample preparation process. For the 'open' non-crosslinked Brd4$^{BD2}$-MZ1-(NEDD8)-CRL2$^{VHL}$-UBE2R1-Ub structure, 6 μM NEDD8-CRL2$^{VHL}$ (1 equivalent), MZ1 (1.5 equivalents), Brd4$^{BD2}$ (1.5 equivalents) and UBE2R1(C93K)-Ub conjugate (1.5 equivalents) were incubated for 10 minutes at 4 °C. For the 'closed' crosslinked (NEDD8)-CRL2$^{VHL}$-MZ1-Brd4$^{BD2}$-Ub(G76S,K48C)-UBE2R1(C93K, S138C, C191S, C223S)-Ub structure, 6 μM NEDD8-CRL2$^{VHL}$ (1 equivalent), MZ1 (1.5 equivalents) and Brd4$^{BD2}$-Ub(G76S,K48C)-UBE2R1(C93K, S138C, C191S, C223S)-Ub (1.5 equivalents) were incubated for 10 minutes at 4 °C. The complexes were desalted on a 0.5 mL 7K MWCO Zeba™ Spin Desalting Column in 20 mM HEPES, 150 mM NaCl, 0.5 mM TCEP, pH 7.5. Quantifoil R1.2/1.3 holey carbon copper 400 mesh grids were glow discharged for 60 seconds at 35 mA using a Quorum SC7620. 3.5 μL of protein at 4 μM was applied to the cryo-EM grids and was vitrified in liquid ethane on a Vitrobot Mark IV (Thermo Fisher Scientific) at 4 °C and 100% humidity (wait time = 10 s, blot force = 4, blot time = 3.5 s, blot total = 1, drain time = 0 s).

Cryo-electron microscopy data acquisition

For the 'open' non-crosslinked Brd4$^{BD2}$-MZ1-(NEDD8)-CRL2$^{VHL}$-UBE2R1-Ub structure, cryo-EM data were collected on Glacios transmission electron microscope (Thermo Fisher Scientific) operating at 200 keV. Micrographs were acquired using a Falcon4i direct electron detector (Thermo Fisher Scientific), operated in electron counting mode. A total electron exposure of 26 $e^-/Å^2$ was applied. EPU (Thermo Fisher Scientific, version 3.0) was used to collect micrographs at 190,000k nominal magnification (0.74 Å/pixel at the specimen level) with a nominal defocus range of -1.7 to -3.2 µm. Stage shifts with aberration-free image shift (AFIS) mode was used to centre multiple foil holes and image shift was used to acquire high magnification images in the centre of each targeted hole. 4,961 movies were collected in EER format.
For the 'closed' crosslinked (NEDD8)-CRL2$^{VHL}$-MZ1-Brd4$^{BD2}$-Ub(G76S,K48C)-UBE2R1(C93K, S138C, C191S, C223S)-Ub structure, cryo-EM data were collected on Krios transmission electron microscope (Thermo Fisher Scientific) operating at 300 keV. Micrographs were acquired using a K3 direct electron detector (Gatan), operated in electron counting mode. A total electron exposure of 38 $e^-/Å^2$ was applied. EPU (Thermo Fisher Scientific, version 3.0) was used to collect micrographs at 105,000k nominal magnification (0.825 Å/pixel at the specimen level) with a nominal defocus range of -1.2 to -3.0 µm. Stage shifts with aberration-free image shift (AFIS) mode was used to centre multiple foil holes and image shift was used to acquire 8 high magnification images following a template around the edge of each targeted hole. 14,047 movies were collected in MRC format.

Cryo-electron microscopy image analysis and model building

Image processing pipelines are described in Fig. S3 and Fig. S10. Cryo-EM movies were imported into CryoSPARC v.4.4.0-v4.4.1 (*9*) for patch motion correction, patch CTF estimation and manual curation. For the 'open' non-crosslinked Brd4$^{BD2}$-MZ1-(NEDD8)-CRL2$^{VHL}$-UBE2R1-Ub, manual picking was performed on 84 micrographs which were used for template picking and 2D classification. Good templates were used for Topaz training and particle picking (*10*). 405,567 particles were extracted with a 512 pixel box size (2x binning). Classification was achieved using *ab initio* reconstruction into 4 classes. Particles for the best class were submitted to refinement, re-extracted with the full 512 pixel box size and submitted to non-uniform refinement in CryoSPARC (*11*). For the closed' crosslinked (NEDD8)-CRL2$^{VHL}$-MZ1-Brd4$^{BD2}$-Ub(G76S,K48C)-UBE2R1(C93K, S138C, C191S, C223S)-Ub structure, particle picking was performed on 4,458 movies with crYOLO using a general model for low-pass filtered images (*12*). Particles were submitted to 2D classification and good templates were used for Topaz training and particle picking (*10*). 748,020 particles were extracted with a 432 pixel box size (3x binning). Classification was achieved 2D classification followed by *ab initio* reconstruction and heterogeneous refinement into 3 classes. The best class was submitted to 3D classification into 3 classes. The particles from the best classes were combined, re-extracted to 480 pixels and submitted to non-uniform refinement CryoSPARC (*11*). Representative 2D classes, orientation diagnostics, local resolution estimations and gold-standard Fourier shell correlation (GSFSC) curves were generated with CryoSPARC (*11*) and are shown in Fig. S3 and Fig. S10. Model building was achieved using atomic models from AlphaFold and PDB entries 5T35, 5N4W, 4AP4 and 6TTU, which were docked into the cryo-EM maps using rigid-body fitting with UCSF

ChimeraX (*13*). The model was refined using ISOLDE (*14*) until reasonable agreement between the model and data were achieved.

Nuclear Magnetic Resonance

Solution NMR data was acquired at 298 K on a Bruker Avance III 600 MHz spectrometer and cryogenic TCI probe. Each protein sample was prepared at a concentration of 150 µM and recorded in buffer 20 mM sodium phosphate, 150 mM NaCl, 0.5 mM TCEP, 5% $D_2O$, pH 7.0. $^{15}N$-$^{1}H$-HSQC spectra were recorded with 128 points in the $^{15}N$ dimension and processed with 256 points using Bruker TopSpin version 3.6. Previous assignment for Ub (*5*) matched well for Ub($^{15}N$) and signals from Ub in Ube2R1$^{C93K}$~Ub($^{15}N$) were able to be assigned. Spectra were analysed and printed using CARA. The chemical shift perturbation (CSP) was calculated according to $CSP = [(\delta_{HA} - \delta_{HB})^2 + ((\delta_{NA} - \delta_{NB})/5)^2]^{1/2}$.

Mass photometry

Interferometric scattering microscopy (*15*) was carried out using the commercially available oneMP (Refeyn). The cryo-EM buffer, 20 mM HEPES, 150 mM NaCl, 0.5 mM TCEP, pH 7.5 was filtered through a 0.22 µm syringe filter, gaskets wells (Grace Bio-labs CW-50R-1.0) along with high precision 24 x 50 mm coverslips (Marienfeld) were prepared at measurement according to (*16*). 10 µl of buffer was used for focusing in regular mode (128 x 34 binned pixels 18.0 $\mu m^2$ detection area) and all data was recorded for 60 seconds using AquireMP software (Refeyn). Calibration standards for mass calibration consisted of conalbumin ($M_r$ 75 000), aldolase ($M_r$ 158 000), ferritin ($M_r$ 440 000), and thyroglobulin ($M_r$ 669 000) were prepared from the gel filtration HMW calibration kit (Cytiva). The various complexes containing CRL$^{VHL}$ were measured in the same buffer in a concentration range of 20-30 nM. All data was processed and analyzed in DiscoverMP (Refeyn).

Intact LC-MS

For all intact LC-MS measurements 0.5 µg protein was diluted in the appropriate buffer and injected in a 20 µL volume. LC-MS was carried out with an Agilent 1200 LC-MS system fitted with a Max-Light Cartridge flow cell coupled to a 6130 Quadrupole spectrometer. An Agilent ZORBAX 300SB-C3 5 µm, 2.1 x 150 mm column was employed unless otherwise stated. Protein UV absorbance was monitored at 214 and 280 nm. MS acquisition was carried out in positive ion mode and total protein masses were calculated by deconvolution within the MS Chemstation software (Agilent Technologies).

Mass spectrometry identification of bromodomain lysine ubiquitination sites

Proteins from the regions of the gel relating to ubiquitin modified Brd4$^{BD2}$ forms were in-gel trypsin digested (*17*), alkylated with chloroacetamide and final peptides were resuspended in 0.1% TFA 0.5% acetic acid before analysis by LC-MS/MS. Samples were analysed on a Thermo scientific Lumos Tribrid mass spectrometer coupled with a Thermo Dionex Ultimate 3000 RSLC HPLC. The Buffers used for HPLC were 0.1% formic acid as buffer A and 80% acetonitrile with 0.08% formic acid as buffer B. Trap column Acclaim pepmap 100 (C185 µM, 100 µM x 2cm) was

used before the main column for sample concentration and clean up. The peptides samples were loaded onto the trap column using loading pump with 3% acetonitrile 0.1% trifluoroacetic acid at flow rate of 5 µL/min. The main column used was EASY-Spray column (C18, 2 µM, 75 µm x 50 cm) with a nano electrospray emitter built in. The flow rate of 300 nL/min was maintained throughout the run. Peptides were separated with a 55 min segmented gradient starting from 3%~35% buffer B over 45 min, 35%~95% buffer B over 47 mins and held for 5 min. The separated peptides were then analysed on Lumos Mass spectrometer. Spray voltage was set to 2 kV, RF lens level was set at 30%, and ion transfer tube temperature was set to 275 °C. The mass spectrometer was operated in data-dependent settings with 3 seconds cycle time. The full scan was performed in the range of 375—1500 m/z at nominal resolution of 120,000 at 200 m/z and AGC set to 400000 with a custom maximum injection time of 50 ms. This was followed by selection of the most intense ions above an intensity threshold of 5000 for higher-energy collision dissociation (HCD) fragmentation, with normalised collision energy set to 30. MS2 scans were acquired for charge states 2 to 7 using an isolation width of 1.6 m/z and a 30 second dynamic exclusion duration. MS2 scans were done using an AGC target set to 50000 and a maximum fill time of 50 ms.
Data analysis used MaxQuant version 2.4.0.0 (*18*). Default settings were used with a few exceptions. A database of all the recombinant proteins included in the *in vitro* ubiquitination assay was used. Digestion was set to Trypsin with a maximum of 3 missed cleavages. Match between runs was not enabled. Oxidation (M), Acetyl (Protein N-term) and GlyGly (K) were included as variable modifications, with a maximum of 3 per peptide allowed. Carbamidomethyl (C) was included as a fixed modification. Only peptides of maximum mass 8,000 Da were considered. Protein and peptide level FDR was set to 1% but no FDR filtering was applied to identified sites. Manual MS/MS sequence validation was used to verify GlyGly (K) peptide identifications and only peptides with an Andromeda score >100, localization probability >0.75 and a mass error <1 ppm were considered.

Verification of the ubiquitin loading assay with Ube2R1(C93K) and identification of UDPRP crosslinks

Proteins from the regions of the gel relating to modified and unmodified Ube2R1 or UDPRP photo-crosslinked products were in-gel trypsin digested (*17*). alkylated with chloroacetamide and final peptides were resuspended in 0.1% TFA 0.5% acetic acid before analysis by LC-MS/MS. This was performed using a Q Exactive mass spectrometer (Thermo Scientific) coupled to an EASY-nLC 1000 liquid chromatography system (Thermo Scientific), using an EASY-Spray ion source (Thermo Scientific) running a 75 µm x 500 mm EASY-Spray column at 45ºC. Two MS runs (of 60 and 150 minutes) were prepared using approximately 15% total peptide sample each. To boost sensitivity a top 3 data-dependent method was applied employing a full scan (m/z 300–1800) with resolution R = 70,000 at m/z 200 (after accumulation to a target value of 1,000,000 ions with maximum injection time of 20 ms). For the 60 minute gradient the 3 most intense ions were fragmented by HCD and measured with a resolution of R = 70,000 (60 minute run) or 35,000 (150 minute run) at m/z 200 (target value of 1,000,000 ions and maximum injection time of 500 ms) and intensity threshold of $2.1 \times 10^4$. Peptide match was set to 'preferred'. Ions were ignored if they had unassigned charge state 1, 8 or >8 and a 10 second (60 minute run) or 25 second (150 minute run) dynamic exclusion list was applied.

**Verification of the ubiquitin loading assay with Ube2R1(C93K).** Data analysis used MaxQuant version 1.6.1.0 (*18*). Default settings were used with a few exceptions. A database of all the recombinant proteins included in the ubiquitin-loading assay was used. Digestion was set to Trypsin/P (ignoring lysines and arginines N-terminal to prolines) with a maximum of 3 missed cleavages. Match between runs was not enabled. Oxidation (M), Acetyl (Protein N-term) and GlyGly (K) were included as variable modifications, with a maximum of 4 per peptide allowed. Carbamidomethyl (C) was included as a fixed modification. Only peptides of maximum mass 8000 Da were considered. Protein and peptide level FDR was set to 1% but no FDR filtering was applied to identified sites. Manual MS/MS sequence validation was used to verify GlyGly (K) peptide identifications and only peptides with an Andromeda score >100, localization probability>0.75 and a mass error <1ppm were considered.

**Identification of UDPRP crosslinks.** Data analysis was performed with MaxQuant version 2.4.0.0 (*18*). Default settings were used with a few exceptions. A database of all the recombinant proteins included in the UDPRP crosslinking assay was used. Digestion was set to Trypsin/P (ignoring lysines and arginines N-terminal to prolines) with a maximum of 8 missed cleavages. Match between runs was enabled. Prior to running the search, a new crosslinker was added to MaxQuant configurations describing, N-Maleimido-diazirine and was named 'NMD'. Linked composition H(9)O(2)C(8)N, mass 151.0633285383 Da. Hydrolyzed composition H(9)O(2)C(8)N(3), mass 179.0694765487. Specificity 1 was C, position in peptide 1 was set to anywhere. Protein N-term 1 and C-term 1 were selected. Specificity 2 set for any amino acid (ACDEFGHIKLMNPQRSTVWY), position in peptide 2 was anywhere, protein N-term 2 and protein C-term 2 were selected. To search for crosslinked peptides the crosslinker NMD (non-cleavable) was selected, minimum length for a paired sequence was set to 3 and the maximum peptide mass 12,000 Da. The minimum peptide length for unspecified peptide search was 8 and maximum 25. The search included both intra-protein and inter-protein crosslinked peptides. Oxidation (M), Acetyl (Protein N-term) and Carbamidomethyl (C) were included as variable modifications, with a maximum of 5 per peptide allowed. First search was performed with Oxidation (M) and Acetyl (Protein N-term). Protein and crosslinked peptide level FDR was set to 1% but no FDR filtering was applied to identified sites. Manual MS/MS sequence validation was used to verify inter-protein NMD crosslinked peptide identifications and only peptides with an Andromeda score >95, localization probability >0.75 and a mass error <1 ppm were considered.

Multiple sequence alignments

Coding sequences for human BRD2 (P25440), BRD3(Q15059), BRD4 (O60885), and BRDT (Q58F21) were taken from Uniprot. Indivisual FASTA files were generated using the boundries of the BD1 and BD2. Multiple sequence alignments were carried out in Jalview (*19*) using MUSCLE (*20*). Alignments for figures were exported directly from Jalview.

Synthesis of 1-(2-(3-methyl-3H-diazirin-3-yl)ethyl)-1H-pyrrole-2,5-dione

All chemicals unless otherwise stated, were commercially available and used without further purification. Commercially available dry solvents were used. Flash column chromatography was performed using a Teledyne Isco Combiflash Rf with prepacked Redisep

RF Normal phase disposable columns. NMR Spectra were recorded on a Bruker 400 MHz or 500 MHz as specified. $^{13}C$ spectra were $^{1}H$ decoupled. Chemical shifts ($\delta$) are reported in ppm and referenced to the residual solvent signals: $^{1}H$ NMR $\delta$ (ppm) = 7.26 ($CDCl_3$), $^{13}C$ NMR $\delta$ (ppm) = 77 ($CDCl_3$). Signal splitting patterns are described as singlet (s) and triplet (t). Coupling constants (*J*) are measured in Hertz (Hz).

Diisopropyl azodicarboxylate (360 µL, 1.8 mmol) was added dropwise to a solution of 2-(3-methyldiazirin-3-yl)ethanol (150 mg, 1.5 mmol), triphenylphosphine (432 mg, 1.65 mmol) and maleimide (160 mg, 1.65 mmol) in THF (3 mL) at 0 °C and the reaction mixture was stirred overnight at room temperature. After solvent evaporation, the residue was purified by column chromatography, elution gradient of 0 to 50% of EtOAc in heptane, to afford 1-(2-(3-methyl-3H-diazirin-3-yl)ethyl)-1H-pyrrole-2,5-dione as a colorless oil (139 mg, 52%). $^{1}H$ NMR (500 MHz, $CDCl_3$): 6.74 (2H, s), 3.59 (2H, t, $J$ = 7.1 Hz), 1.62 (2H, t, $J$ = 7.1 Hz), 1.08 (3H, s). $^{13}C$ NMR (100 MHz, $CDCl_3$): 170.4, 134.3, 33.4, 33.2, 23.9, 19.1. NMR spectra (Fig. S14) were in agreement with the published data (*8*).

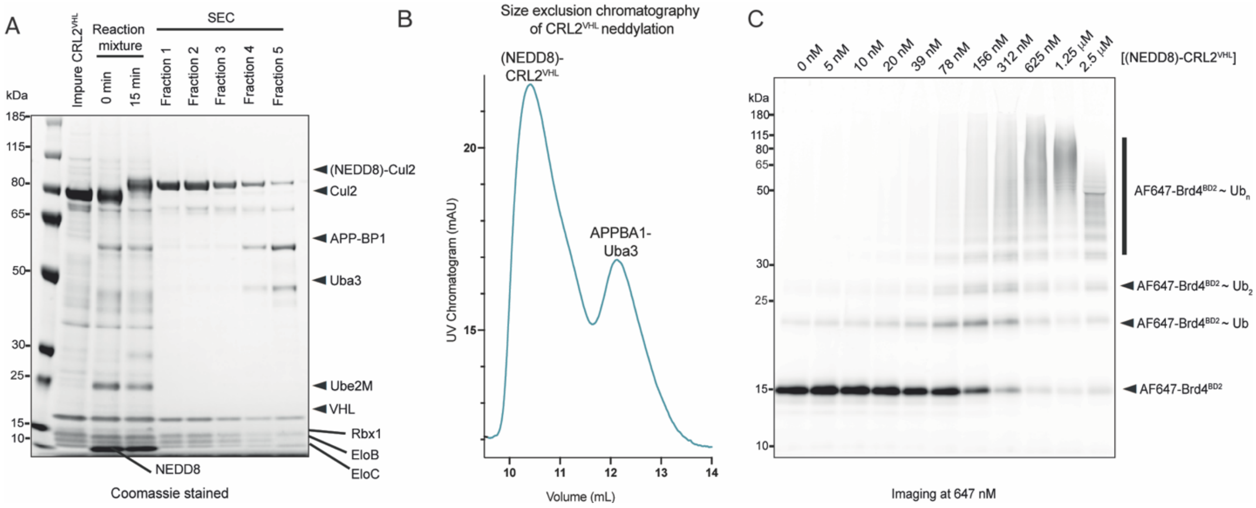

**Fig. S1. (NEDD8)-CRL2$^{VHL}$ purification and *in vitro* ubiquitination assay.** (**A**) SDS-PAGE of CRL2$^{VHL}$ neddylation reaction and (NEDD8)-CRL2$^{VHL}$ purification fractions by size exclusion chromatography. (**B**) Size exclusion chromatography purification UV trace showing elution of (NEDD8)-CRL2$^{VHL}$ and APPBP1-Uba3 (cropped to elution volumes of 9-14 mL). (**C**) SDS-PAGE (imaged at 647 nm) of Alexa Fluor 647-labelled Brd4$^{BD2}$ *in vitro* ubiquitination, demonstrating the enzymatic activity of recombinant (NEDD8)-CRL2$^{VHL}$. The reactant species were mixed to final concentrations of Ube1 (150 nM), UBE2D2 (5 µM), Brd4$^{BD2}$ (4 µM), Alexa Fluor 647 maleimide-labelled Brd4$^{BD2}$ (1 µM), NEDD8-CRL2$^{VHL}$ (0-2.5 mM), MZ1 (5 µM) and ubiquitin (100 µM) and incubated in 20 mM HEPES, 150 mM NaCl, 0.5 mM TCEP, 5 mM $MgCl_2$, pH = 7.5 for 5 minutes at room temperature. The reaction was launched with ATP (3 mM) and the samples were quenched after 30 minutes with SDS sample buffer. The protein species were resolved by SDS-PAGE on a 12% Bis-Tris NuPAGE gel using MOPS running buffer, run at 200 V for 40 minutes. The gel was fluorescently imaged at 647 nm to identify Alexa Fluor 647 maleimide-labelled Brd4$^{BD2}$.

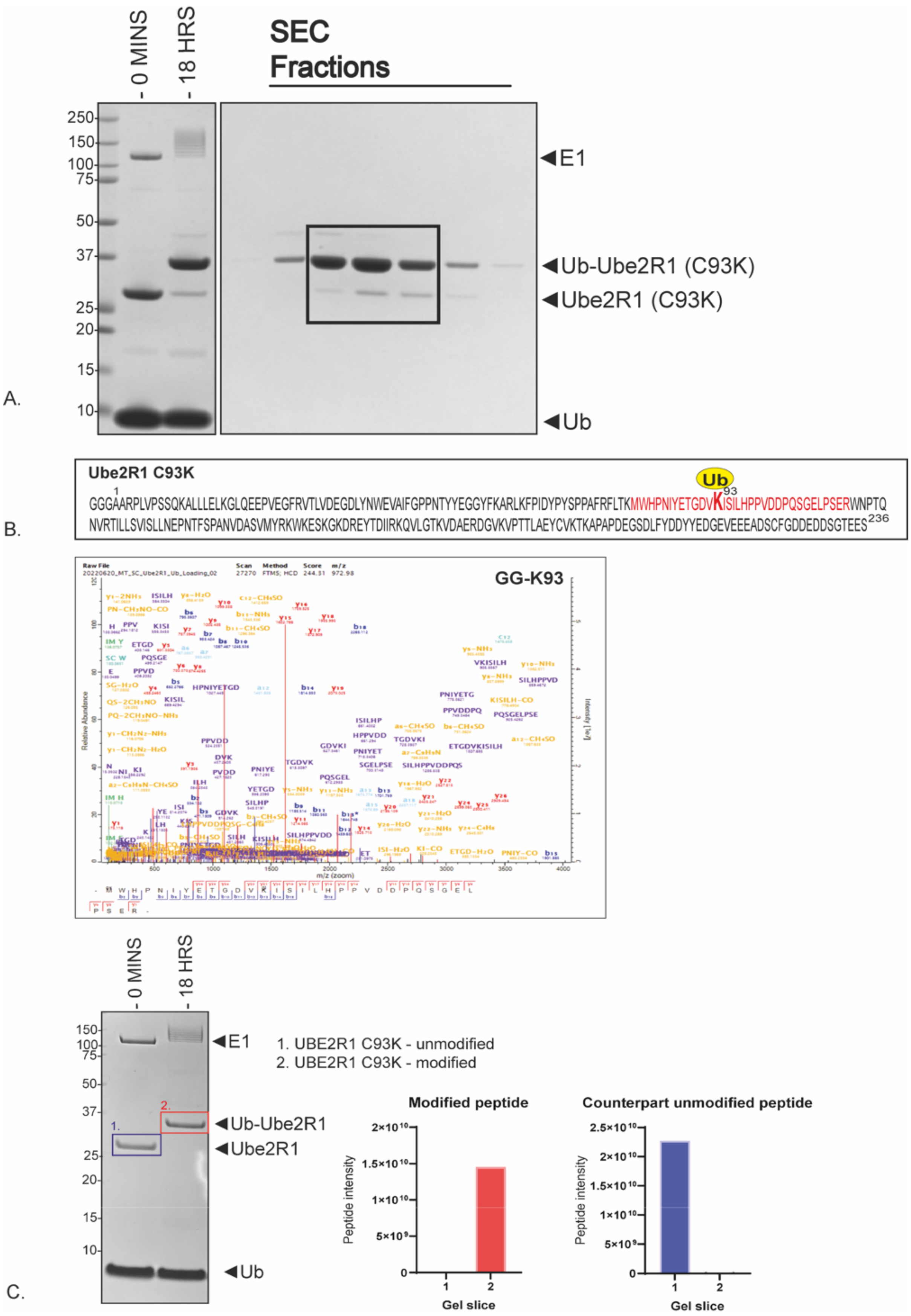

0 MINS
18 HRS
SEC Fractions
250
150
100
75
50
37
25
20
15
10
E1
Ub-Ube2R1 (C93K)
Ube2R1 (C93K)
Ub
A.
Ube2R1 C93K
Ub
1
93
GGGAARPLVPSSQKALLLELKGLQEEPVEGFRVTLVDEGDLYNWEVAIFGPPNTYYEGGYFKARLKFPIDYPYSPPAFRFLTKMWHPNIYETGDVKISILHPPVDDPQSGELPSERWNPTQ
NVRTILLSVISLLNEPNTFSPANVDASVMYRKWKESKGKDREYTDIIRKQVLGTKVDAERDGVKVPTTLAEYCVKTKAPAPDEGSDLFYDDYYEDGEVEEEADSCFGDDEDDSGTEES
236
B.
Raw File
20220620_MT_SC_Ube2R1_Ub_Loading_02
Scan
27270
Method
FTMS; HCD
Score
244.31
m/z
972.98
GG-K93
Relative Abundance
m/z (zoom)
Intensity [e7]
0 MINS
18 HRS
150
100
75
50
37
25
20
15
10
E1
Ub-Ube2R1
Ube2R1
Ub
1. UBE2R1 C93K - unmodified
2. UBE2R1 C93K - modified
Modified peptide
Counterpart unmodified peptide
Peptide intensity
Gel slice
C.

**Fig. S2. Preparation and assembly of UBE2R1(C93K)-Ub for cryo-EM.** (**A**) Coomassie stained gel of preparative scale ubiquitin loading on UBE2R1(C93K). SEC (Size exclusion chromatography). The fractions containing Ub-Ube2R1(C93K) within the black box were pooled (**B**) Mass spectrometry analysis of ubiquitin loading onto the active site of UBE2R1(C93K). Schematic of the amino acid sequence of Ube2R1 (C93K) with ubiquitin modification (yellow, Ub) on the active site C93K, the GlyGly(K) modified peptide sequence is colored in red and the spectrum for this peptide is shown below. Andromeda score, 244.31 (**C**) The active site (C93K) residue is almost fully occupied with Ubiquitin. Coomassie stained gel of ubiquitin loading on UBE2R1 for mass spectrometry analysis. Peptide intensity for the GlyGly(K) modified and counterpart unmodified active site peptide (depicted in B) from the unmodified band (blue, gel slice 1) and the unmodified band (red, gel slice 2).

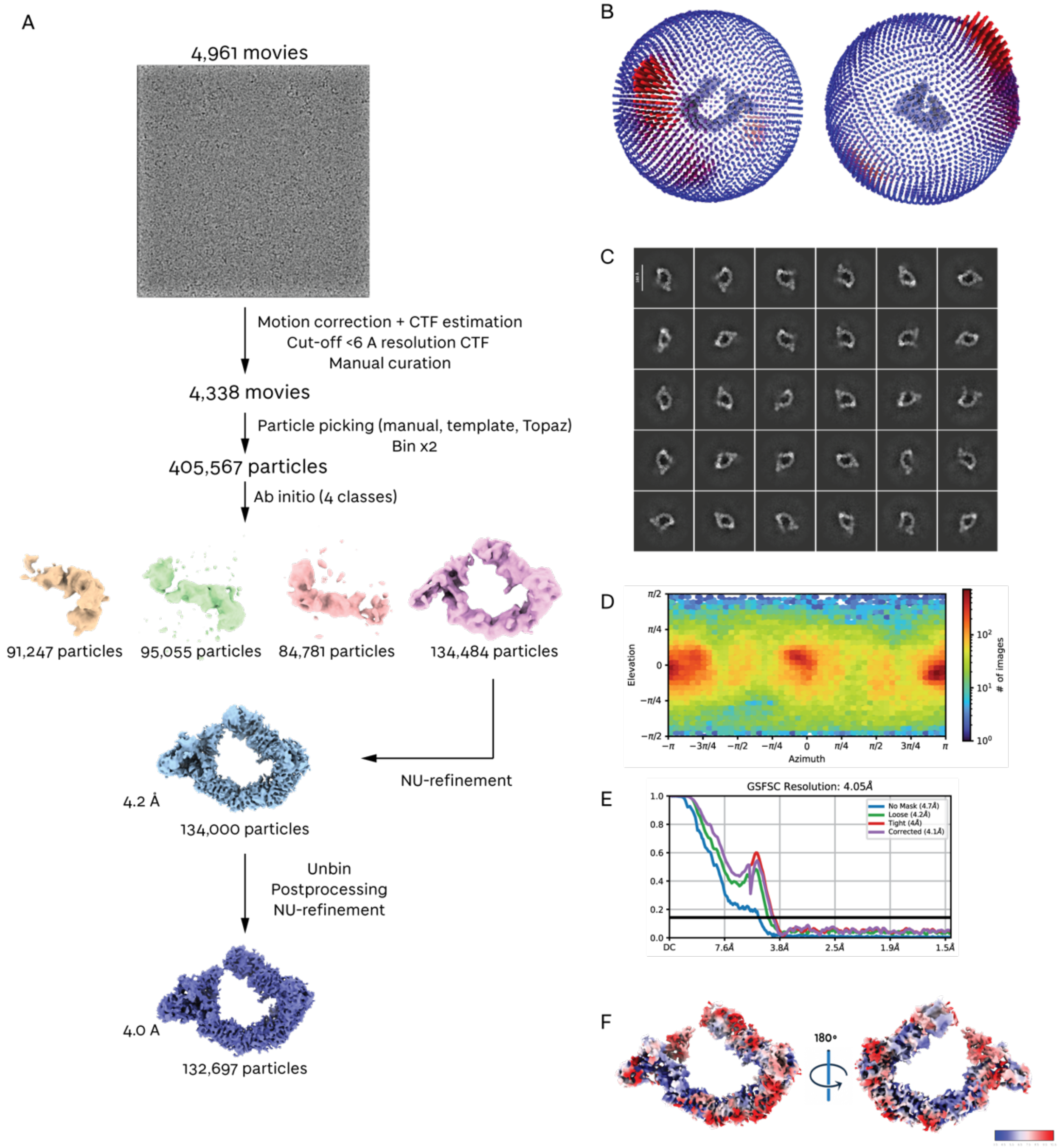

**Fig. S3. Cryo-EM image analysis for Brd4$^{BD2}$-MZ1-(NEDD8)-CRL2$^{VHL}$-UBE2R1-Ub.** (**A**) A schematic for the processing workflow for the 'open' non-crosslinked Brd4$^{BD2}$-MZ1-(NEDD8)-CRL2$^{VHL}$-UBE2R1-Ub complex used to generate cryo-EM maps. (**B**) 3D viewing direction distribution. (**C**) Selected 2D classes. (**D**) 2D viewing direction distribution. (**E**) Gold-standard Fourier shell correlation plot at 0.143. (**F**) Local resolution estimation.

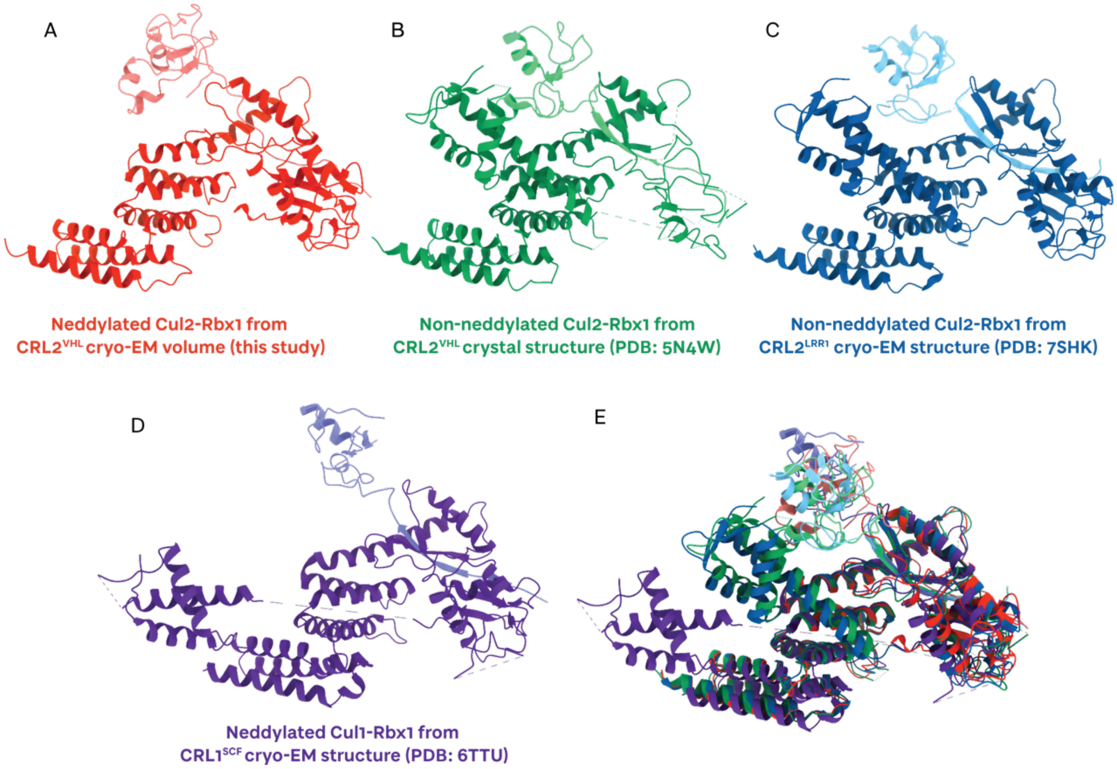

**Fig. S4. Structural alignments of CRL2 C-terminal domains (CTDs).** (**A**) CTD of Cullin 2 (red) and Rbx1 (pink) in the atomic model built from the ‘open’ cryo-EM structure in this study. The WHB domain of Cullin 2 is not modelled. The complex is neddylated, and Rbx1 appears to be in a mobile state. (**B**) CTD of Cullin 2 (dark green) and Rbx1 (light green) in the atomic model of the crystal structure of full-length Cul2-Rbx1 in complex with VHL-EloB-EloC (PDB: 5N4W) (*21*). The complex is unneddylated and the WHB domain of Cullin 2 and Rbx1 are closely packed. (**C**) CTD of Cullin 2 (dark blue) and Rbx1 (light blue) in the atomic model of the cryo-EM structure of full-length Cul2-Rbx1 in complex with LRR1 (PDB: 7SHK, EMDB-25127) (*22*). (**D**) CTD of Cullin 1 (dark purple) and Rbx1 (light purple) in the atomic model of the cryo-EM structure of full-length Cul1-Rbx1 in complex with SCF (PDB: 6TTU, EMDB-10585) (*23*). (**E**) Overlay all A, B, C and D atomic models. Rbx1 adopts a range of different orientations.

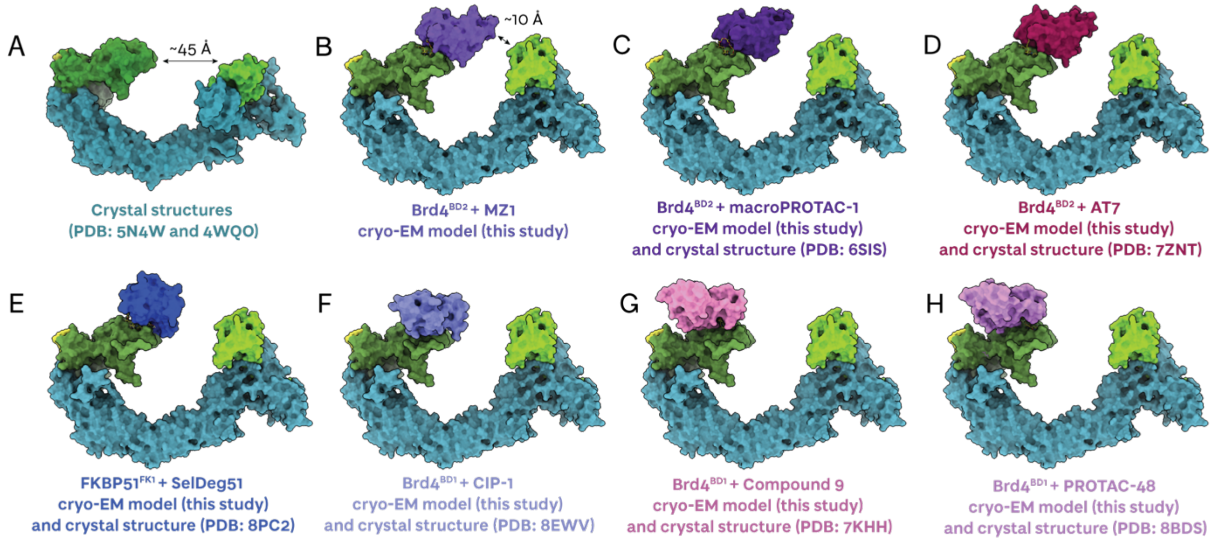

**Fig. S5. Comparison of degrader ternary complex crystal structures aligned with the cryo-EM volume from this study.** (**A**) Surafce representation of the aligned crystal structures of VHL-EloC-EloC-Cul2(NTD) (*24*) with VHL-EloC-EloC-Cul2-Rbx1 (5N4W) (*21*). (**B**) Surface representations of the atomic model of the Brd4$^{BD2}$-MZ1-(NEDD8)-CRL2$^{VHL}$ complex generated from this study. (**C**) Surface representations of the atomic model for the Brd4$^{BD2}$-macroPROTAC1-(NEDD8)-CRL2$^{VHL}$ complex (built from the cryo-EM volume from this study aligned with the crystal structure PDB: 6SIS) (*25*); (**D**) Surface representations of the atomic model of the Brd4$^{BD2}$-AT7-(NEDD8)-CRL2$^{VHL}$ complex (built from the cryo-EM volume from this study aligned with the crystal structure PDB: 7ZNT) (*26*); (**E**) Surface representations of the atomic model of the FKBP51$^{FK1}$-SelDeg51-(NEDD8)-CRL2$^{VHL}$ complex (built from the cryo-EM volume from this study aligned with the crystal structure PDB: 8PC2) (*27*); (**F**) Surface representations of the atomic model of the Brd4$^{BD1}$-CIP1-(NEDD8)-CRL2$^{VHL}$ complex (built from the cryo-EM volume from this study aligned with the crystal structure PDB: 8EWV) (*28*); (**G**) Surface representations of the atomic model of the Brd4$^{BD1}$-Compound9-(NEDD8)-CRL2$^{VHL}$ complex (built from the cryo-EM volume from this study aligned with the crystal structure PDB: 7KHH) (*29*); (**H**) Surface representations of the atomic model of the Brd4$^{BD1}$-PROTAC-48-(NEDD8)-CRL2$^{VHL}$ complex (built from the cryo-EM volume from this study aligned with the crystal structure PDB: 8BDS) (*30*).

A

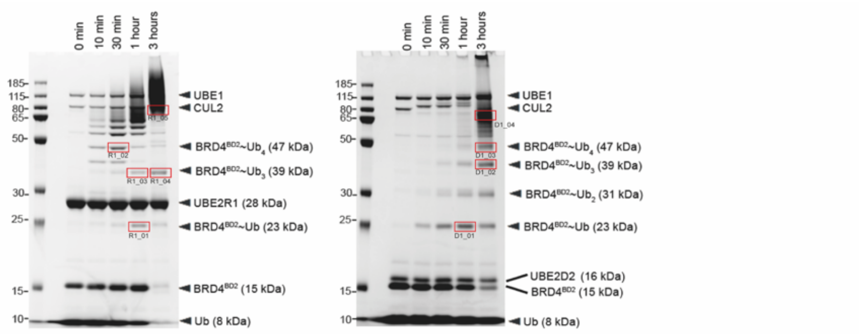

B

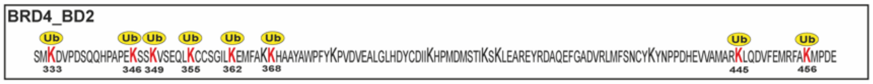

C

| | Modified by UBE2R1 | | | | | Modified by UBE2D2 | | | |
|---|---|---|---|---|---|---|---|---|---|
| Gel slice | R1_01 | R1_02 | R1_03 | R1_04 | R1_05 | D1_01 | D1_02 | D1_03 | D1_04 |
| BRD4_BD2 peptide sequence coverage (%) | 83.1 | 95.4 | 50.8 | 50.0 | 95.4 | 95.4 | 74.6 | 74.6 | 95.4 |

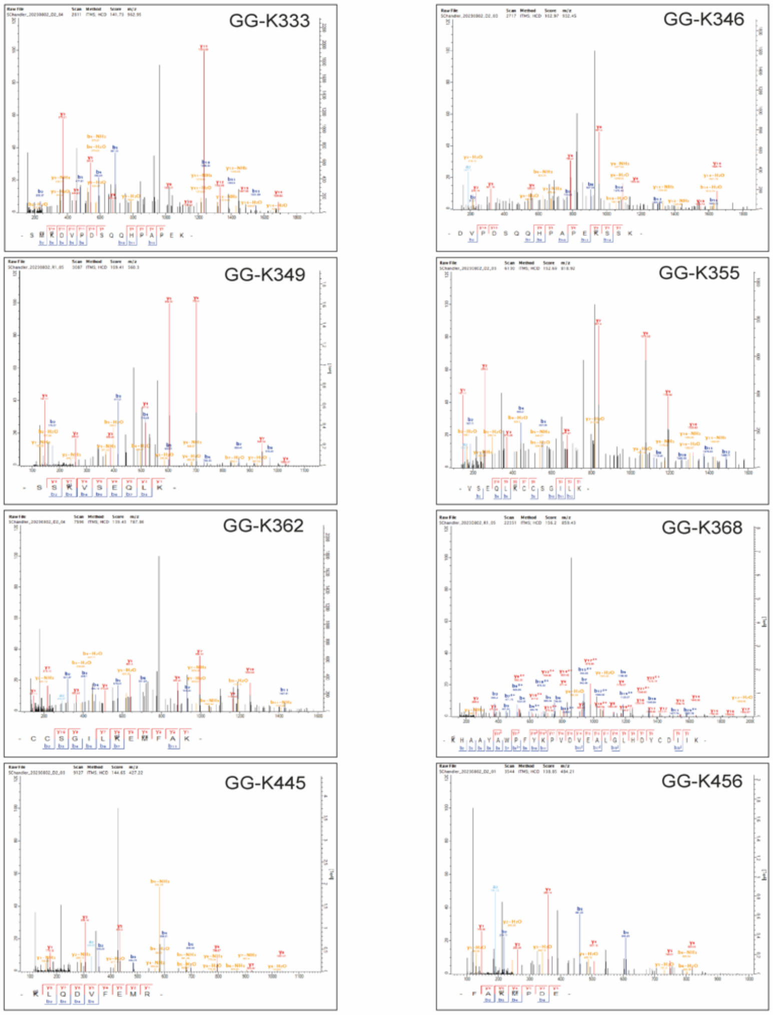

D

**Fig. S6. Identification of Brd4$^{BD2}$ ubiquitination sites by mass spectrometry.** (**A**) Coomassie-stained SDS-PAGE of *in vitro* Brd4$^{BD2}$ ubiquitination assays perfomed in the presence of

UBE2R1 or UBE2D2. The red boxes show the regions of the gel that were excised for MS analysis. Below the red box is the name of each slice matching to the raw MS data file and spectra below. (**B**) Linear sequence-based representation of the ubiquitination sites identified on Brd4$^{BD2}$ ('Ub' in yellow circle above lysine residue represents ubiquitin-modification). (**C**) Peptide sequence coverage (%) of Brd4$^{BD2}$ by gel slice. (**D**) Best identified spectra for the Brd4$^{BD2}$ GlyGly(K) modified peptides across all gel slices. Beneath each spectrum is the y (red) and b (blue) series ions that were detected for each peptide. In the top right corner is the GG-K residue, numbering based on the Brd4$^{BD2}$ construct used in the assay, depicted in B. 'Score' is Andromeda score; GG-K333 (141.73), GG-K346 (102.97), GG-K349(169.41), GG-K355(152.69), GG-K362(139.43), GG-K368(156.2), GG-K445(144.65), GG-K456(138.85).

| | | Modified by UBE2R1 | | | | | | | | | | Modified by UBE2D2 | | | | | | | |
|---|---|---|---|---|---|---|---|---|---|---|---|---|---|---|---|---|---|---|---|
| Time (hrs) | | 0.5 | 1 | 1 | 3 | 3 | 0.5 | 1 | 1 | 3 | 3 | 1 | 3 | 3 | 3 | 1 | 3 | 3 | 3 |
| Molecular weight of excised band (kDa) | | 47 | 23 | 39 | 39 | 65 | 47 | 23 | 39 | 39 | 79 | 23 | 39 | 47 | 63 | 23 | 39 | 47 | 79 |
| Proteins | GG-K Position | Andromeda Score | | | | | Log10 peptide intensity | | | | | Andromeda Score | | | | Log10 peptide intensity | | | |
| BRD4_BD2 | 333 | | | | | 103 | | | | | 6.26 | 87 | 106 | 115 | 142 | 6.05 | 6.86 | 7.25 | 7.47 |
| BRD4_BD2 | 346 | | | | | 62 | | | | | 6.02 | 82 | 88 | 103 | 98 | 6.12 | 6.69 | 7.16 | 7.31 |
| BRD4_BD2 | 349 | | | | | 169 | | | | | 6.74 | 128 | 109 | 150 | 126 | 7.39 | 7.16 | 7.45 | 7.74 |
| BRD4_BD2 | 355 | | | | | 113 | | | | | 6.77 | 127 | 101 | 153 | 131 | 6.78 | 7.28 | 7.66 | 8.14 |
| BRD4_BD2 | 362 | | | | | | | | | | | 64 | 91 | 78 | 139 | 6.23 | 6.51 | 6.83 | 7.13 |
| BRD4_BD2 | 368 | | | | | 156 | | | | | 8.88 | | | | 137 | | | | 7.96 |
| BRD4_BD2 | 445 | | | | | 86 | | | | | 7.18 | 79 | 114 | 145 | 98 | 6.33 | 7.00 | 7.17 | 7.28 |
| BRD4_BD2 | 456 | 91 | | | | 107 | 7.44 | | | | 8.10 | 139 | 128 | 128 | 121 | 8.27 | 8.58 | 8.89 | 9.04 |
| UBIQUITIN | 6 | 122 | 111 | | 71 | 159 | 6.76 | 6.63 | | 6.48 | 7.53 | | 176 | 158 | 201 | | 7.39 | 7.75 | 8.55 |
| UBIQUITIN | 11 | 141 | 182 | 127 | 136 | 331 | 7.47 | 7.55 | 6.37 | 7.63 | 9.13 | 135 | 267 | 273 | 362 | 7.19 | 9.08 | 9.53 | 10.08 |
| UBIQUITIN | 27 | 163 | 175 | | | 188 | 6.67 | 6.47 | | | 7.44 | | 71 | 135 | 93 | | 5.79 | 6.10 | 6.71 |
| UBIQUITIN | 33 | | | | | | | | | | | | | 63 | 99 | | | | 6.25 |
| UBIQUITIN | 48 | 199 | 172 | 116 | 163 | 320 | 10.22 | 10.26 | 7.97 | 9.16 | 10.74 | 112 | 159 | 178 | 179 | 7.11 | 7.40 | 9.02 | 9.99 |
| UBIQUITIN | 63 | 108 | 139 | | 126 | 285 | 7.30 | 7.44 | | 7.23 | 8.79 | 59 | 223 | 236 | 239 | 6.07 | 8.64 | 9.56 | 10.19 |

**Fig. S7. Ubiquitination sites on Brd4$^{BD2}$ were mapped by mass spectrometry following an *in vitro* ubiquitination assay containing neddylated-CRL2$^{VHL}$, Brd4$^{BD2}$, MZ1, Uba1 and either UBE2R1 or UBE2D2.** Time points were taken at 0 minutes, 10 minutes, 0.5 hours, 1 hour and 3 hours. Proteins were separated by SDS-page analysis. Gel slices containing ubiquitin modified Brd4$^{BD2}$ across timepoints and increasing molecular weight were excised from the gel and were analyzed by mass spectrometry. The Log10 peptide intensities and andromeda scores of the identified Brd4$^{BD2}$ and ubiquitin peptides are listed.

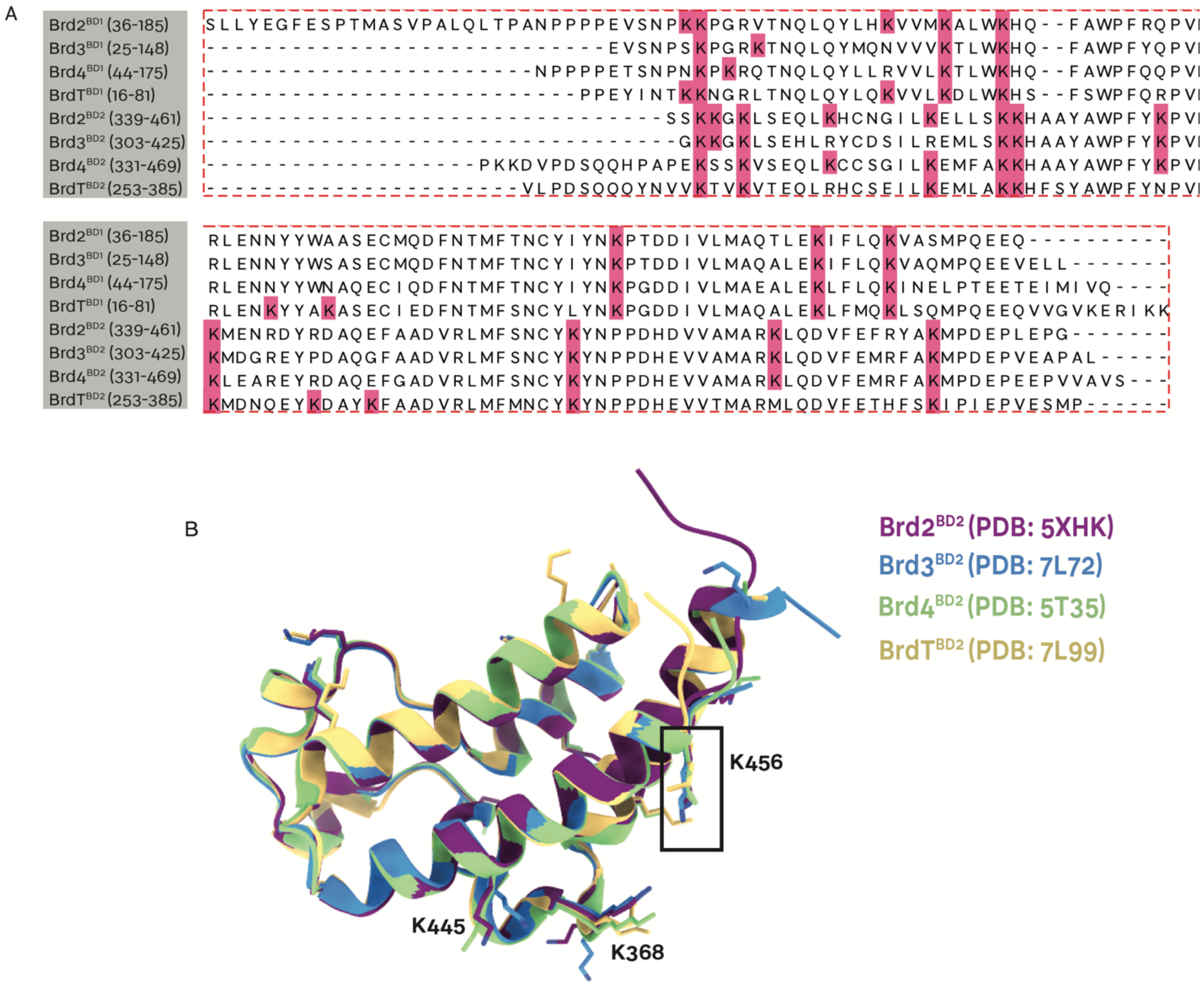

**Fig. S8. Sequence and structural alignments of BET bromodomains, highlighting lysine residues available for ubiquitination.** (**A**) Multiple sequence alignment of bromodomain-1 and bromodomain-2 for Brd2, Brd3, Brd4, and BrdT. The position of all lysines residues are highlighted in pink. (**B**) Structural alignment of bromodomain-2 of Brd2 (purple), Brd3 (blue), Brd4 (green), and BrdT (yellow) from the crystal structures PDB: 5XHK (*31*), 7L72 (*32*), 5T35 (*6*) and 7L99 (*33*). The lysine residues are shown as sticks with coloring according to the bottom legend. K456 of $Brd4^{BD2}$ is annotated, along with K445 and K368.

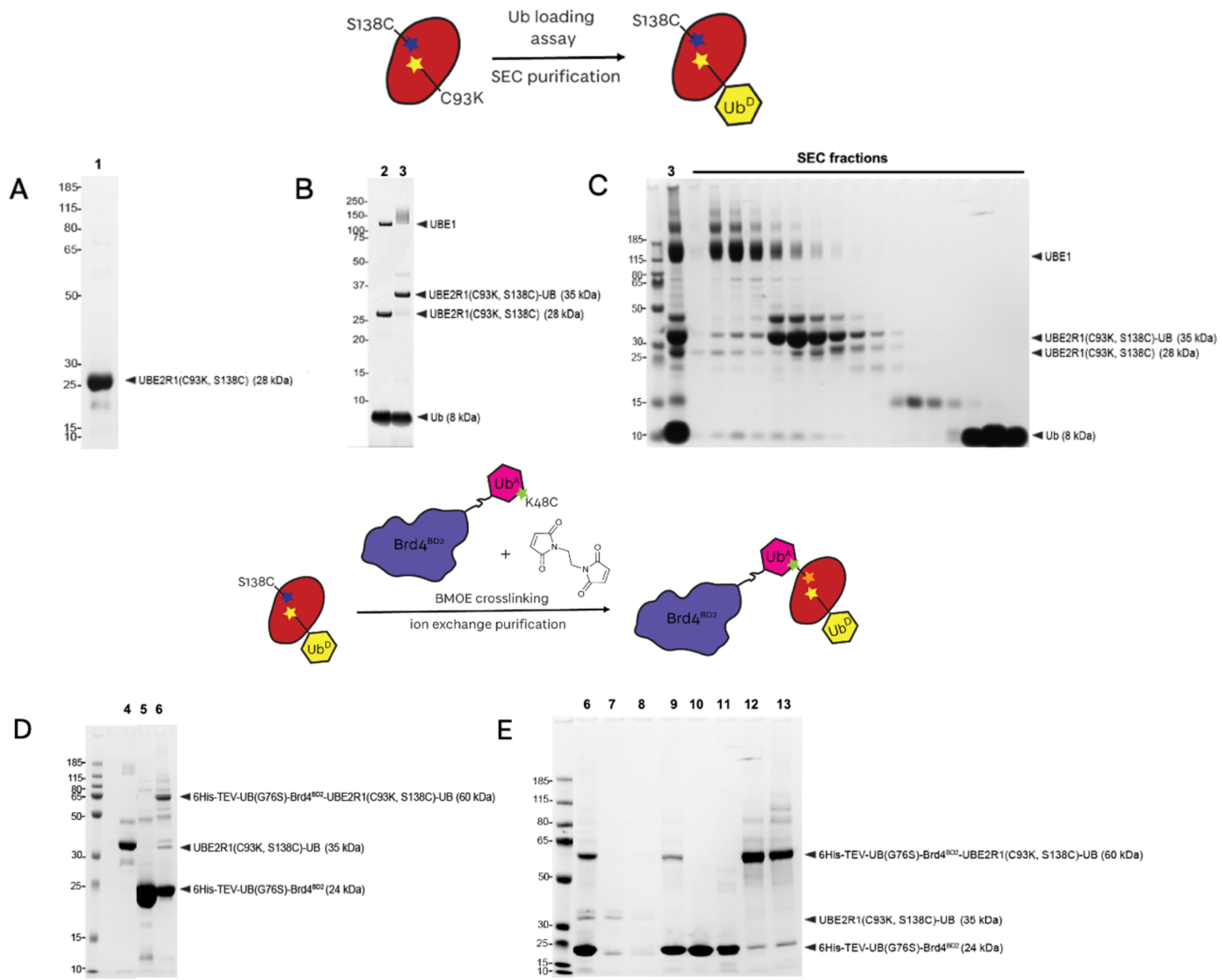

**Fig. S9. Preparation of the His$_6$-TEV-Ub(G76S)-Brd4$^{BD2}$-BMOE- Ube2R1(C93K,S138C, C191S,C223S)-Ub species for the 'closed' crosslinked cryo-EM structure, resolved by SDS-PAGE and Coomassie staining.** (**A**) Lane 1: Ube2R1(C93K,S138C,C191S,C223S) [hereafter referred to as Ube2R1(C93K,S138C)] is expressed and purified. (**B**) Lane 2: Ube1 and ubiquitin are added to Ube2R1(C93K,S138C), the reaction is at the 0 minute timepoint ; Lane 3: ATP has been added, the reaction is at the 18 hour timepoint and Ube2R1(C93K,S138C) is loaded with ubiquitin forming Ube2R1(C93K,S138C, C191S, C223S)-Ub. (**C**) Lane 3: the reaction mixture where Ube2R1(C93K,S138C)-Ub is formed is purified by SEC shown in Fractions 16 to 32; (**D**) Lane 4: purified Ube2R1(C93K,S138C)-Ub is reacted with BMOE; Lane 5: His$_6$-TEV-Ub(G76S,K48C)-Brd4BD2 is expressed, purified and desalted; Lane 6: BMOE-Ube2R1(C93K,S138C)-Ub is reacted with His$_6$-TEV-Ub(G76S,K48C)-Brd4$^{BD2}$, forming His$_6$-TEV-Ub(G76S,K48C)-Brd4$^{BD2}$-BMOE- Ube2R1(C93K,S138C)-Ub ; (**E**) Lane 6: the crude reaction mixture where His$_6$-TEV-Ub(G76S)-Brd4$^{BD2}$-BMOE- Ube2R1(C93K,S138C)-Ub is formed ; Lane 7: Ni NTA flow through (20 mM imidazole) ; Lane 8: Ni NTA wash (20 mM imidazole) ; Lane 9: Ni NTA elution (500 mM imidazole) ; Lanes 10-11: ion exchange chromatography elution fractions containing excess His$_6$-TEV-Ub(G76S, K48C)-Brd4$^{BD2}$ ; Lanes 12-13: ion exchange chromatography elution fractions containing the purified product Brd4$^{BD2}$-Ub(G76S,K48C)-BMOE-Ube2R1(C93K,S138C)-Ub.

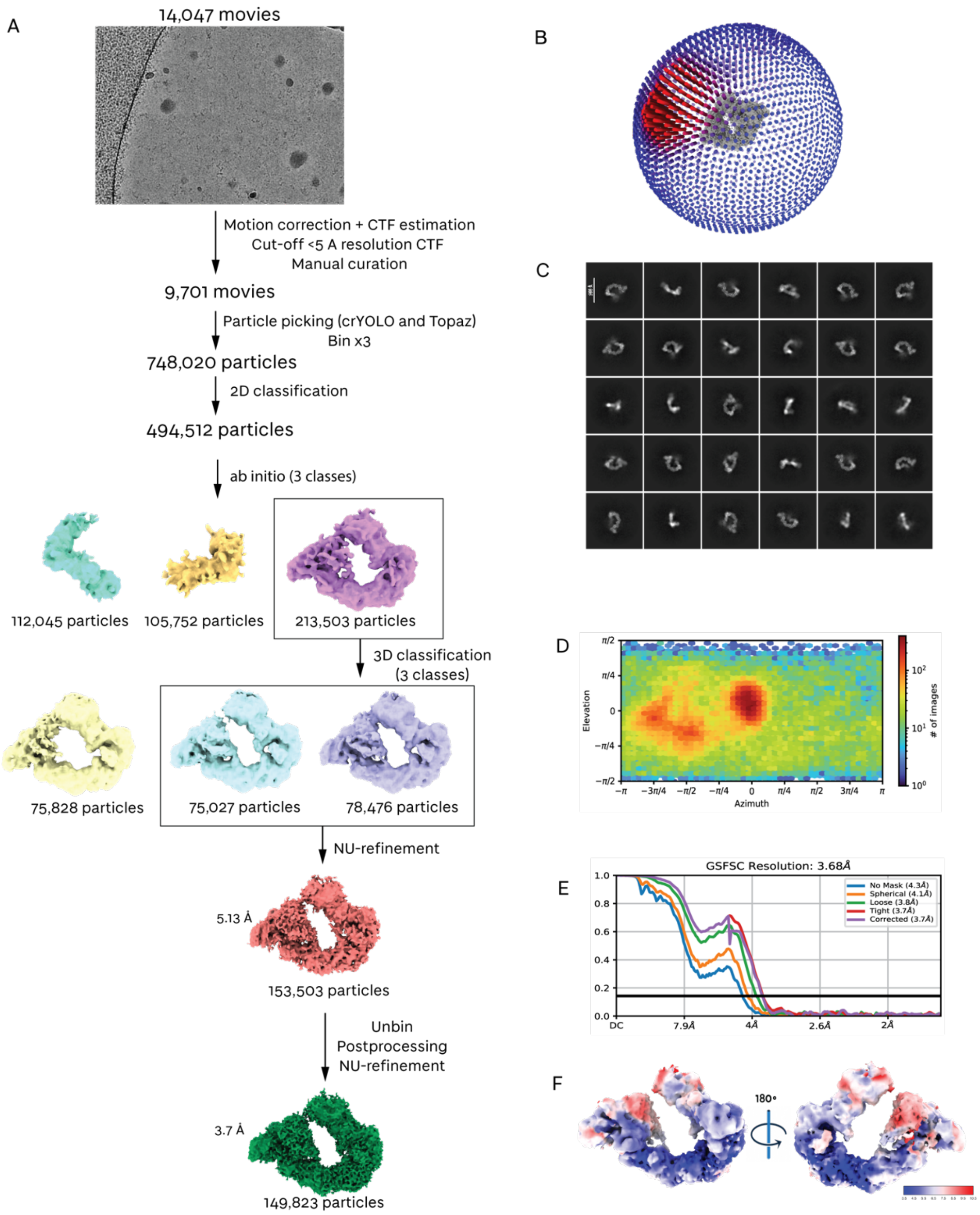

**Fig. S10. Cryo-EM image analysis for 'closed' crosslinked (NEDD8)-CRL2$^{VHL}$-MZ1-Brd4$^{BD2}$-Ub(G76S, K48C)-UBE2R1(C93K, S138C, C191S, C223S)-Ub complex.** (**A**) A schematic for the processing workflow for the 'closed' crosslinked (NEDD8)-CRL2$^{VHL}$-MZ1-Brd4$^{BD2}$-Ub(G76S, K48C)-UBE2R1(C93K, S138C)-Ub complex used to generate cryo-EM maps. (**B**) 3D viewing direction distribution. (**C**) Selected 2D classes. (**D**) 2D viewing direction distribution. (**E**) Gold-standard Fourier shell correlation plot at 0.143. (**F**) Local resolution estimation.

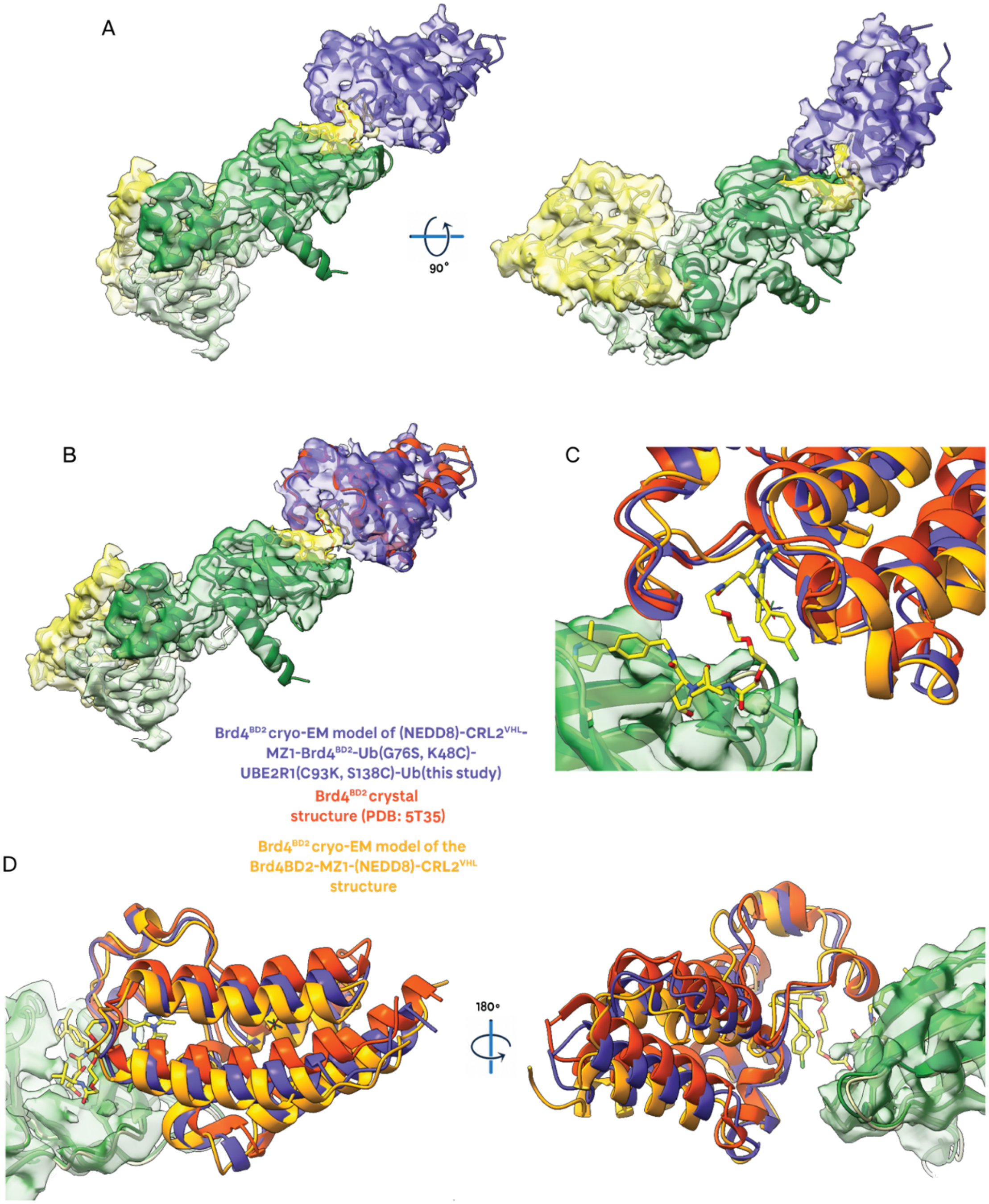

**Fig. S11. Cryo-EM volume and fitted atomic models from the ‘closed’ crosslinked structure of (NEDD8)-CRL2$^{VHL}$-MZ1-Brd4$^{BD2}$-Ub(G76S, K48C)-UBE2R1(C93K, S138C)-Ub.** (**A**) Cryo-EM map (transparent volume) and atomic model for the (NEDD8)-CRL2$^{VHL}$-MZ1-Brd4$^{BD2}$-Ub(G76S, K48C)-UBE2R1(C93K, S138C)-Ub structure. (**B**) Cryo-EM map (transparent volume) and atomic model for the (NEDD8)-CRL2$^{VHL}$-MZ1-Brd4$^{BD2}$-Ub(G76S, K48C)-UBE2R1(C93K, S138C)-Ub structure (purple). The atomic model for the crystal structure 5T35 overlays exactly with the atomic model from cryo-EM (alignment against VHL), with deviation observed for Brd4$^{BD2}$ (red). (**C**) The VHL-MZ1-Brd4$^{BD2}$ interface with Brd4$^{BD2}$ from (NEDD8)-CRL2$^{VHL}$-MZ1-Brd4$^{BD2}$-Ub(G76S, K48C)-UBE2R1(C93K, S138C)-Ub shown in purple (this study)., Brd4$^{BD2}$ from the crystal structure (5T35) shown in red (*6*), and the Brd4$^{BD2}$ from Brd4$^{BD2}$-MZ1-(NEDD8)-CRL2$^{VHL}$-UBE2R1-Ub shown in orange (this study). (**D**) Visualisation of the re-orientation of Brd4$^{BD2}$ relative to the interface VHL-MZ1-Brd4$^{BD2}$ interface, with Brd4$^{BD2}$ from (NEDD8)-CRL2$^{VHL}$-MZ1-Brd4$^{BD2}$-Ub(G76S, K48C)-UBE2R1(C93K, S138C)-Ub shown in purple (this study)., Brd4$^{BD2}$ from the crystal structure (5T35) shown in red (*6*), and the Brd4$^{BD2}$ from Brd4$^{BD2}$-MZ1-(NEDD8)-CRL2$^{VHL}$-UBE2R1-Ub shown in orange (this study).

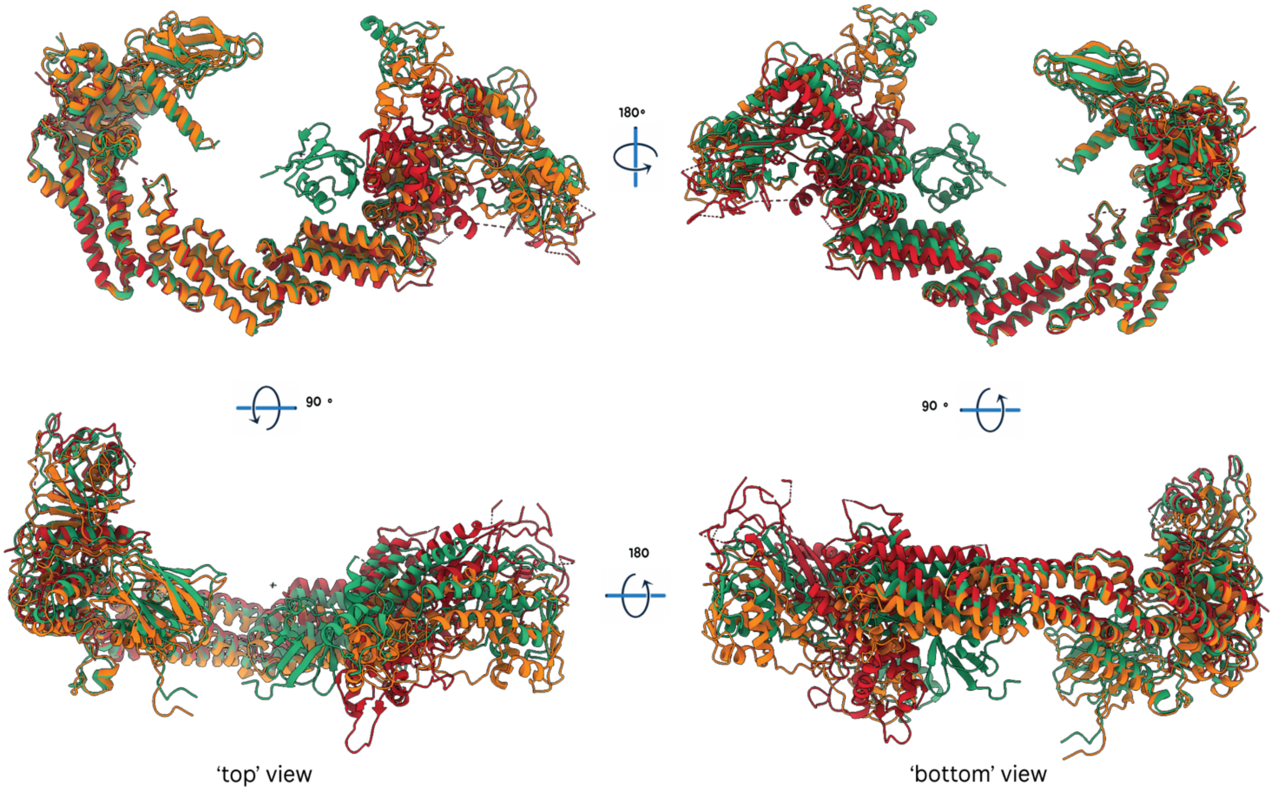

**Fig. S12. Superimposition of CRL2$^{VHL}$ atomic models.** Atomic models include: the atomic model built into the cryo-EM map of the (NEDD8)-CRL2$^{VHL}$-MZ1-Brd4$^{BD2}$-Ub(G76S, K48C)-UBE2R1(C93K, S138C)-Ub structure (this study, green); the atomic model built into the cryo-EM map of the Brd4$^{BD2}$-MZ1-(NEDD8)-CRL2$^{VHL}$-UBE2R1-Ub structure (this study, orange); the atomic model of the crystal structure (PDB: 5N4W) of the unneddylated CRL2$^{VHL}$ ((*21*) red). The atomic models were aligned again the first 170 residues of Cullin 2. Hinging of Cullin 2 was observed at the C-terminus of the helical bundle 2, leading to a range of conformations being sampled for the Cullin C-terminal domain. Rbx1 adopted different conformations dependent on whether the complex was neddylated, crosslinked with UBE2R1 or non-crosslinked.

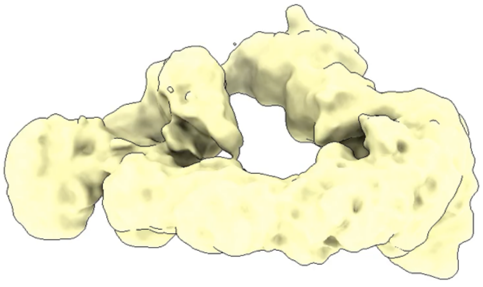

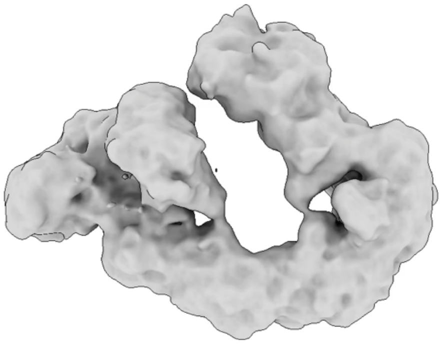

**Movies S1 and S2. Results of 3D Variability analysis of the 'closed' crosslinked structure (NEDD8)-CRL2$^{VHL}$-MZ1-Brd4$^{BD2}$-Ub(G76S, K48C)-UBE2R1(C93K, S138C)-Ub.** The analysis was performed on 149,823 particles in CryoSPARC v4.4.1 (*9, 34*). The output was clustered into groups of 20 frames and combined as linear 'movies' of volumes. The trajectories displayed highlight the extent to which the complex is dynamic and outline the range of motion available.

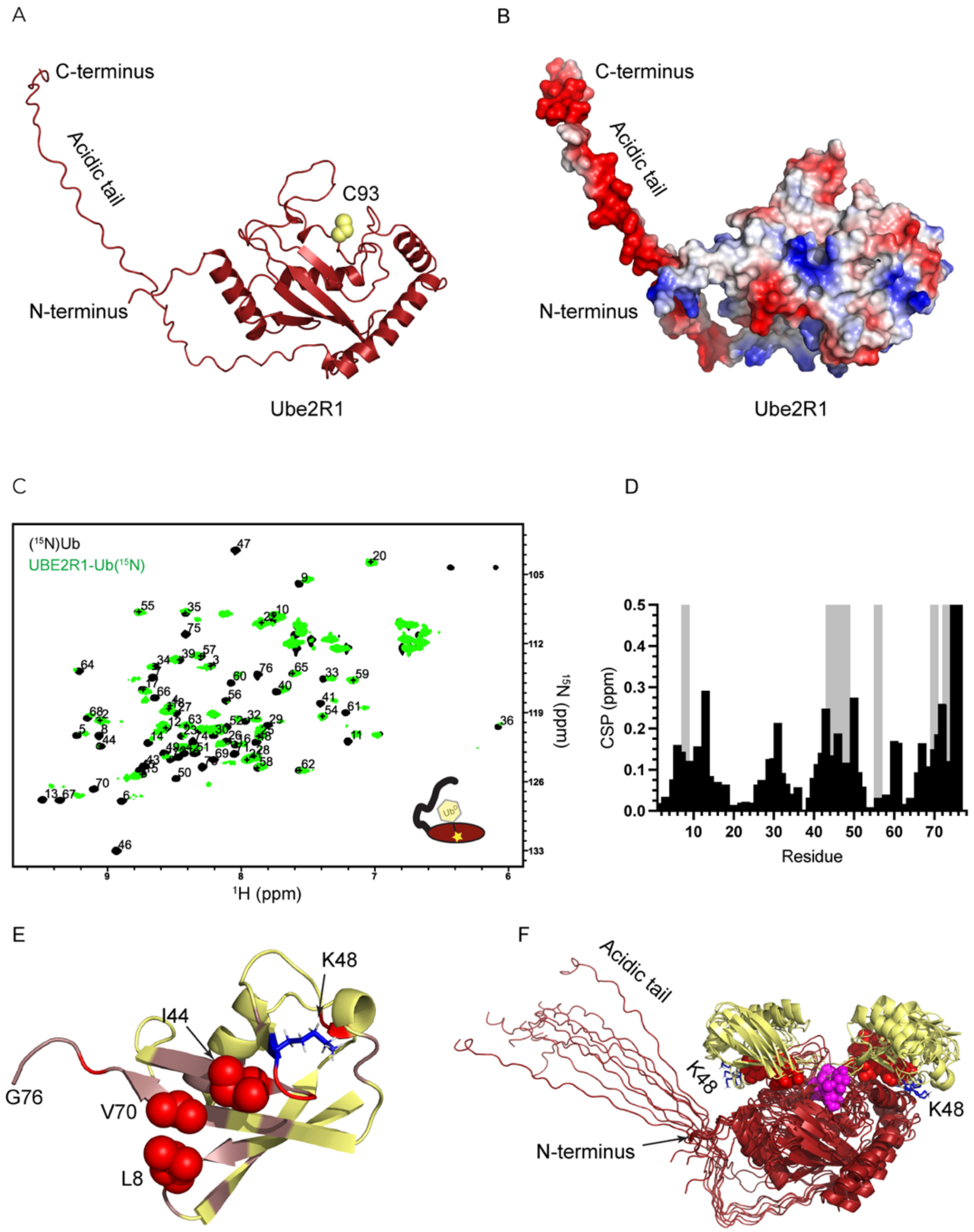

**Fig. S13. NMR analysis of Ub$^{D}$ on Ube2R1.** The stable Ube2R1$^{C93K}$-Ub($^{15}$N) conjugate was made on full length human Ube2R1 with (**A**) AlphaFold2 model and (**B**) surface representation with electrostatic potential highlighting the acidic tail and regulator loops on the active site C93. (**C**) Overlay of $^{1}$H-$^{15}$N-HSQC spectra of unconjugated Ub($^{15}$N) (black) and Ube2R1$^{C93K}$-Ub($^{15}$N) (green). (**D**) Signal attenuations (red) and above average CSPs (salmon) are mapped on Ub highlighting the interaction with the L8, I44, V70 hydrophobic path and K48 (red sitck). (**E**) Residue specific CSPs (black bars) and signal attenuations (grey) for residues. (**F**) Poses from top clusters of modeling Ub$^{D}$ on the active site of UBE2R1 with HADDOCK (*35*), restrained with C93 of UBE2R1 and G76 of Ub$^{D}$ (magenta spheres). Residues from Ub$^{D}$ model the interaction through hydrophobic patch (L8,I44,V70) green sticks and K48 as blue sticks.

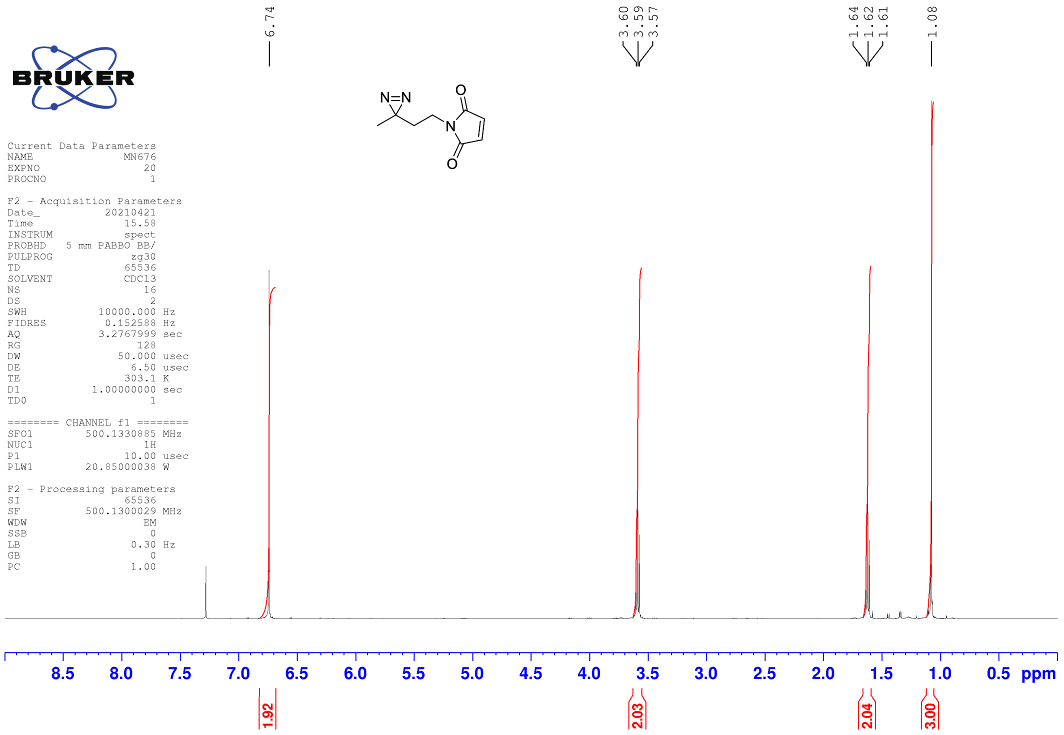

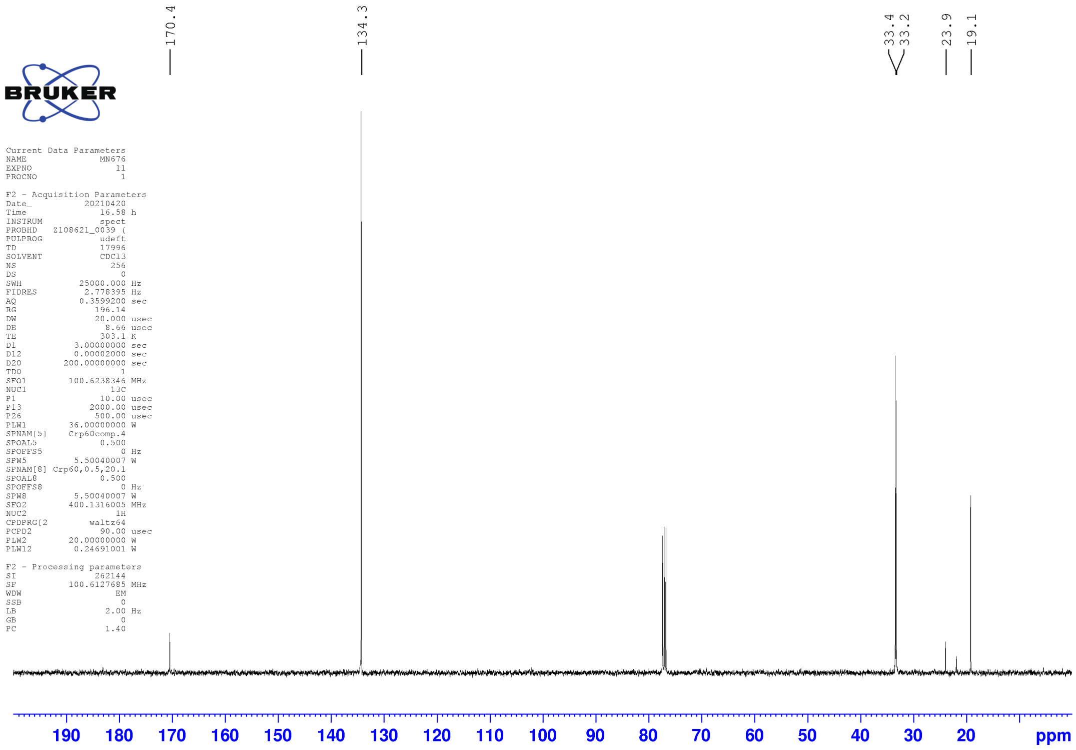

**Fig. S14. NMR spectra of 1-(2-(3-methyl-3H-diazirin-3-yl)ethyl)-1H-pyrrole-2,5-dione.**

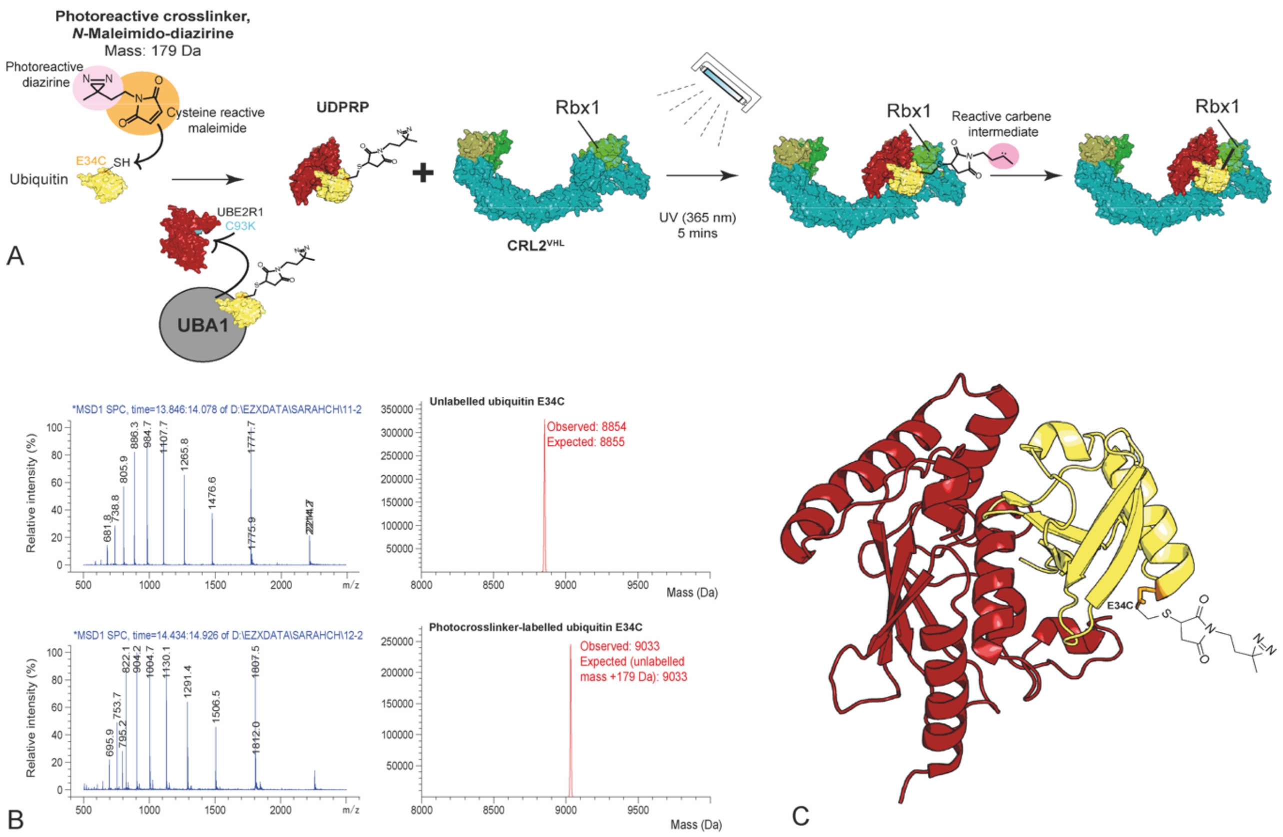

**Fig. S15. Design of ubiquitin-directed photoreactive probe (UDPRP) to capture CRL2$^{VHL}$ in-solution.** The production of the probe starts with the site-specific labelling of ubiquitin E34C with the photoreactive crosslinker N-maleimido-diazirine. The UDPRP is generated by conjugating the photocrosslinker-labelled ubiquitin to UBE2R1 C93K using recombinant human E1 (UBA1) forming a stable isopeptide linked E2-Ub conjugate. The UDPRP is then purified by size exclusion chromatography. The modified UBE2R1(C93K)-Ub(E34C)-crosslinker UDPRP species now displays the photoreactive crosslinker N-maleimido diazirine which when exposed to UV light (365 nm) forms a reactive carbene intermediate which can rapidly insert into X-H bonds (including O-H, N-H, S-H and C-H bonds) and can therefore trap proteins in close proximity (*8*). This forms a crosslinked product (MW 47 kDa) that is resistant to separation by SDS-page analysis and can therefore be visualized by Coomassie or by immunoblotting for components of the photo-crosslinked product, or the photo-crosslinked peptide can be detected by mass-spectrometry. (**A**) Schematic representation of the design and application of the ubiquitin-directed photoreactive probe (UDPRP). Upon reaction with (NEDD8)-CRL2$^{VHL}$ and exposure to UV light (365 nm), the carbene can react with X-H bonds in Rbx1. PDB codes; ubiquitin (1UBQ), UBA1 in complex with ubiquitin (6DC6), structure of UBE2R2 (3RZ3) was used for UBE2R1, CRL2$^{VHL}$ (5N4W). (**B**) Site specific labelling of ubiquitin E34C with the photocrosslinker N-Maleimido-diazirine. Raw ESI-MS spectrum of samples containing unlabeled and labelled ubiquitin E34C. Observed mass shift was equal to the mass of N-maleimido-diazirine (179 Da). (**C**) Structural representation of the UDPRP. Crystal structure of a stable isopeptide linked UBE2R2-Ub conjugate (PDB: 6NYO). The presented crystal structure contains wild-type ubiquitin however in the UDPRP an E48C mutant ubiquitin was used to enable the site-specific labelling of N-maleimido-diazirine.

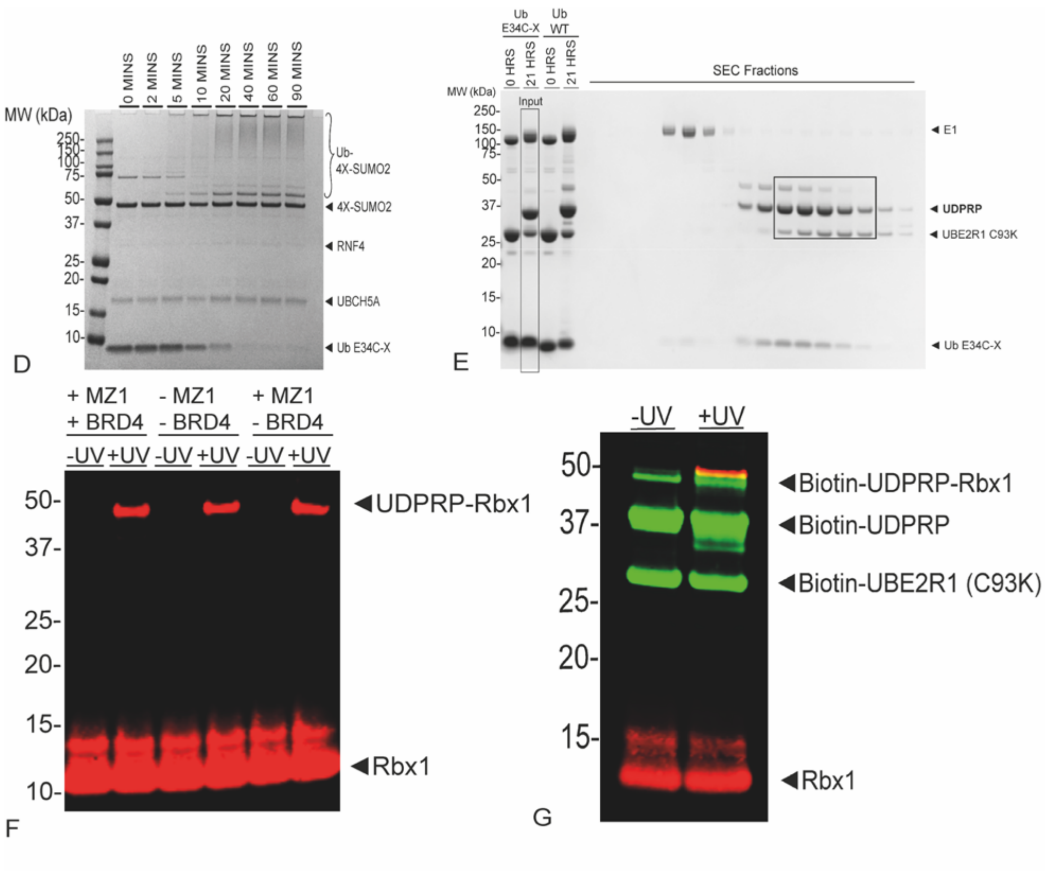

(**D**) Coomassie stained gel to demonstrate that the photocrosslinker labelled ubiquitin E34C (Ub E34C-X) is active in an *in vitro* ubiquitination assay with the SUMO-targeted E3 ligase, the E2 UbcH5a/UBE2D2 and UBA1. (**E**) Coomassie stained gel of the preparative loading of photocrosslinker labelled ubiquitin E34C to UBE2R1 (C93K) and purification of the ubiquitin-directed photoreactive probe (UDPRP). The loading efficiency of photocrosslinker labelled ubiquitin (Ub E34C-X) to the active site of UBE2R1 is comparable to WT ubiquitin. A 0 hour time point taken prior to the addition of ATP, after 21 hours the reaction containing the Ub E34C-X (input) was separated by size exclusion chromatography (SEC). The black box indicates the the fractions which were pooled. (**F**) Immunoblot against Rbx1 shows the recruitment of the UDPRP to Rbx1 occurs independently of BRD4$^{BD2}$ and MZ1. UDPRP photo-crosslinking assay containing (NEDD8)-CRL2$^{VHL}$ (3 µM), UDPRP (10 µM), +/- BRD4$^{BD2}$ (17 µM), +/- MZ1 (4.5 µM). **(G)** Immunoblot for Rbx1 (red) and biotinylated-UBE2R1 (green) indicates that the photo-crosslinked product (orange) contains biotinylated UDPRP (UBE2R1-Ub) and Rbx1. UDPRP photo-crosslinking assay containing (NEDD8)-CRL2$^{VHL}$ (3 µM) and biotinylated-UDPRP (10 µM). The biotinylated-UDPRP contained some unmodified UBE2R1 (C93K) a remnant of the ubiquitin loading assay.

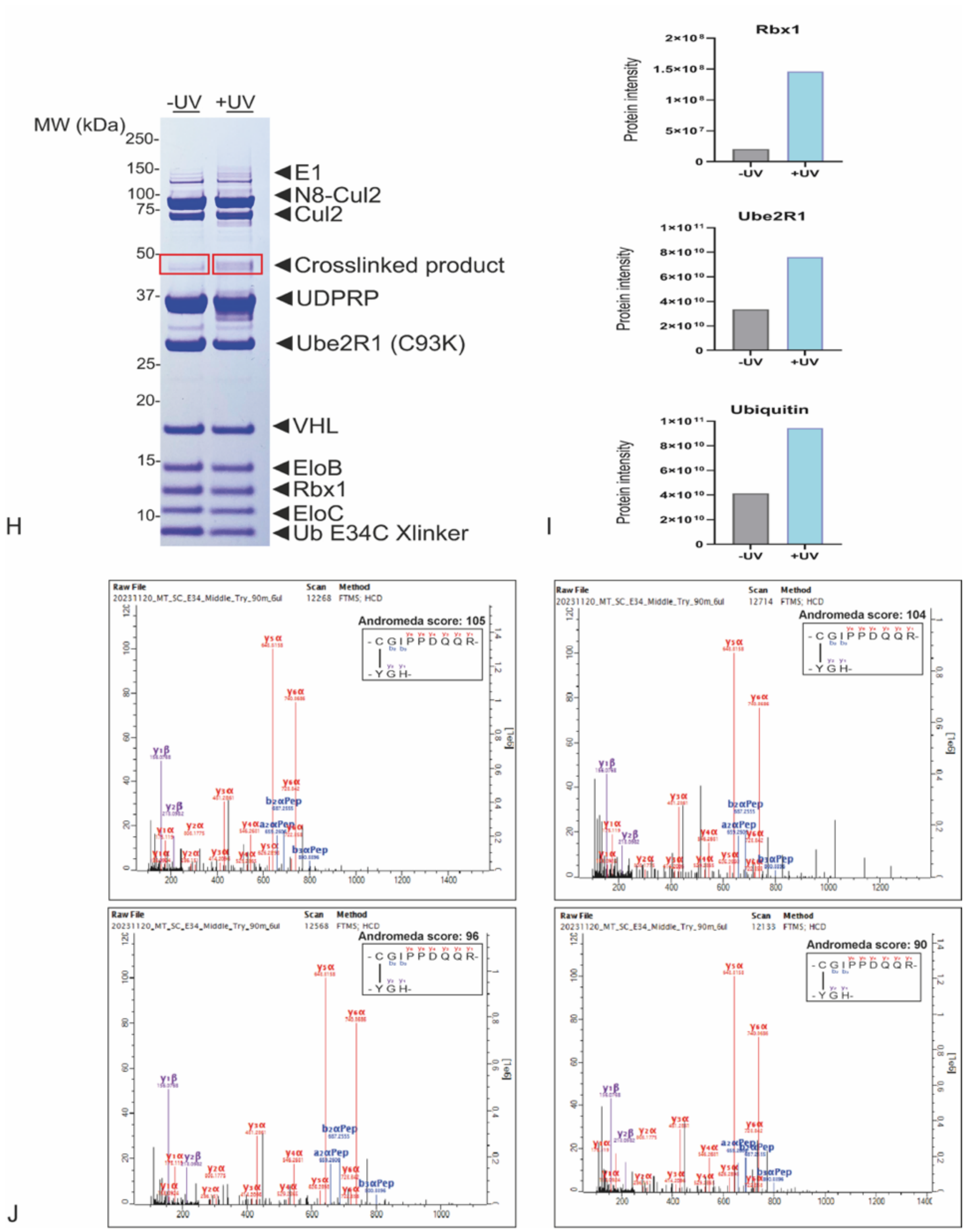

(**H**) Coomassie stained gel of the photocrosslinking UDPRP assay for MS analysis. The red boxes show the regions of the gel that were excised. (**I**) Protein intensity determined by mass spectrometry verifies the presence of Rbx1 in the photo-crosslinked product. Intensities for Rbx1, UBE2R1, and Ub from the two bands shown in red in (**H**)**,** left hand side (-UV) and right-hand side (+UV). (**J**) Ubiquitin-Rbx1 inter-protein crosslinked peptides detected linking ubiquitin E34C to the C-terminus of Rbx1. Spectra for crosslinked peptides identified in the +UV gel slice (shown in (**H**)). The crosslinked peptide is shown in the top right-hand corner with the y (purple and red) and b (blue) series ions detected.

A

| | UBE2R1 atom | Brd4BD2 atom | Distance (Å) |
|---|---|---|---|
| Cα-atom to Cα-atom | **C93 Cα-atom** | **K456 Cα-atom** | **24.08** |
| | C93 Cα-atom | K445 Cα-atom | 19.58 |
| | C93 Cα-atom | K368 Cα-atom | 17.62 |
| | C93 Cα-atom | K362 Cα-atom | 26.58 |
| | C93 Cα-atom | K355 Cα-atom | 32.64 |
| Thioester carbonyl C-atom to N-atom | **C93 thioester carbonyl C-atom** | **K456 N-atom** | **18.41** |
| | C93 thioester carbonyl C-atom | K445 N-atom | 19.22 |
| | C93 thioester carbonyl C-atom | K368 N-atom | 15.15 |
| | C93 thioester carbonyl C-atom | K362 N-atom | 27.92 |
| | C93 thioester carbonyl C-atom | K355 N-atom | 35.81 |

B

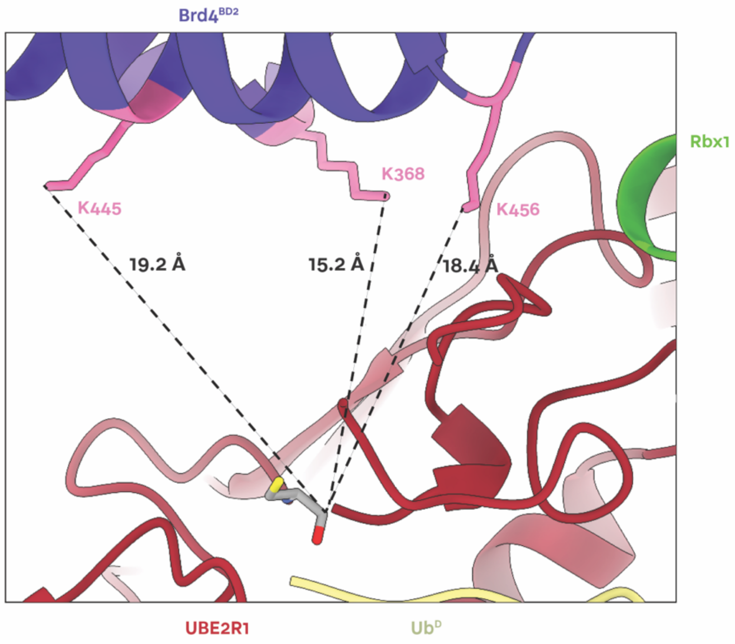

**Fig. S16. Measurements between ubiquitinated lysine residues on Brd4$^{BD2}$ (as identified by mass spectrometry) and the catalytic cysteine site of UBE2R1.** (**A**) Summary of the lysine residues on Brd4$^{BD2}$ identified by mass spectrometry as being ubiquitinated by UBE2R1. The measurements were performed between the Cα carbons of the ubiquitintated Lys and the Cα carbon of the Cys93 residue of UBE2R1. Measurements were also were performed between the nucleophilic nitrogen atom of the ubiquitintated Lys side-chain, and the Cys93 thioester carbonyl C-atom of UBE2R1, corresponding to electrophilic carboxyl targeted during ubiquitination. Measurements were performed using UCSF ChimeraX (*13*). (**B**) Schematic representation of the distances measured between the three closest lysine residues of Brd4$^{BD2}$ to the active site cysteine site of UBE2R1. All three lysines appear to be positioned at a similar distance, but adopt different geometries for the nucleophilic attack of the active site cysteine.

| | *'Open' non-crosslinked structure: Brd4$^{BD2}$-MZ1-(NEDD8)-CRL2$^{VHL}$-UBE2R1(C93K)-Ub (EMDB-19569) (PDB 8RWZ)* | *'Closed' crosslinked structure: (NEDD8)-CRL2$^{VHL}$-MZ1-Brd4$^{BD2}$-Ub(G76S, K48C)-UBE2R1(C93K, S138C, C191S, C223S)-Ub (EMDB-19567) (PDB 8RX0)* |
|---|---|---|
| ***Data collection*** | | |
| *Microscope* | Glacios | Krios |
| *Detector* | Falcon 4i (counting) | Gatan K3 (counting) |
| *Voltage (kV)* | 200 | 300 |
| *Magnification* | 190,000x | 105,000x |
| *Total electron exposure ($e^{-}$/Å$^2$)* | 26 | 38 |
| *Exposure rate ($e^{-}$/pix/s)* | 7.1 | 15.1 |
| *Defocus range (μm)* | -1.7 to -3.2 μm | -1.2 to -3.0 μm |
| *Pixel size at detector (Å/pixel)* | 0.74 | 0.825 |
| *Total exposure (s)* | 2.0 | 1.8 |
| *Automation software* | EPU | EPU |
| *Energy filter slit width (eV)* | N/A | 20 |
| *Movies collected (#)* | 4,961 | 14,047 |
| *Movies used (#)* | 4,338 | 9,701 |
| ***Reconstruction*** | | |
| *Image processing package* | CryoSPARC | CryoSPARC |
| *Symmetry imposed* | C1 | C1 |
| *Initial particle images (#)* | 405,567 | 748,020 |
| *Final particle images (#)* | 132,697 | 149,823 |
| *Map resolution (Å) at FSC 0.143* | 4.0 | 3.7 |
| *Map resolution range (Å) at FSC 0.5* | 3.0 to 13.0 | 2.5 to 10.0 |
| ***Model composition*** | | |
| *Protein* | Brd4$^{BD2}$, NEDD8, Cul2, Rbx1, EloB, EloC, VHL, UBE2R1(C93K), ubiquitin | Brd4$^{BD2}$, NEDD8, Cul2, Rbx1, EloB, EloC, VHL, UBE2R1(C93K, S138C, S191C, S223C), ubiquitin, ubiquitin (K48C, G76S) |
| *Ligands* | MZ1 | MZ1 |
| *RNA/DNA* | N/A | N/A |
| ***Model refinement*** | | |
| *Initial model(s) used* | 5T35, 5N4W, AlphaFold | 5T35, 5N4W, 4AP4, 6TTU AlphaFold |
| *Model-map cross-correlation* | 0.61 | 0.43 |
| *R.m.s deviations from ideal values:* | | |
| *Bond lengths (Å)* | 0.027 | 0.019 |
| *Bond angles (°)* | 1.930 | 1.661 |
| ***Validation*** | | |
| *MolProbity score* | 1.90 | 2.15 |
| *Clashscore (all atoms)* | 10.66 | 22.08 |
| *Poor rotamers (%)* | 1.27 | 1.09 |
| *Ramachandran plot* | | |
| *Favoured (%)* | 97.01 | 96.03 |
| *Outliers (%)* | 0.58 | 0.46 |
| *CaBLAM outliers (%)* | 1.4 | 1.1 |

**Table S2.** Cryo-EM data collection, image analysis, atomic modeling, refinement, and validation statistics.

### References

1. H. Walden, M.S. Podgorski, B.A. Schulman. Insights into the ubiquitin transfer cascade from the structure of the activating enzyme for NEDD8. *Nature* **422**, 330-334 (2003).
2. H. Walden *et al.*, The structure of the APPBP1-UBA3-NEDD8-ATP complex reveals the basis for selective ubiquitin-like protein activation by an E1. *Mol Cell* **12**, 1427-1437 (2003).
3. S. Diaz, L. Li, K. Wang, X. Liu, Expression and purification of functional recombinant CUL2*RBX1 from E. coli. *Sci Rep* **11**, 11224 (2021).
4. E. Branigan, A. Plechanovova, R. T. Hay, Methods to analyze STUbL activity. *Methods Enzymol* **618**, 257-280 (2019).
5. M. A. Nakasone, N. Livnat-Levanon, M. H. Glickman, R. E. Cohen, D. Fushman, Mixed-linkage ubiquitin chains send mixed messages. *Structure* **21**, 727-740 (2013).
6. M. S. Gadd *et al.*, Structural basis of PROTAC cooperative recognition for selective protein degradation. *Nat Chem Biol* **13**, 514-521 (2017).
7. M. A. Nakasone *et al.*, Structure of UBE2K-Ub/E3/polyUb reveals mechanisms of K48-linked Ub chain extension. *Nat Chem Biol* **18**, 422-431 (2022).
8. M. Walko, E. Hewitt, S. E. Radford, A. J. Wilson, Design and synthesis of cysteine-specific labels for photo-crosslinking studies. *RSC Adv* **9**, 7610-7614 (2019).
9. A. Punjani, J. L. Rubinstein, D. J. Fleet, M. A. Brubaker, cryoSPARC: algorithms for rapid unsupervised cryo-EM structure determination. *Nat Methods* **14**, 290-296 (2017).
10. T. Bepler *et al.*, Positive-unlabeled convolutional neural networks for particle picking in cryo-electron micrographs. *Nat Methods* **16**, 1153-1160 (2019).
11. A. Punjani, H. Zhang, D. J. Fleet, Non-uniform refinement: adaptive regularization improves single-particle cryo-EM reconstruction. *Nat Methods* **17**, 1214-1221 (2020).
12. T. Wagner *et al.*, SPHIRE-crYOLO is a fast and accurate fully automated particle picker for cryo-EM. *Commun Biol* **2**, 218 (2019).
13. E. C. Meng *et al.*, UCSF ChimeraX: Tools for structure building and analysis. *Protein Sci* **32**, e4792 (2023).
14. T. I. Croll, ISOLDE: a physically realistic environment for model building into low-resolution electron-density maps. *Acta Crystallogr D Struct Biol* **74**, 519-530 (2018).
15. G. Young *et al.*, Quantitative mass imaging of single biological macromolecules. *Science* **360**, 423-427 (2018).
16. D. Wu, G. Piszczek, Standard protocol for mass photometry experiments. *Eur Biophys J* **50**, 403-409 (2021).
17. A. Shevchenko, H. Tomas, J. Havlis, J. V. Olsen, M. Mann, In-gel digestion for mass spectrometric characterization of proteins and proteomes. *Nat Protoc* **1**, 2856-2860 (2006).
18. J. Cox, M. Mann, MaxQuant enables high peptide identification rates, individualized p.p.b.-range mass accuracies and proteome-wide protein quantification. *Nat Biotechnol* **26**, 1367-1372 (2008).
19. A. M. Waterhouse, J. B. Procter, D. M. Martin, M. Clamp, G. J. Barton, Jalview Version 2--a multiple sequence alignment editor and analysis workbench. *Bioinformatics* **25**, 1189-1191 (2009).
20. R. C. Edgar, MUSCLE: multiple sequence alignment with high accuracy and high throughput. *Nucleic Acids Res* **32**, 1792-1797 (2004).

21. T. A. F. Cardote, M. S. Gadd, A. Ciulli, Crystal Structure of the Cul2-Rbx1-EloBC-VHL Ubiquitin Ligase Complex. *Structure* **25**, 901-911 e903 (2017).
22. H. Zhou, M. S. Zaher, J. C. Walter, A. A.-O. Brown, Structure of CRL2Lrr1, the E3 ubiquitin ligase that promotes DNA replication termination in vertebrates. *Nucleic Acids Res* **49**, 13194-13206 (2021).
23. K. Baek *et al.*, NEDD8 nucleates a multivalent cullin-RING-UBE2D ubiquitin ligation assembly. *Nature* **578**, 461-466 (2020).
24. H. C. Nguyen, H. Yang, J. L. Fribourgh, L. S. Wolfe, Y. Xiong, Insights into Cullin-RING E3 ubiquitin ligase recruitment: structure of the VHL-EloBC-Cul2 complex. *Structure* **23**, 441-449 (2015).
25. A. Testa, S. J. Hughes, X. Lucas, J. E. Wright, A. Ciulli, Structure-Based Design of a Macrocyclic PROTAC. *Angew Chem Int Ed Engl* **59**, 1727-1734 (2020).
26. A. Hanzl *et al.*, Functional E3 ligase hotspots and resistance mechanisms to small-molecule degraders. *Nat Chem Biol* **19**, 323-333 (2023).
27. T. M. Geiger *et al.*, Discovery of a Potent Proteolysis Targeting Chimera Enables Targeting the Scaffolding Functions of FK506-Binding Protein 51 (FKBP51). *Angew Chem Int Ed Engl* **63**, e202309706 (2024).
28. J. W. Mason *et al.*, DNA-encoded library-enabled discovery of proximity-inducing small molecules. *Nat Chem Biol* **20**, 170-179 (2024).
29. P. S. Dragovich *et al.*, Antibody-Mediated Delivery of Chimeric BRD4 Degraders. Part 2: Improvement of In Vitro Antiproliferation Activity and In Vivo Antitumor Efficacy. *J Med Chem* **64**, 2576-2607 (2021).
30. J. Krieger *et al.*, Systematic Potency and Property Assessment of VHL Ligands and Implications on PROTAC Design. *ChemMedChem* **18**, e202200615 (2023).
31. S. Mathur, P. Deshmukh, S. Tripathi, P. Marimuthu, B. Padmanabhan, Insights into the crystal structure of BRD2-BD2 - phenanthridinone complex and theoretical studies on phenanthridinone analogs. *J Biomol Struct Dyn* **36**, 2342-2360 (2018).
32. R. M. Karim *et al.*, Differential BET Bromodomain Inhibition by Dihydropteridinone and Pyrimidodiazepinone Kinase Inhibitors. *J Med Chem* **64**, 15772-15786 (2021).
33. Z. Yu *et al.*, Discovery and characterization of bromodomain 2-specific inhibitors of BRDT. *Proc Natl Acad Sci U S A* **118**, (2021).
34. A. Punjani, D. J. Fleet, 3D variability analysis: Resolving continuous flexibility and discrete heterogeneity from single particle cryo-EM. *J Struct Biol* **213**, 107702 (2021).
35. G. C. P. van Zundert *et al.*, The HADDOCK2.2 Web Server: User-Friendly Integrative Modeling of Biomolecular Complexes. *J Mol Biol* **428**, 720-725 (2016).
